# Representational learning by optimization of neural manifolds in an olfactory memory network

**DOI:** 10.1101/2024.11.17.623906

**Authors:** Bo Hu, Nesibe Z. Temiz, Chi-Ning Chou, Peter Rupprecht, Claire Meissner-Bernard, Benjamin Titze, SueYeon Chung, Rainer W. Friedrich

## Abstract

Cognitive brain functions rely on experience-dependent internal representations of relevant information. Such representations are organized by attractor dynamics or other mechanisms that constrain population activity onto “neural manifolds”. Quantitative analyses of representational manifolds are complicated by their potentially complex geometry, particularly in the absence of attractor states. Here we trained juvenile and adult zebrafish in an odor discrimination task and measured neuronal population activity to analyze representations of behaviorally relevant odors in telencephalic area pDp, the homolog of piriform cortex. No obvious signatures of attractor dynamics were detected. However, olfactory discrimination training selectively enhanced the separation of neural manifolds representing task-relevant odors from other representations, consistent with predictions of autoassociative network models endowed with precise synaptic balance. Analytical approaches using the framework of *manifold capacity* revealed multiple geometrical modifications of representational manifolds that supported the classification of task-relevant sensory information. Manifold capacity predicted odor discrimination across individuals better than other descriptors of population activity, indicating a close link between manifold geometry and behavior. Hence, pDp and possibly related recurrent networks store information in the geometry of representational manifolds, resulting in joint sensory and semantic maps that may support distributed learning processes.

## INTRODUCTION

Learning generates organized representations of relevant information in the brain that generalize to novel inputs and serve as a basis for cognition. Representational learning is thought to occur by modifications of synaptic connectivity in memory networks that map patterns of input activity to specific subspaces of neuronal state space according to semantic relationships between inputs^1,2^. One way to generate such mappings is through attractor dynamics. Classical models predict that learning enhances recurrent excitation among specific ensembles of neurons, resulting in convergent attractor dynamics that supports the classification of information by pattern completion^3-7^. Moreover, recurrent networks can give rise to a continuum of stable attractor states as observed, for example, in cognitive maps of space^4,8^. Alternatively, representational learning may be mediated by mechanisms that do not rely on attractor dynamics. For example, learning may modify the geometry of neuronal state space and, thus, define *neural manifolds* that represent relevant information without being attractor states^1,2,9-15^. However, as geometrical analyses of neuronal population activity remain challenging, the organization and experience-dependent plasticity of representational neural manifolds in memory networks are still poorly understood. Using large-scale activity measurements in zebrafish and novel mathematical approaches, we examined whether representational memory in a higher olfactory area is consistent with either attractor dynamics or neural manifolds.

Representational memory has been proposed to be a primary function of piriform cortex, a paleocortical brain area that receives distributed sensory input from the olfactory bulb. Within piriform cortex, excitatory neurons are recurrently connected through an “association fiber system” that is plastic and under neuromodulatory control^16-19^. These observations gave rise to the hypothesis that learning results in the formation of neuronal assemblies that represent odor objects and mediate pattern classification by convergent dynamics^20^. However, recent analyses of neuronal activity in piriform cortex did not reveal obvious signatures of attractor states. For example, odor-evoked activity is not persistent but phasic and curtailed by inhibition^21-23^. Moreover, while convergent dynamics are expected to reduce variability, intra-and inter-trial variability of neuronal activity appears high in comparison to inputs from the olfactory bulb^23-26^. Nonetheless, passive odor exposure and active learning modify odor-evoked population activity in piriform cortex depending on stimulus statistics or task structure^25,27-31^. For example, activity patterns evoked by different odors become more highly correlated after presentation of odors in a binary mixture^25^, or after association of multiple odors with a common reward^29^. Hence, experience may reorganize odor representations in piriform cortex in a task-relevant fashion in the absence of discrete attractor dynamics.

Autoassociative memory has recently been explored in computational models of recurrent networks that were constrained by data from telencephalic area pDp of adult zebrafish^32^, the teleost homolog of the piriform cortex^33,34^. When information was stored in these models by enhancing connectivity among small assemblies of excitatory and inhibitory neurons, a transition from discrete attractor dynamics to continuous representational manifolds occurred when networks entered a regime of precise synaptic balance^32^. This regime is characterized by strong and correlated excitatory and inhibitory synaptic currents in individual neurons and thought to be characteristic of cortical circuits^35,36^.

A prerequisite for the analysis of neural manifolds are simultaneous measurements of activity across large neuronal populations. Basic information about the relative separation and orientation of manifolds can then be obtained by different distance measures^32^. Geometrical features of neural manifolds relevant for pattern classification can be further analyzed using “Geometry Linked to Untangling Efficiency” (GLUE) theory^37^, a mathematical framework that extends previous analytical approaches^2,15,38^. GLUE theory quantifies the *untangledness* (“degree of separability”) of manifolds by the measure of *manifold capacity*, which is directly linked to geometrical parameters and neuronal co-variability (see Supplementary Note 1 and ref. ^37^ for details). Hence, manifold capacity quantifies the amount of linearly decodable information per neuron as a function of geometry parameters and correlations. Moreover, manifold capacity can be directly interpreted in the context of computation because it is mathematically linked to the efficiency, capacity and robustness of pattern classification^37^. Recently, GLUE theory has been used successfully to quantify computationally relevant, interpretable features of neuronal population activity in different brain areas and tasks^37^. Manifold capacity analysis can therefore be used to determine how learning modifies geometrical features of representational manifolds, and how these modifications collectively affect the classification of activity representing learned and novel inputs.

To explore how learning modifies representational manifolds we measured population activity in pDp after training juvenile or adult zebrafish in an odor discrimination task. Previous studies showed that odor-evoked activity in pDp is distributed, variable, modified by experience, and under strong inhibitory control^39-44^, consistent with observations in piriform cortex^19-21,23-25,29-31,45,46^. Moreover, in both brain areas, synaptic currents are dominated by recurrent rather than afferent input during an odor response^18,40,42,47^. In pDp, voltage clamp recordings demonstrated that excitatory and inhibitory synaptic currents in individual neurons are strong and co-tuned, providing direct evidence for precise synaptic balance^42^. The analysis of neuronal population activity in pDp therefore allowed us to examine how learning modifies representational manifolds in a precisely balanced autoassociative network.

Consistent with previous observations we found no obvious signatures of global fixed-point attractor dynamics in pDp. Behavioral odor discrimination training had only minor effects on the mean amplitude and variability of odor responses in pDp. Nonetheless, training modified activity manifolds representing learned and related odors, with minimal effects on representations of dissimilar odors. Training modified multiple geometrical features that, in combination, enhanced the separability of manifolds representing task-relevant odors from each other and from manifolds representing novel odors (Extended Data Fig. 1). The separability of activity patterns was correlated with behavioral odor discrimination across trained individuals, indicating that representational manifolds directly contribute to learned behaviors. These results indicate that task-relevant information is stored in pDp by geometrical modifications of the olfactory map, resulting in improved classification of sensory information along relevant directions in coding space. Hence, representational learning in pDp, and possibly in other recurrent balanced-state networks, integrates sensory and semantic information into a continuous map, which may support distributed learning processes.

## RESULTS

### Basic features of neural dynamics in pDp

We measured odor-evoked activity in an ex-vivo preparation of the juvenile and adult zebrafish brain by volumetric 2-photon calcium imaging^48,49^ in transgenic fish expressing the calcium indicator GCaMP6s (Tg[alphaTubulin:GCaMP6s]) throughout the telencephalon and other brain areas^50^ (Fig. 1a,b). 3D scanning was performed by remote focusing with a volume rate of 7.5 Hz and calcium signals (ΔF/F) were transformed into firing rate estimates using CASCADE^43^. Activity across populations of neurons was represented by n-dimensional activity vectors, each representing the estimated firing rates of n neurons within a time bin. Most experiments were performed in juvenile fish (46 – 56 days post fertilization; 9.0 – 13.5 mm body length), which allowed us to measure activity across the majority of neurons in pDp. Consistent with previous results from adult zebrafish^39-42^, odor-evoked activity in juvenile pDp exhibited an initial phasic increase followed by a slowly decaying plateau (Fig. 1c,d; Extended Data Fig. 2a), and responses of individual neurons were scattered throughout pDp without an obvious odor-related topography^39^.

**Fig. 1.**
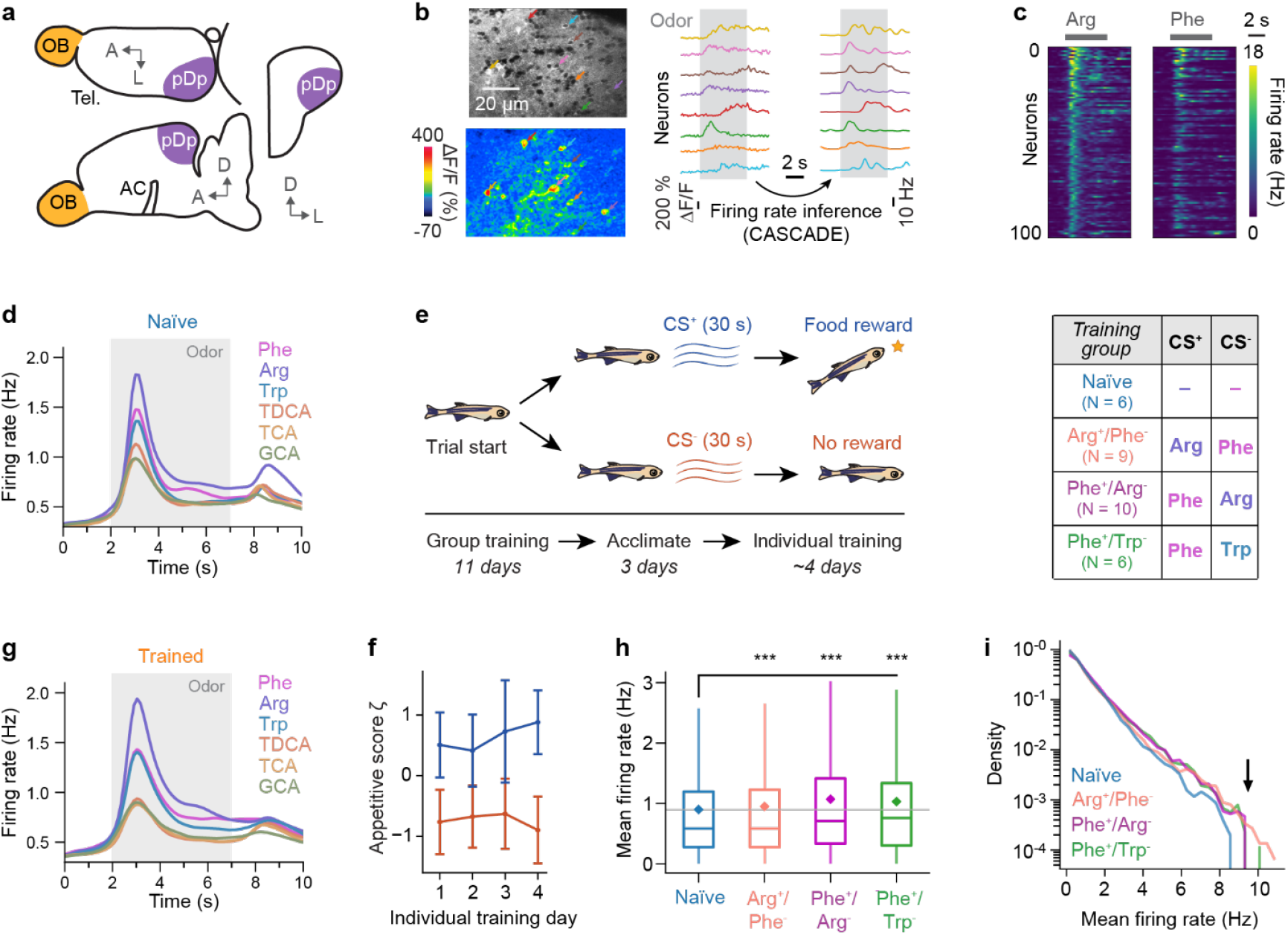
Dynamics of odor-evoked activity in pDp and discrimination training of juvenile zebrafish. **a**, Location of pDp in the telencephalon (Tel.) of juvenile zebrafish. OB: olfactory bulb; AC: anterior commissure; A: anterior; L: lateral; D: dorsal. **b**, Left: αTubulin:GCaMP6s expression in juvenile pDp (2-photon optical section; subregion of entire field of view; top) and relative change in fluorescence (ΔF/F) evoked by odor stimulation (Arg) in the same field of view, averaged over 3 s (bottom). Right: ΔF/F traces and inferred firing rates of neurons depicted by arrows with matching colors. Gray shading indicates odor application. **c**, Inferred firing rates of 100 randomly selected pDp neurons showing responses to Arg and Phe, sorted by mean response intensity to Arg in both panels. **d**, Inferred firing rate as a function of time, averaged over all neurons (n = 9059) and fish (n = 6; naïve juvenile group). **e**, Odor discrimination task. Juvenile fish were trained as a group in a home tank (11 days) and further trained individually for 2 - 4 days after acclimatization to single-tank housing. Behavior was quantified during individual training. Right: training groups. **f**, Appetitive scores (ζ; mean ± SD) for CS^+^ and CS^-^ odors on each day of individual training from all trained fish. Appetitive scores were significantly higher for CS^+^ than CS^-^ on all days (day 1: n = 25, P = 6.0 × 10^-8^; day 2: n = 25, P = 1.8 × 10^-7^; day 3: n = 21, P = 6.7 × 10^-6^; day 4: n = 14, P = 1.2 × 10^-4^; Wilcoxon signed-rank test). **g**, Mean inferred firing rate evoked by each odor, averaged across all neurons and trials from trained fish (N = 25 fish, n = 34152 neurons; all training groups combined). **h**, Mean activity evoked by amino acid odors in different training groups. Significance levels for statistical comparisons (Mann-Whitney U test) in all panels: ns, P ≥ 0.05; ***, P < 0.001; see Supplementary Note 2 for details. **i**, Distribution of firing rates evoked by amino acid odors in different training groups. Density is normalized by the area under each curve. High firing rates were more frequently observed in trained fish (arrow).

Classical models of autoassociative memory assume that pattern classification is based on convergent intrinsic attractor dynamics, which can map noisy variations of the same input onto a common, less variable output^3-5^. In olfaction, considerable variability of odor-evoked activity arises, for example, from varying background odors, fluctuations in odor concentration, neuronal adaptation, variations in odor sampling behavior, variations in brain state, and neural noise^19,20^. Convergent intrinsic attractor dynamics may therefore facilitate the classification of different activity patterns representing a common odor. Alternatively, classification may be supported by mapping semantically related inputs into a common neural subspace (manifold) in the absence of attractor dynamics. In this scenario, activity within the subspace may itself be variable but stimulus information could nonetheless be retrieved efficiently by the selective readout of activity from the manifold^32,51,52^. Hence, neural manifolds may support classification and learning by organizing representations according to semantic variables and facilitating access to relevant information. We examined whether odor-evoked activity in pDp is consistent with either convergent intrinsic attractor dynamics or with representational manifolds.

Autoassociative circuits can exhibit intrinsic attractor states that persist after cessation of sensory input^4^. Previous results from pDp and piriform cortex, however, failed to provide strong evidence for such attractor dynamics in response to novel or learned odors^22-26,30,39-42,45^. We further examined signatures of attractor states by testing two predictions (Extended Data Fig. 3a): (1) as a consequence of convergent dynamics, trial-to-trial variability is expected to be lower in output activity patterns than in input patterns, and (2) because attractor states are stable, activity is expected to persist for some time when the input is switched off abruptly. To test the first prediction, we measured odor-evoked activity in pDp and in the olfactory bulb of the same juvenile zebrafish. Consistent with previous observations^39,40,53-55^, trial-to-trial variability in pDp was substantially higher than in the olfactory bulb (Extended Data Fig. 3b). Moreover, variability increased, rather than decreased, during odor presentation (Extended Data Fig. 3c). These observations do not support the hypothesis that variability of activity in pDp is reduced by convergence onto a distinct attractor state.

To test the second prediction we examined changes in correlations between odor-evoked activity patterns as a function of time. After termination of odor application, mean firing rates showed a rebound and decayed slowly thereafter but activity across the population changed rapidly, as revealed by a sudden decrease in pattern correlations (Extended Data Fig. 3d,e). Hence, global activity patterns did not persist, which can be attributed to the classical off-response. Nonetheless, prolonged activity was observed in odor-specific subsets of neurons (Extended Data Fig. 4). This observation may indicate that subpopulations of neurons, within pDp or elsewhere, engage in attractor states that outlast the stimulus. Alternatively, slowly decaying activity may reflect the gradual decline in stimulus concentration after termination of the odor supply (Extended Data Fig. 4).

To further address the second prediction in the absence of complex dynamics after odor offset we abruptly silenced sensory input from the olfactory bulb by optogenetic stimulation of inhibitory interneurons during an odor response (Extended Data Fig. 3f). We then analyzed activity in pDp using whole-cell current and voltage clamp recordings. Upon silencing, membrane potentials and synaptic currents decayed nearly exponentially with time constants (membrane potential: 65 ± 23 ms; currents: 60 ± 21 ms) similar to the membrane time constant of pDp neurons (52 ± 16 ms; Extended Data Fig. 3f-h). A fast decay of activity after optogenetic silencing at different time points was also observed using 2-photon calcium imaging (Extended Data Fig. 3i-k). Together, these and additional results (see below) indicate that odor-evoked activity in pDp does not exhibit global fixed-point attractor dynamics (although attractor dynamics in neuronal subpopulations within or outside pDp cannot be excluded). We therefore further explored the hypothesis that odors are represented by experience-dependent neural manifolds in pDp.

### Experience-dependent modification of neural dynamics in pDp

To examine how odor representations in pDp are modified by experience we trained juvenile zebrafish expressing GCaMP6s^56^ in the telencephalon in an odor discrimination task using procedures developed for adult fish^57^ with minor modifications. Tanks were continuously perfused with fish water and two amino acid odors (CS^+^, CS^-^) were each added to the perfusion 9 times per day for 30 s. Presentation of the CS^+^, but not the CS^-^, was followed by food delivery at a specific location. Fish were initially pre-trained as a group for 11 days before individual fish were further trained for 2 to 4 days in separate tanks (Methods; Fig. 1e). Three groups of fish were trained on different CS^+^/CS^-^ assignments (Fig. 1e, right). Learning was monitored by quantifying appetitive behavior of individual fish during odor presentation as described^57^. After training, appetitive behavior was significantly more pronounced in response to the CS^+^ than to the CS^-^ (Fig. 1f), as observed in adult fish^57,58^.

We measured activity in pDp of trained and naïve fish in response to three amino acids (Phe, Arg, Trp) and three bile acids (TDCA, TCA, GCA; 5 s; three trials each), which represent two classes of natural odors for aquatic animals^59^. In the olfactory bulb, activity patterns evoked by these odorants are modestly correlated within each chemical class but uncorrelated across classes^39^. The time course of odor responses was similar in trained and naïve fish although tonic components appeared somewhat more pronounced after training (Fig. 1g; Extended Data Fig. 2a, 3d,e). Mean responses to amino acids were slightly but significantly larger in each of the trained groups (Fig. 1h), which could be attributed, in part, to large odor responses in a small subset of individual neurons (Fig. 1i).

The cosine distance between time-averaged activity patterns evoked by the same odor in different trials was only slightly lower than the cosine distance between activity patterns evoked by different odors from the same category (Fig. 2a). In trained fish, cosine distances between activity patterns evoked by the CS^+^ and the CS^-^ were similar to the corresponding distances in naïve fish while other cosine distances were slightly higher (Fig. 2a-c; Extended Data Fig. 3b). In particular, cosine distances between representations of dissimilar odors (amino acids versus bile acids) were increased, resulting in a broader overall distribution of cosine distances (p = 0.04; F-test; Fig. 2a, Extended Data Fig. 2b). No major changes were observed in the dimensionality of activity patterns (Extended Data Fig. 2c). Hence, training did not substantially reduce trial-to-trial variability but slightly modified the similarity between odor-evoked activity patterns.

**Fig. 2.**
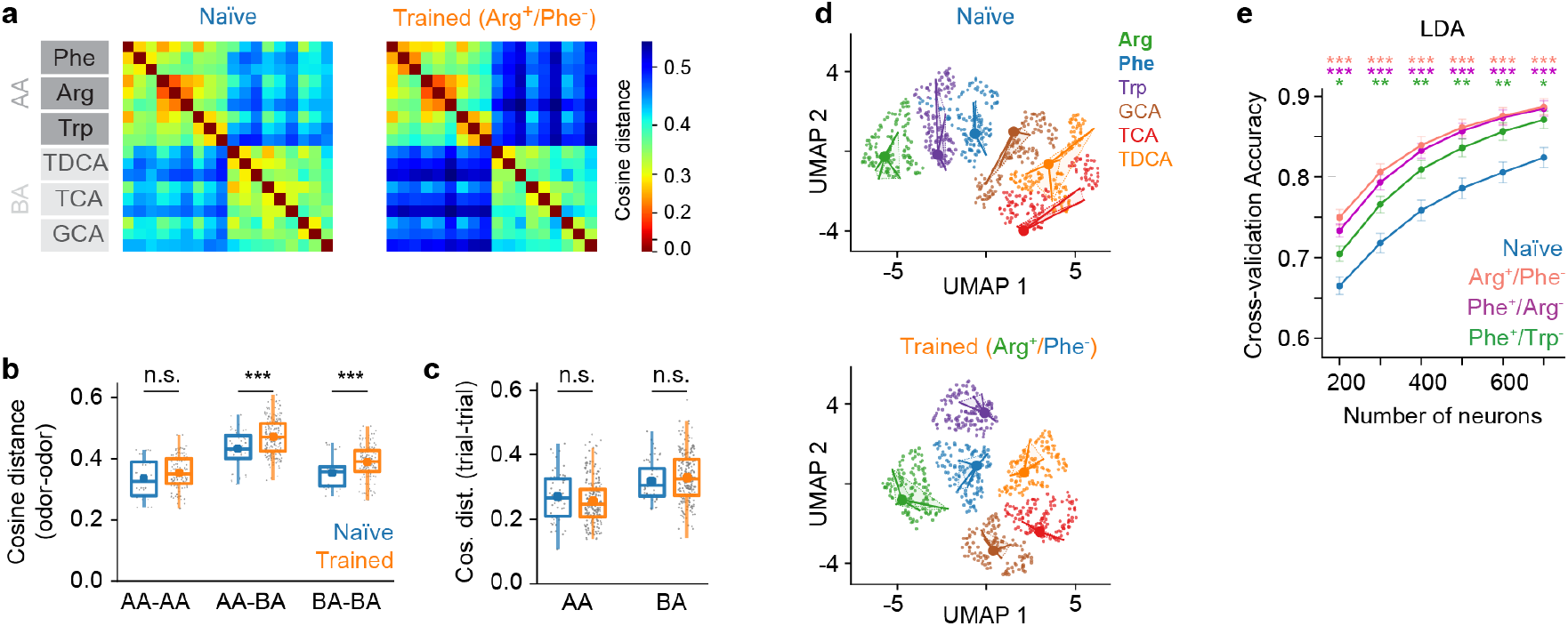
Effects of odor discrimination training on population activity in pDp. **a**, Mean cosine distance between activity patterns evoked by different odors (3 trials each) in naïve fish (left) and in fish trained on Arg (CS^+^) vs. Phe (CS^-^). **b**, Cosine distances between trial-averaged activity vectors representing different odor categories in naïve and trained fish. **c**, Cosine distance between responses to the same odors (trial-trial variability), analyzed separately for amino acids (AA) and bile acids (BA) in naïve and trained fish. **d**, Projections of activity patterns evoked by different odors (colors) onto the first two UMAP components in naïve (N = 6) and Arg^+^/Phe^-^-trained fish (N = 9). Small symbols depict activity in individual trials at different time points (1 s – 6 s after response onset). Filled circles and lines represent the mean ± SD along the directions of the first two principal components in original activity space. **e**, Cross-validation accuracy of pairwise odor classification by linear discriminant analysis (LDA) as a function of neuron number (subsampling) across training groups (mean ± SEM; n = 50 repeats). Color-coded asterisks show statistical significance for comparison to Naïve group (*, P < 0.05; **, P < 0.01; ***, P < 0.001).

While high variability is atypical for fixed-point attractors it does not exclude chaotic attractor states. A signature of chaotic attractor states are persistent pattern correlations, even when variability is high^6^. However, correlations between activity patterns decreased rapidly after stimulus offset in trained as well as in naïve animals (Extended Data Fig. 3d,e). Hence, our observations do not support the hypothesis that training establishes convergent attractor dynamics representing specific odors.

To further explore effects of training we visualized odor representations by projecting population activity onto the first two UMAP components. In trained fish, activity patterns representing different odors appeared more distinct than in naïve fish (Fig. 2d). Consistent with this observation, the accuracy of odor classification by linear discriminant analysis (LDA) was consistently higher in trained fish (Fig. 2e). Similar results were obtained using a support vector machine for classification (Extended Data Fig. 2d). Hence, training (passive odor exposure or active learning) enhanced odor classification in pDp without establishing obvious attractor dynamics.

### Basic analysis of manifold geometry in a computational model

To examine representational learning in more detail we analyzed odor-evoked population activity in pDp as neural manifolds. We first used two distance measures, Euclidean distance (dE) and Mahalanobis distance (dM), to characterize basic geometrical relationships between neural manifolds in activity space (the state space where each dimension represents activity of one neuron). dE was defined as the distance between the centers of two distributions (mean of data points representing a manifold). Hence, dE quantifies the separation of prototypical odor representations in state space. dM, in contrast, also depends on the shape of distributions: it measures the distance between a point (individual activity vector) and a reference distribution (neural manifold) in units that depend on the shape and extent (covariance pattern) of the reference distribution. dM between two manifolds is thus computed by averaging over the dM between each datapoint in one manifold and the distribution representing the other manifold. As manifolds typically have different shapes, dM in the direction from manifold A to manifold B (dM_A→B_) is usually different from dM in the opposite direction (dM_B→A_), providing information about manifold geometry (Fig. 3a,b). For example, dM_A→B_ > dM_B→A_ implies that A is more elongated than B along the A-B axis. Because dM is normalized by covariance it is closely related to the separability of two distributions.

**Fig. 3.**
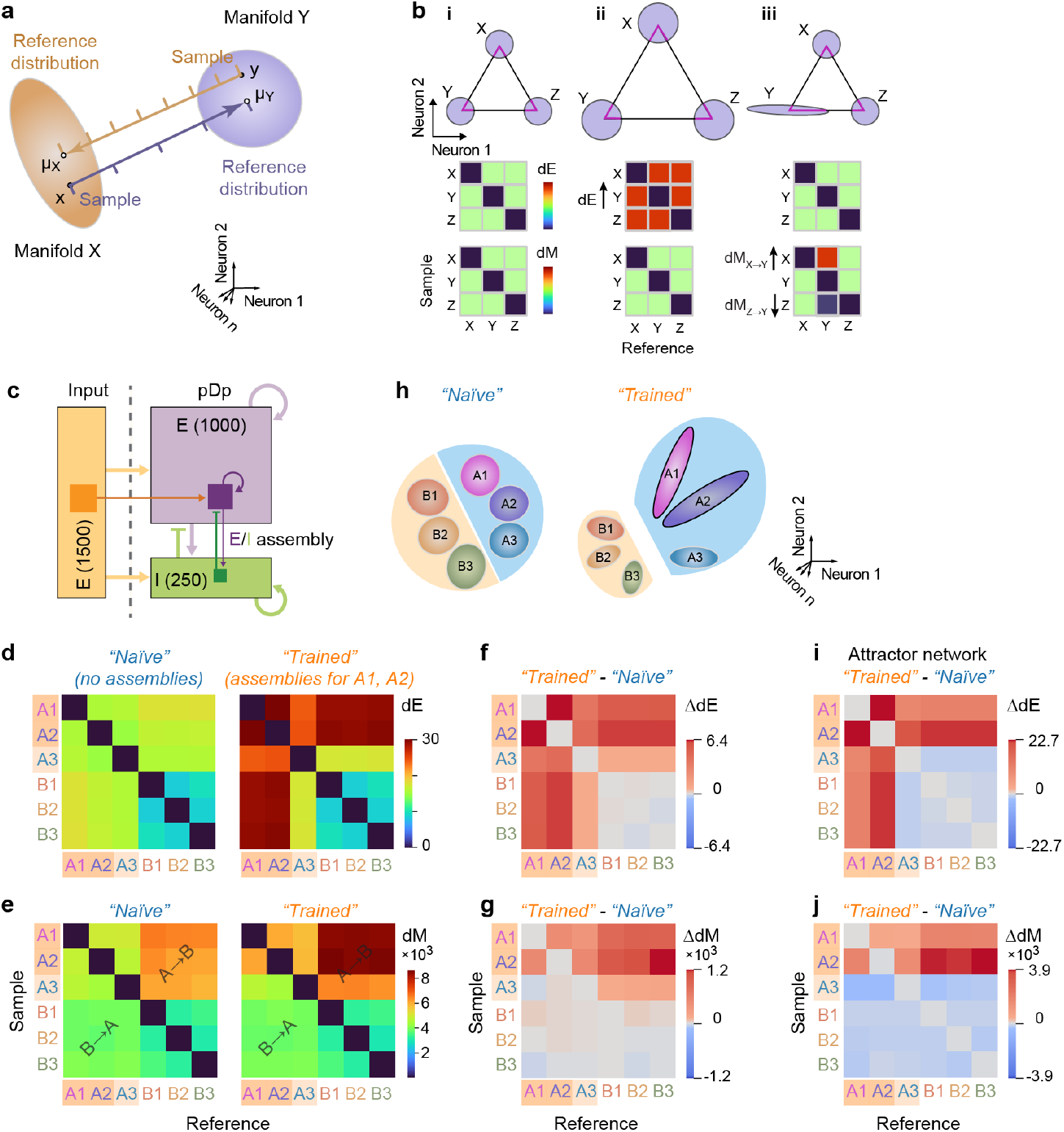
Analysis of distances between neural manifolds: computational model. **a**, Mahalanobis distance (dM): schematic illustration. Yellow and purple ellipsoids depict neural manifolds (sets of activity vectors evoked by an odor, pooled over trials and time points). dM from a sample point x in manifold X to the reference distribution computed from manifold Y is the distance from x to the centroid of Y (μ_Y_), relative to the directional variability in Y (ruler). dM_X→Y_ is the mean dM from each point in X to manifold Y. Note that dM is usually asymmetric because manifolds are usually different. **b**, Example sets of three manifolds (X, Y, Z) illustrating effects of linear scaling (ii) and geometrical modifications (iii) on dE (top) and dM (bottom). **c**, Spiking network model. 1000 recurrently connected excitatory pDp neurons (E; purple) and 250 recurrently connected inhibitory neurons (I; green) received input from 1500 excitatory mitral cells (yellow). Information about defined odors (“memory”) was stored in the recurrent connectivity matrix (“learning”) by increasing the connection probabilities among strongly activated E and I neurons (“E/I assembly”). **d**, Matrix showing Euclidean distances dE between manifolds representing six input patterns (odors) in a randomly connected network without E/I assemblies (left, “Naïve”) and after introducing E/I assemblies representing odors A1 and A2 (right; “Trained”). Odors were separated into two classes (A and B) by their correlations. **e**, Equivalent matrices for dM. **f**, Difference between dE matrices in **d**. Note that E/I assemblies (“learning”) increased dE between learned and other odors, and that this effect generalized to a related odor (A3). **g**, Difference between dM matrices in **f**. Note that dM increased selectively in the direction from learned to other odors (dM_A1→other_; dM_A2→other_), and this effect generalized to the related odor (dM_A3→other_). **h**, Simplified illustration of learning-related changes in manifold geometry: E/I assemblies for learned odors (A1, A2) result in a directional displacement and extension of the corresponding manifolds that can be attributed, at least in part, to a modest amplification of activity within the assembly. This effect partially generalizes related representations (A3). Note that the arrangement of manifolds in a two-dimensional plane is a simplification to facilitate visualization; the true dimensionality is higher (Extended Data Fig. 2c). **i**,**j**, Difference matrices showing changes in dE and dM, respectively, after introducing assemblies representing A1 and A2 into a closely related network with attractor dynamics. Note that increases in dE and dM do not generalize to the related third odor (A3).

We define a manifold representing a given odor as the distribution of all activity vectors (“point clouds”) measured in response to the odor, pooled over time points and trials. By analyzing the separability of point cloud manifolds we therefore ask how well activity evoked by one odor at different time points and in different trials can be distinguished from activity evoked by other odors at different time points and in different trials. dE is symmetric and depends both on the angular separation of manifold centers and their amplitudes (firing rates). dM was determined between individual activity vectors x (activity evoked by an odor in an individual trial and time bin) and activity distributions Y (the set of activity vectors evoked by another odor in multiple trials and time bins). dM is defined as

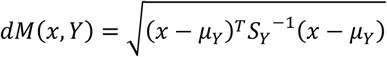

where μ_Y_ and S_Y_ are the mean and the covariance matrix, respectively, of Y. Hence, dM is the effective distance between x and the mean of Y, taking into account the variability in Y. dM between manifolds representing odors X and Y in the direction X→Y (dM_X→Y_) was calculated by averaging over dM between each vector x in X and manifold Y. The same procedure was used to determine dM in the opposite direction (dM_Y→X_). Because dM_X→Y_ is usually not equal to dM_Y→X_ matrices quantifying dM between multiple manifolds contain information about their separation and geometry along selected directions (Fig. 3a,b).

We then jointly quantified dE and dM for manifolds representing different odors. As illustrated in Fig. 3bi, all dE and dM are equal when manifolds are equidistant and spherical with equal variance. When manifolds are scaled uniformly, dE changes while dM remains constant because Euclidean distance and variance co-scale (Fig. 3bii). When only the geometry of one manifold is modified, dE is unaffected but dM changes between specific subsets of manifolds (Fig. 3biii). Matrices of dE and dM therefore contain basic information about distances between representational manifolds and about their geometry in a subset of state space dimensions.

To predict effects of learning on dE and dM we simulated odor-evoked activity using recurrently connected, precisely balanced networks of spiking neurons (Fig. 3c). Previous models of adult pDp^32,44^ consisted of 4000 excitatory and 1000 inhibitory neurons. Learning was assumed to enhance connectivity between small assemblies of strongly activated neurons (100 excitatory and 25 inhibitory neurons), which resulted in the separation of manifolds representing learned and related odors from other manifolds. To adapt these computational models to juvenile fish we reduced network size by a factor of four while maintaining precise synaptic balance. We then simulated responses to six input patterns with response amplitudes and correlations matching the correlations between activity patterns evoked by the experimental panel of amino acid and bile acid odors across mitral cells of the olfactory bulb (Methods). Virtual odors were thus separated into inputs corresponding to amino acids (odors A1, A2, A3) or bile acids (B1, B2, B3) based on their correlations. In networks without assemblies, dE was higher between representations of A-odors than between representations of B-odors, and highest across odor classes (Fig. 3d, left). dM was also higher between A-odor than between B-odor representations. Across odor classes, however, dM was higher in the direction from vectors representing A-odors to reference distributions (manifolds) representing B-odors (dM_A→B_) than in the opposite direction (dM_B→A_; Fig. 3e, left).

Introducing assemblies representing two A-odors increased dE between manifolds representing these “learned” and other odors (Fig. 3d, f), which can be attributed to a modest increase in firing rates of neurons within but not outside an assembly^32^. This effect generalized partially to the third A-odor, but not to B-odors. dM was increased for learned odors specifically in the direction dM_A→B_ (from A-odors to B-odors), indicating that manifold geometry changed non-uniformly (Fig. 3e,g). This asymmetric change in dM also generalized to the third A-odor, while no obvious effects were observed on distances between B-odors. As summarized in Fig. 3h, the introduction of autoassociative memories therefore resulted in specific geometrical modifications of manifolds representing learned and related odors, consistent with previous observations in the larger computational model^32^.

We further observed that assemblies increased cosine distances between responses to dissimilar inputs (A versus B odors) while distances between activity patterns evoked by repeated applications of the same odor were slightly decreased (Extended Data Fig. 5). As a consequence, the distribution of cosine distances was significantly broader than in randomly connected networks (p = 0.02; F-test). This broadening can be attributed to systematic effects of assemblies (Extended Data Fig. 5) and resembled the changes in cosine distances observed in fish after odor discrimination training (Fig. 2g).

We next studied odor representations in simulated networks that included only excitatory neurons in assemblies while inhibition was increased globally to prevent runaway activity. As described previously^32^, this modification disrupted precise balance and resulted in convergent attractor dynamics in response to “learned” odors. Because assembly neurons fired at high rates in the attractor state^32^, dE between representations of learned and other odors was strongly increased and dM was increased asymmetrically from learned to other odors (dM_A1,A2→other_; Fig. 3i,j). Unlike in precisely balanced networks, however, these effects did not generalize to the related odor (A3) because attractor states were discrete. Hence, the generalization of learning-related effects across related odors distinguishes gradual modifications of representational manifolds from discrete attractor states.

### Learning-related modifications of representational manifolds in pDp

To determine how discrimination training modified representational manifolds in pDp we first compared matrices of dE and dM in naïve and trained fish (Fig. 4). dE was increased between the CS^+^, the CS^-^, and the third amino acids (AA3), and between amino acids and bile acids, whereas dE between bile acids was not significantly different. This pattern, including the generalization of effects from conditioned to related odors, was observed in all training groups. Matrices of dM were asymmetric, with higher dM in the direction from amino acids to bile acids (dM_AA→BA_) than in the opposite direction (dM_BA→AA_; Fig. 4b). In all groups of trained fish, dM was increased asymmetrically between amino acid representations (CS^+^, CS^-^ and AA3; AA-AA) and between amino acids and bile acids (AA-BA) but not between bile acid representations (BA-BA; Fig. 4d,f). Largest increases were observed between representations of amino acids and bile acids (AA-BA) in the direction from amino acid activity vectors to bile acid reference distributions (dM_AA→BA_), whereas dM in the opposite direction (dM_BA→AA_) remained almost constant. Shuffling of odor labels decreased all dE and dM and abolished significant differences between naïve and trained groups (Extended Data Fig. 6). Hence, effects of discrimination training on representational manifolds were consistent with predictions derived from precisely balanced network models but not from attractor models.

**Fig. 4.**
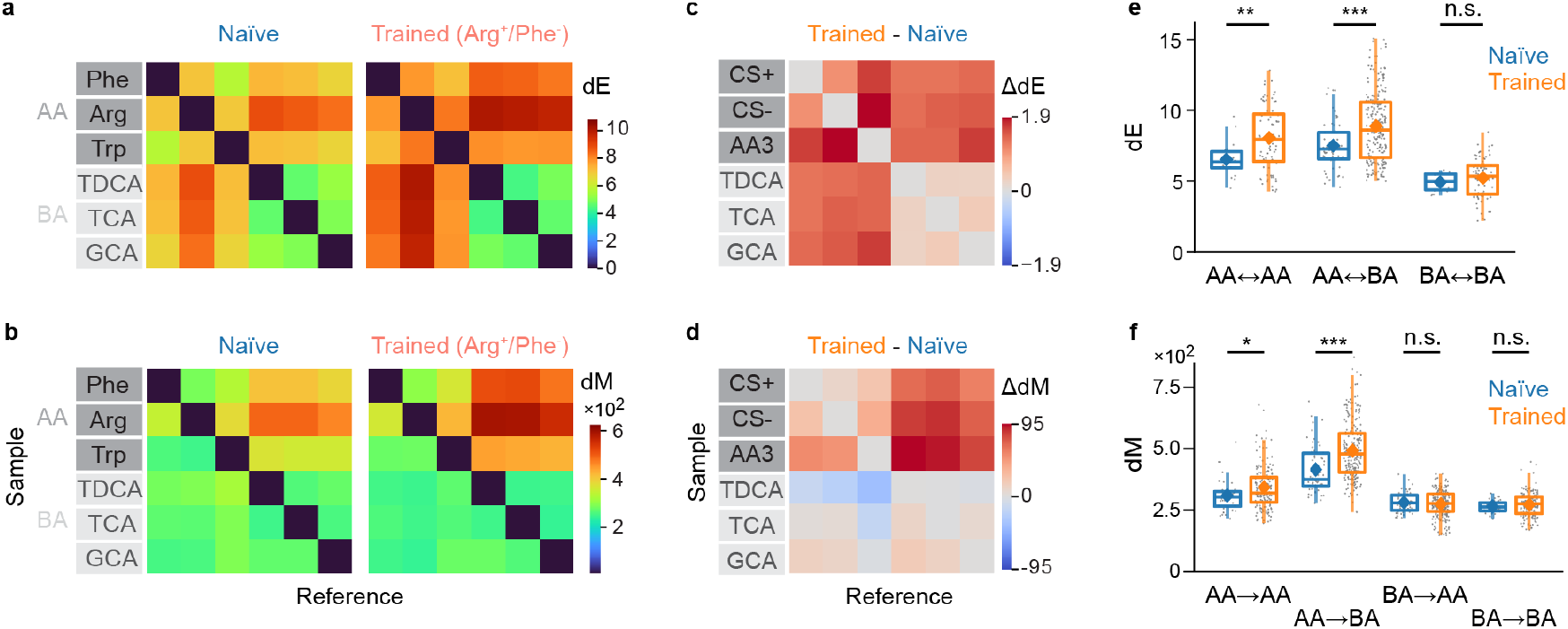
Learning-related changes of representational manifolds in pDp of juvenile zebrafish. **a**,**b**, Pairwise dE and dM between manifolds representing amino acid (AA) and bile acid (BA) odors in naïve juvenile fish (left; N = 6 fish) and in Arg^+^/Phe^-^-trained fish (N = 9). **c**,**d**, Difference in dE and dM between trained (all training groups; N = 25) and naïve fish (N = 6). AA odors were reordered by reward assignment (CS^+^, CS^-^, third amino acid) prior to averaging of matrices over training groups. **e**, dE between pairs of manifolds as a function of odor categories. Significance levels (Mann-Whitney U test) in all panels: ns, P ≥ 0.05; *, P < 0.05; **, P < 0.01; ***, P < 0.001; see Supplementary Note 2 for more details. **f**, Pairwise dM as a function of odor categories. Conventions as in **e**.

These observations indicate that training had two major effects on representational manifolds of conditioned and related odors: first, manifold centers moved away from each other and from other manifolds, as revealed by the increase in dE. Second, manifold geometry changed in a non-uniform, odor-specific manner. The finding that training increased dM_AA→BA_ but not dM_BA→AA_ implies that representational manifolds for amino acids expanded in the direction from amino acids to bile acids because the increase in dE must be accompanied by an increase in variability of similar magnitude in the same direction. In other directions, however, dM increased, indicating that variability increased less than dE. As dM directly reflects the linear separability of a given pattern from a reference distribution, these results directly imply that learning enhanced the linear discriminability of activity vectors representing learned or related odors from manifolds representing other odors. This result is consistent with the increase in odor classification accuracy after training observed by LDA (Fig. 2e; see also Extended Data Fig. 2d) and provides insights into the underlying geometrical modifications.

### Changes in manifold geometry support pattern classification

To further examine how learning-related changes in manifold geometry affect pattern classification we used GLUE theory^37^. This analytical framework quantifies the untangledness (“separability”) of manifolds using manifold capacity^2,15,38^, a metric that assesses how efficiently manifolds are stored in neural state space for retrieval via linear readout. GLUE theory uses a set of parameters to describe geometrical features and correlations of manifolds and analytically links these parameters to classification error. Hence, GLUE theory provides a mathematical framework to understand how the separability of manifolds depends on their geometry and correlations^37^.

Manifold capacity reflects the number of randomly chosen neurons (state space dimensions) required to classify a given set of manifolds (Fig. 5a). Higher manifold capacity indicates greater discriminability, i.e., a downstream neuron, modeled as a linear sum followed by thresholding, can distinguish between manifolds by accessing a smaller number of upstream neurons. Manifold capacity is not based on Gaussian approximations. Moreover, unlike methods such as linear discriminant analysis or support vector machines, manifold capacity does not quantify the performance of an optimal decoder with access to all neurons. Rather, manifold capacity measures the “suitability” of representational manifolds to learn pattern classification tasks. This measure reflects “untangledness” and is analytically linked to computationally relevant properties of representations such as their efficiency, capacity and robustness^37^. For a more detailed account of GLUE theory including a brief review of the underlying mathematics see Supplementary Note 1 and ref ^37^.

**Fig. 5.**
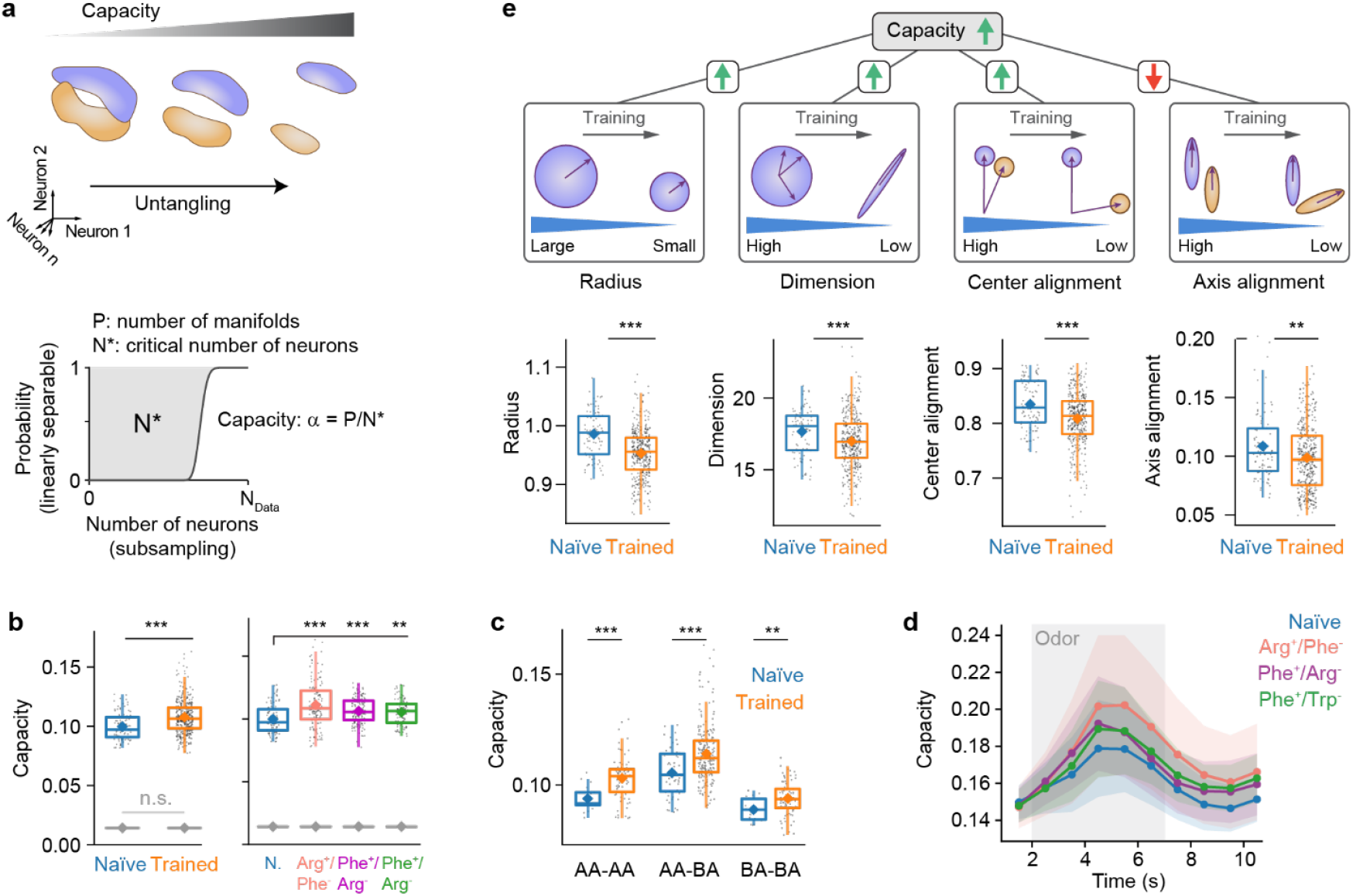
Manifold capacity analysis. **a**, Top: schematic illustration of untangling by geometrical modifications of manifolds. Bottom: The manifold capacity α is determined by the critical number of neurons N* required to classify P manifolds with given geometrical and correlational properties. See Supplementary Note 1 and ref. ^37^ for details. **b**, Manifold capacity in pDp of naïve (N = 6) and trained juvenile fish (N = 25). Left: all trained groups combined (gray: after shuffling of labels). Right: separated by training group. For all panels: ns, P ≥ 0.05; *, P < 0.05; **, P < 0.01; ***, P < 0.001; see Supplementary Note 2 for more detailed statistical information. **c**, Manifold capacity in naïve and trained fish, separated by odor class, pooled over all training groups. **d**, Manifold capacity as a function of time in different training groups (3 s time window, centered on the specified time points). Shading depicts odor application. **e**, Effective geometric measures contributing to manifold capacity with schematic summary. Effective radius measures how far the manifold extends from its center. A decrease indicates a more compact manifold. Effective dimension measures the number of directions in which the manifold extends. A decrease indicates a “flatter” manifold. Effective alignment measures are defined for pairs of manifolds. Effective center alignment measures the correlation between manifold centers. Effective axis alignment measures the amount of directional similarity between manifolds. A decrease indicates that manifolds extend in more different directions. Note that these geometric measures are effective measures computed after reweighting manifold points based on their importance in classification, and thus can be different from the intrinsic geometric measures of the manifolds alone. Arrows depict how the observed effects of training on each geometric measure affected manifold capacity: decreases in effective radius, dimension and center alignment increased capacity while a decrease in effective axis alignment decreased capacity.

Standard GLUE theory expresses manifold capacity as a function of four parameters that measure interpretable geometrical features of manifolds and their correlations^37^: (1) effective manifold radius (related to “compactness”), (2) effective dimension (related to “flatness”), (3) effective center alignment (related to “pattern correlation”, i.e., the correlation between activity vectors representing manifold centers), and (4) effective axis alignment (related to the “relative orientation” of manifolds). These measures provide a statistical description of representational manifolds that is linked to manifold capacity through a mathematical characterization of how linear readout interacts with manifold structure. The relationship between manifold capacity and the four parameters is monotonic: manifold capacity increases when effective radius decreases, effective dimension decreases, effective center alignment decreases and effective axis alignment increases.

We quantified manifold capacity based on sets of activity vectors within a 7 s time window starting at response onset, pooled over the three trials for each odor. Across all odors and training groups, manifold capacity was significantly higher in trained (0.108 ± 0.014; n = 375 odor pairs from N = 25 fish; mean ± SD) than in naïve fish (0.100 ± 0.011; P = 2.8 × 10^-7^; n = 90 odor pairs from N = 6 fish; Fig. 5b-d; Extended Data Fig. 7c). Shuffling manifold labels across all activity patterns substantially decreased the resulting capacity and abolished differences between naïve and trained groups (Naïve: 0.014 ± 0.0001; Trained: 0.014 ± 0.0001; P = 0.85). To examine whether changes in manifold capacity were specific to an odor category we separately analyzed different classes of odor pairs (AA-AA, AA-BA, BA-BA). For all three classes, manifold capacity was significantly higher in trained than in naïve fish (AA-AA: 7.4 × 10^-5^; AA-BA: P = 1.7 × 10^-5^; BA-BA: P = 0.001; Fig. 5c). Largest relative differences in manifold capacity were observed for AA-AA pairs (9.6 ± 12.7 %), followed by AA-BA (8.6 ± 16.8 %) and BA-BA pairs (5.6 ± 10.8 %). As expected, manifold capacity was almost independent of the number of subsampled neurons (Extended Data Fig. 7c). We therefore conclude that training resulted in geometrical modifications of representational manifolds that support odor classification.

To further understand these geometrical modifications we analyzed the contributions of different geometrical parameters (Fig. 5e; Extended Data Fig. 7). In trained fish, the effective radius, dimensionality, center alignment and axis alignment were all significantly lower than in naïve fish (effective radius: 0.99 ± 0.04 [Naïve] versus 0.95 ± 0.04 [Trained]; P = 4.0 × 10^-10^; effective dimensionality: 17.7 ± 1.6 versus 17.0 ± 1.9; P = 0.0007; effective center alignment: 0.84 ± 0.03 versus 0.81 ± 0.05; P = 5.0 × 10^-5^; effective axis alignment: 0.109 ± 0.028 versus 0.099 ± 0.028; P = 0.006). The increase in manifold capacity was therefore due to decreases in effective manifold radius, dimensionality, and center alignment that outweighed opposite effects of effective axis alignment on manifold capacity (Fig. 5e).

Analyses using shorter time windows (3 s) revealed that manifold capacity increased during odor presentation and decreased again after odor offset (Fig. 5d). The underlying geometrical and alignment measures followed a similar timecourse (Extended Data Fig. 7d,e). Hence, odor representations became more suitable for classification over time even though mean firing rates decreased (Fig. 1d,g) and variability increased (Extended Data Fig. 3c). This effect was more pronounced in trained fish (Fig. 5d; Extended Data Fig. 7d,e).

In summary, training increased manifold capacity, which can be attributed primarily to decreases in effective manifold radius, dimensionality and center alignment. Similar changes in manifold capacity and the underlying effective geometrical and correlational parameters were observed in computational models after introducing assemblies (Extended Data Fig. 8). A decrease in effective radius indicates that manifolds become more “compact”, thus enhancing their separability, while a decrease in effective dimensionality implies that manifolds become more constrained in specific state space dimensions (more “flat”). A decrease in effective center alignment supports pattern classification by decorrelating representations. Our results indicate that these changes contribute most prominently to the untangling of task-relevant representational manifolds after training.

### Experience-dependent modification of representational manifolds in adult fish

We next examined odor representations in adult zebrafish, where pDp is substantially larger. Effects of training may thus be expected to generalize less from conditioned stimuli to related stimuli because a larger network allows for more specific local modifications of manifold geometry. Adult zebrafish were trained^57^ using Trp as CS^+^ and alanine (Ala) as CS^-^. After training, activity patterns were measured in response to Trp, Ala, and two additional amino acids, serine (Ser) and histidine (His), that are structurally similar to the conditioned odors and evoke correlated responses in the adult olfactory bulb^53,60^. As observed in juvenile fish, training slightly increased odor responses, strong responses were observed more frequently, and trial-to-trial similarity between odor responses was low in comparison to the olfactory bulb^41,53-55^ (Extended Data Fig. 9a-c). Moreover, the distribution of cosine distances between odor-evoked activity patterns was significantly broader in trained fish (p < 0.01, F-test; Extended Data Fig. 9d), as observed in juvenile fish (Fig. 2a) and in simulations (Extended Data Fig. 5).

Training significantly increased dE and dM in adult pDp (Extended Data Fig. 9e-h). The increase in dE was largest between the conditioned odors (Trp, Ala) and smallest between non-conditioned odors. dM was increased predominantly in the direction from conditioned to other odors (i.e., from an activity vector representing a conditioned odor to a distribution of activity vectors representing another conditioned or a novel odor), with largest increases in the direction from the CS^+^ to non-conditioned odors (dM_Trp→Ser_, dM_Trp→His_; Extended Data Fig. 9g,h). These observations were largely abolished after shuffling of odor labels (Extended Data Fig. 6g-l). Hence, as observed in juvenile fish, training resulted in an odor-specific, asymmetric increase in dM that primarily enhanced the discriminability of responses to conditioned odors.

Consistent with these findings, manifold capacity in adult pDp was significantly higher in trained as compared to naïve fish (Extended Data Fig. 9i). Parameter-specific analyses showed that the effective radius, effective dimensionality, effective center alignment and effective axis alignment were all significantly lower in trained than in naïve fish (Extended Data Fig. 9j). Hence, as in juveniles, the increase in manifold capacity can be attributed to changes in the effective radius, effective dimensionality and effective center alignment of representational manifolds, which collectively outweighed a negative contribution of effective axis alignment. The time course of manifold capacity and the underlying geometrical and alignment measures was also similar to that in juveniles (Extended Data Fig. 9k). Together, these observations indicate that training resulted in consistent geometrical modifications of representational manifolds in juvenile and adult zebrafish that enhance the capacity for pattern classification.

### Manifold geometry predicts discrimination behavior

To examine whether the geometry of representational manifolds in pDp affects behavior we asked whether geometrical features of manifolds predict behavioral odor discrimination. Positive but non-significant correlations were found between the behavioral discrimination score (difference between appetitive behavioral response to CS^+^ versus CS^-^) and dE, cosine distance and LDA cross-validation accuracy (Fig. 6). Likewise, a non-significant negative correlation was found between the behavioral discrimination score and the Pearson correlation between activity patterns (*r* = -0.38, P = 0.06; not shown). A significant positive correlation was found between the behavioral discrimination score and dM_CS+→CS-_ (dM between a vector representing CS^+^ and the distribution representing CS^-^; *r* = 0.42, P = 0.04)) but not dM_CS-→CS+_ (Fig. 6). Moreover, a high significant correlation was detected between the behavioral discrimination score and manifold capacity (r = 0.56; P = 0.004; Fig. 6; see Extended Data Fig. 9l for results from adult fish). Hence, measures of separability based on geometrical features of representational manifolds predicted olfactory discrimination behavior in juvenile fish, indicating that geometrical modifications of representational manifolds contributed to learned changes in behavior.

**Fig. 6.**
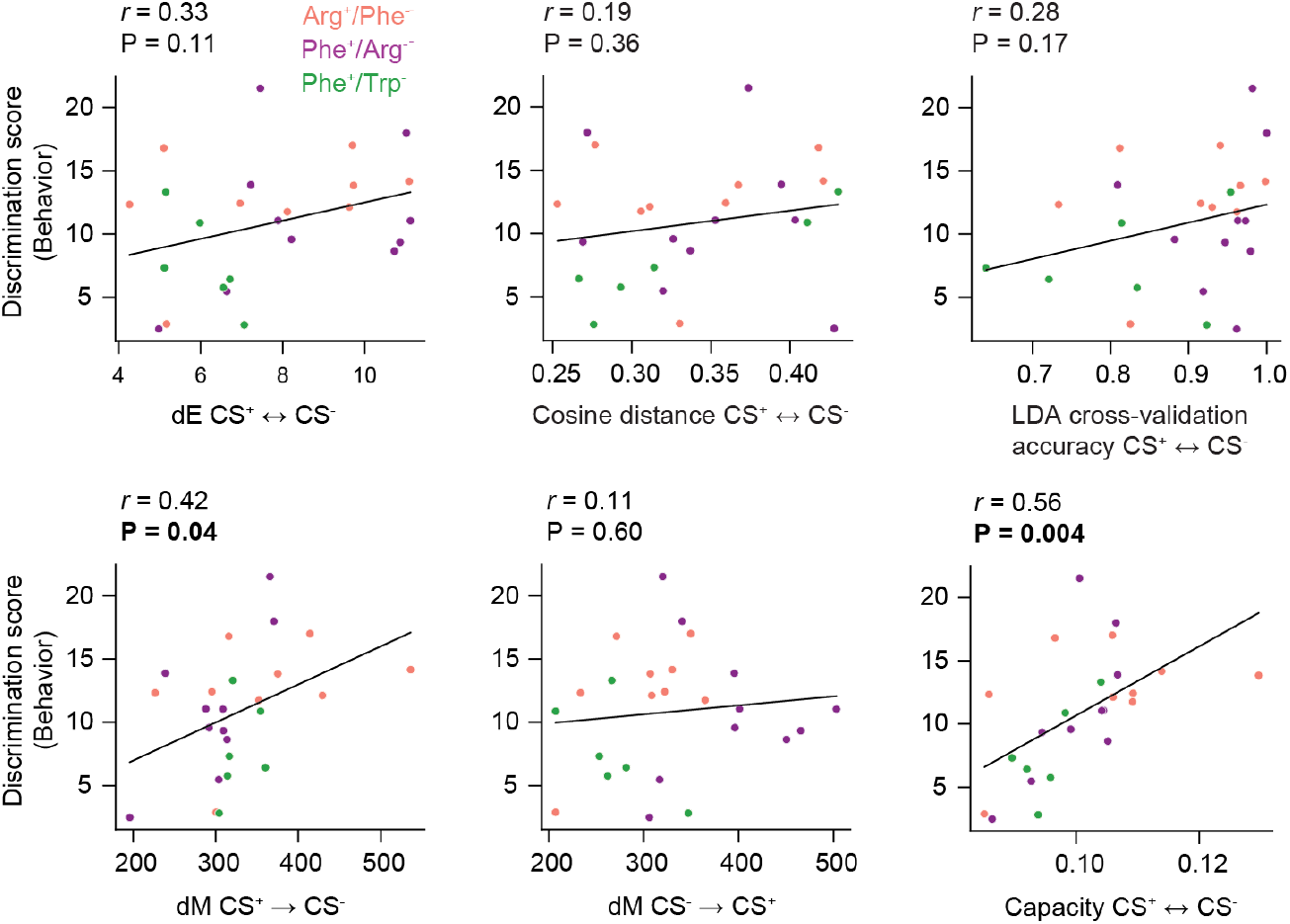
Separability of representational manifolds predicts odor discrimination behavior. Relationship between odor discrimination behavior and different measures related to the separability of representational manifolds (dE, cosine distance, LDA classification accuracy, dM, manifold capacity) across juvenile fish. Discrimination behavior was quantified by the discrimination score. Annotations show Pearson correlations (*r*) and the corresponding P-values.

## DISCUSSION

### Neural manifolds in an autoassociative network

We explored the organization of neural manifolds representing task-relevant information in an autoassociative memory network. Activity evoked by novel or conditioned odors did not exhibit obvious signs of classical or chaotic global attractor states. Nonetheless, analyses of distance relationships and manifold capacity demonstrated that training resulted in an untangling of task-relevant odor representations in pDp, indicating that pDp and, by extension, piriform cortex support odor classification across brain areas.

In principle, associative learning may be mediated by any mechanism that maps relevant inputs to defined activity subspaces according to their semantic relations. For example, activity patterns representing task-relevant sensory stimuli (inputs) may be associated with different task outcomes (semantic categories) by mapping them to separate neural manifolds (activity subspaces). Stimuli may thus be classified by a systematic organization of neural manifolds according to semantic categories, even if manifolds are not attractors. The formation of manifolds representing semantic categories is the basis for pattern classification by convolutional networks, which lack attractor dynamics by design^14,61^. Unlike convolutional networks, however, biological memory networks are often recurrent, pre-structured, and modified by local learning rules. The small size of pDp together with computational modeling and GLUE theory allowed us to directly analyze representational manifolds and their plasticity in an autoassociative area of the vertebrate brain.

The notion that neural manifolds organize relevant information in neuronal state space implies that geometrical and statistical properties of manifolds are relevant for neural computation^2^. Quantitative analyses of features such as the effective radius, dimension and alignments should thus drive insights into neuronal computations at the population level, similar to analyses of receptive fields at the single-neuron level. Consistent with this notion, defined manifold features were systematically modified by discrimination training and manifold capacity predicted discrimination behavior across individuals. These findings indicate that manifold geometry is linked to behaviorally relevant neuronal computations.

### Representational manifolds and olfactory memory

Our findings indicate that pDp contains a continuous representation of odor space that is modified, but not discretized, by constraining activity onto neural manifolds. This scenario is consistent with multiple experimental observations in pDp and piriform cortex: (1) Training of animals in olfactory memory tasks led to the emergence of task-specific responses but the continuous representation of odor space in naïve animals was not fundamentally reorganized (Figs. 1,2)^25,27,29-31^. This observation is consistent with the notion that pDp or piriform cortex do not store discrete items of information but jointly represent odor space and semantic information in continuous maps^19,20,28^. (2) Experimental results in pDp and piriform cortex revealed only minor effects of training on the variability of single-neuron responses or global population activity^25,30,31^. These observations are consistent with the assumption that representational manifolds constrain neuronal activity only in a subset of state space dimensions. (3) Accurate odor discrimination requires reliable readout of information from population dynamics despite high variability of neuronal activity in pDp and piriform cortex. Manifold-based representations can reconcile these issues because activity is constrained to subspaces and, thus, information can be retrieved reliably by integrating activity over the manifold^32,51,52^. (4) In pDp^42^, and possibly in piriform cortex^62^, network activity enters a state of precise synaptic balance during an odor response that may not support discrete attractor states. Representational manifolds can directly account for this observation and allow for pattern classification without attractor dynamics.

Discrimination training primarily enhanced the ability to classify conditioned or related odors. Hence, task-relevant information is combined with a map of odor space, resulting in joint representations of sensory and semantic information. These observations could, in theory, be explained by the formation of neuronal assemblies in synaptically balanced autoassociative networks^32^. Consistent with this hypothesis, training recapitulated diagnostic observations in simulated networks including the generalization of effects across related odors. However, direct mechanistic insights into the mechanisms defining manifold geometry will require reconstructions of network connectivity at synaptic resolution.

Behavioral training consistently increased manifold capacity and, thus, optimized representational manifolds for classification. Increased manifold capacity could, in part, be attributed to higher “compactness” (decreased effective radius) and lower correlation (decreased effective center alignment) of representational manifolds, in agreement with increases in dE and dM. Manifold capacity was further enhanced by a decrease in dimensionality. In computational models, a similar decrease in dimensionality could be attributed, in part, to a modest amplification of activity in an odor-specific direction^32^. Such a directional amplification of activity may also contribute to the decrease in axis alignment observed in pDp, which had a negative effect on manifold capacity. Hence, training may modify multiple geometrical features of manifolds with opposing effects on manifold capacity. In combination, however, positive effects outweighed negative contributions, resulting in an overall enhancement of the capacity for task-relevant odor classification. Importantly, geometrical features of manifolds predicted behavioral odor discrimination, consistent with the hypothesis that semantic information accumulated in the geometry of representational manifolds is read out by neural classifiers controlling behavior.

### Putative computational functions of representational manifolds

The representation of sensory and semantic information by neural manifolds supports task-dependent pattern classification by facilitating access to relevant information. The observed increase in manifold capacity implies that fewer neurons are required for efficient classification of conditioned odors after training, that readout becomes more robust, and that is becomes easier to find an acceptable classifier^15,37,52^. Experience-dependent changes in manifold geometry are therefore likely to facilitate and accelerate relevant learning processes. In addition, modifications of continuous representational manifolds support other computations such as the evaluation of metric relationships between relevant patterns^32^.

Interpretable changes in manifold capacity have recently been described also in brain areas of other species^37^. For example, manifold capacity was found to increase along the ventral stream of the visual system in datasets from monkeys and humans. In convolutional networks trained on image classification, manifold capacity increased systematically along layers prior to the final classification layer^14^. These studies further support the assumption that neural manifolds are informative objects critical for neuronal computation, implying that the systematic analysis of manifold geometry can uncover principles of information processing in the brain.

pDp and piriform cortex^25^ do not appear to function as an integrator network or to classify odor objects by convergent attractor dynamics even after learning. Hence, pDp is functionally more closely related to the intermediate (“representational”) layers than to the final classification layer of deep convolutional networks. We therefore propose that pDp establishes “semantic maps” of odor space to support behaviorally relevant sensory pattern classification and potentially other computations as part of a larger network. This view is consistent with the concept of olfactory cortex as a memory-related structure that comprises multiple recurrent brain areas^19,20^. More generally, representational learning by experience-dependent reorganization of neural manifolds may be critical for cognitive processes that are mediated by distributed computations across the pallium.

## METHODS

### Animals and transgenic lines

Zebrafish (*Danio rerio*) were raised and kept as groups in a standard facility at 26.5–27.5 °C on a 14/10 h light/dark cycle. Juvenile fish were kept in 1.1 L tanks (Tecniplast ZB11TK). Olfactory conditioning of juvenile fish in their home tank started 28-38 days post fertilization (dpf). Juvenile fish used for calcium imaging experiments were 46 – 56 dpf with a body length of 9.0 – 13.5 mm and not selected for gender, which is not yet determined at this stage. Adult fish were 5-6 months old and not selected for gender. The naïve group consisted of 4 females and 4 males while the trained group consisted of 5 females and 4 males. Juvenile fish expressed the calcium indicator GCaMP6s pan-neuronally under the control of the alpha-tubulin promoter (Tg[alphaTubulin:GCaMP6s]^50^). Adult fish were wildtype except for Tg[dlx4/6:ChR2-YFP]^63,64^ used in optogenetic experiments. All experimental protocols were approved by the Veterinary Department of the Kanton Basel-Stadt (Switzerland).

### Ex-vivo preparation of the zebrafish brain

#### Ex vivo preparation of the juvenile zebrafish brain

In juvenile zebrafish, one of the olfactory bulbs and the ipsilateral side of the telencephalon were exposed to provide optical access. Before dissection, juvenile zebrafish were transferred into a beaker containing c.a. 10 mL system water at room temperature. The beaker together with the fish was cooled with ice to 4°C to anesthetize the fish until immobile. Heart rate was visually monitored and anesthesia was ensured by pinching the tail fin. The fish was then transferred into ice-cold teleost artificial cerebrospinal fluid (ACSF: 124 mM NaCl, 2 mM KCl, 1.25 mM KH_2_PO_4_, 1.6 mM MgSO_4_, 22 mM D-(+)-glucose, 2 mM CaCl_2_ and 24 mM NaHCO_3_, pH 7.2.) bubbled with O_2_/CO_2_ (95%/5%)^65^ and fully immersed in this medium during dissection. Peripheral structures (eyes, jaws, gills) were removed and the head was detached from the body. The head was placed on a coverslip, the exposed lateral side was oriented upwards, and the preparation was stabilized using tissue glue (3M Vetbond Tissue Adhesives, No.1469SB). Fine forceps and scissors were used to expose the olfactory bulb and telencephalon by removing soft cartilage and connective tissue without damaging the olfactory nerve. Care was taken not to obstruct the nostrils. For fish with body length above 12 mm, contralateral bones were partially removed to increase exposure of the brain to aerated ACSF. The preparation was transferred together with the coverslip to a custom flow chamber for two-photon imaging that was continuously perfused with ACSF at room-temperature.

#### Ex vivo preparation of the adult zebrafish brain

The ex vivo preparation of adult zebrafish for the electrophysiology experiment was performed as described^42,49^. Briefly, fish were anesthetized by cooling to 4°C and decapitated in ACSF. After dissection of the jaws, the eyes and the bones covering the ventral telencephalon, the dura mater over pDp was removed with fine forceps. The ventral bones covering the olfactory bulbs were removed if necessary (electrophysiological recordings from mitral cells, optogenetic silencing). After surgery, the preparation was slowly warmed up to room temperature under constant perfusion with ACSF as described for juvenile fish.

Recent results from awake behaving fish show that the tuning, dynamics and correlation structure of activity measured in the *ex vivo* preparation is very similar to the *in vivo* situation (T. Caudullo and R.W.F.; unpublished observations), consistent with the finding that *ex vivo* activity measurements can predict behavior^58^. We therefore expect results obtained in the *ex vivo* preparation to be representative for the *in vivo* condition.

#### Injection of a synthetic calcium indicator into adult pDp

Fish were mounted in a custom-made holder for imaging in a horizontal orientation to ensure high reproducibility of the injection procedure as described^58^. Bolus loading of Oregon Green 488 BAPTA-1-AM (OGB-1; ThermoFisher Scientific) was performed as described^66^ with minor modifications. 50 μg of OGB-1-AM was dissolved in 30 μL of DMSO/Pluronic F-127 (80/20; ThermoFisher Scientific), vortexed for 1 min and sonicated for 15 min before being stored in 4 μL aliquots at −20°C. Prior to each experiment, an aliquot was diluted in 15 uL of ACSF, vortexed for 1 min, sonicated for 5 min and purified for 5 min at 5000 rcf in centrifugal filter units (PVDF membrane with 0.22 μm pore size; Millipore). A slowly tapering glass pipette was pulled from a borosilicate glass capillary (borosilicate glass capillaries, 1 mm outer diameter, wall thickness 0.21 mm, capillary length 100 mm; Hilgenberg article number 1810021) with a laser puller (P-2000; Sutter). The tip was broken under a microscope (MF-900 Microforge; Narishige) to a tip diameter of approximately 4 μm.

Pressure injections were targeted using a non-resonant two-photon scanning microscope with a 20x objective that provided both a fluorescent channel for detection of OGB-1 (PMT H7422P-40MOD, Hamamatsu) and a transmitted light channel (PSD, position-sensitive detector) as described^41^. pDp was identified based on its location in the lateral telencephalon, posterior to the prominent furrow. One primary injection was made ∼210 μm dorsal from the ventral-most aspect of Dp and ∼130 μm from the lateral surface of Dp in two intervals of each ca. 2 min. A second injection was performed in the same entry channel of the pipette but less deep (ca. 180 μm dorsal, 60 μm lateral) for 1-2 min. Dye injection was monitored by snapshots of multiphoton images. Pressure was adjusted to prevent fast swelling of the tissue that could be caused if the applied pressure was too high. The entire injection procedure typically took 6-10 min.

After injection, the pipette was retracted from the brain and the fish head was quickly (<2 min) transferred to another holder for imaging in a sagittal orientation using a 2-photon microscope equipped for volumetric resonance scanning. The approximately sagittal orientation allowed for improved optical access to areas of pDp close to the lateral brain surface but deeper from the ventral surface. The orientation of the brain was adjusted to minimize optical aberrations due to tissue curvature. Odor application and calcium imaging started ca. 1 h after dye injection.

### Optogenetic stimulation of the olfactory bulb

Blue light was targeted to the ipsilateral olfactory bulb through an optical fiber (200 μm diameter, Thorlabs) using a 457 nm laser (500 mW before attenuation) as described^49^. Laser intensity was adjusted to obtain 200 μW at the fiber tip. Optical stimulation was restricted to the olfactory bulb with possible minor off-target effects onto adjacent areas.

Odors were applied through a tube in front of the ipsilateral nostril using a peristaltic pump system as described^42^ (duration, 5 s or 10 s). In electrophysiological experiments the onset of optical stimulation (300 ms) was targeted at the plateau phase of the odor response in pDp, typically 400-500 ms after response onset. Current clamp recordings from superficial mitral cells (n = 4) showed that the latency between blue light onset and cessation of action potential firing was approximately 10–20 ms. Recovery of mitral cell activity was observed approximately 500 ms after the end of the light pulse. When activity was measured by 2-photon calcium imaging, optical stimulation started 1 s, 2 s or 3.2 s after response onset and lasted for 4 s. We also used 2-photon calcium imaging to specifically analyze calcium signals in Chr2YFP-expressing neurons during optical stimulation. However, no signals were detected, confirming that pDp neurons were not directly stimulated by light directed at the OB.

### Electrophysiology

Patch-clamp recordings of neurons in pDp were performed as described^42^ using borosilicate pipettes (pipette resistance: 4–8 MΩ) and a Multiclamp 700B amplifier (Molecular Devices). Pipettes were filled with intracellular solution containing (in mM): 132 Cs methanesulfonate, 10 Na_2_-phosphocreatine, 4 MgCl_2_, 4 Na_2_-ATP, 0.4 Na-GTP, 5 L-glutathione, 0.1 EGTA, and 10 HEPES (pH 7.2, 300 mOsm; all from Sigma). Current-clamp recordings were performed with an intracellular solution containing (in mM): 129 K-gluconate, 10 HEPES (free acid), 0.1 EGTA, 4 Na_2_-ATP, 10 Na_2_-phosphocreatine, 0.3 Na-GTP, 5 L-glutathione, and 13.1 KOH (pH 7.2, 305 mOsm; all from Sigma).

Neurons were targeted using the shadow-patching technique^67^ as described^42^, with 0.05 mM Alexa488 or Alexa594 (Invitrogen) included in the internal solution using a custom-designed video-rate multiphoton microscope^48^. When the dura mater over Dp was not completely removed, the pipette was advanced through the dura mater with transient high pressure (100 mbar) to avoid contamination of the pipette tip. Before forming a seal, neurons were approached within the tissue using low pressure (20 mbar). After break-in, series resistance and input resistance were continuously monitored. To measure the cellular time constant of Dp neurons by direct hyperpolarization, brief voltage steps (500 ms, step size ΔV = −5 mV) were applied through the patch pipette during voltage clamp experiments.

Voltage traces triggered on the onset of optical stimulation or hyperpolarizing current injections were averaged for each neuron and normalized by subtracting the pre-event current (mean of the 200 ms window before stimulation onset) and dividing by the asymptote (400-500 ms after stimulation onset). The resulting traces were fitted with an exponential function to estimate the decay time constant. An equivalent procedure was used to estimate decay time constants from normalized current traces.

### 2-photon calcium imaging

2-photon calcium imaging was performed using a custom-built multiphoton microscope^48^ using a 20x objective (NA 1.0; Zeiss) and custom-written Scanimage software^42,68^. Laser pulses for two-photon excitation were centered around 930 nm, with a temporal pulse width of 180 fs below the objective as measured with an autocorrelator (CARPE; APE Berlin). Fluorescence was detected by a GaAsP photo-muliplier tube (PMT, H7422P-40MOD; Hamamatsu) without bandpass filtering of the emitted light to maximize fluorescence yield. The average laser power was adjusted to 42-52 mW for juveniles and 23-24 mW for adults at imaging planes closest to the brain surface and increased by 4 mW at the deepest imaging planes using a Pockels cell (350-80LA; Conoptics) synchronized with the voice coil motor used for fast z-scanning.

Imaging was performed in 8 planes (256 x 512 pixels each) at 7.5 Hz as described^48^. Ca. 7.3 of these 8 planes were scanned during the linear trajectory of the z-scanning, while the remainder was acquired during the fast flyback of the z-scanning unit. The peak-to-peak maximum extension of the imaging volume was ca. 150 μm in juveniles and 100 μm in adults, whereas the extent of the FOV in x and y was around 250 μm and 125 μm in juveniles and 200 μm and 100 μm in adults, respectively, slightly varying with the relative z-position of the plane due to remote z-scanning^48^.

After every 1 or 2 trials, the microscope stage was repositioned in order to compensate for potential drifts. For this purpose, a small z-stack of ± 6 μm around the current location (step size, 2 μm) was acquired and the optimal shift in x, y and z was determined based on the correlation with a reference stack. Sub-resolution interpolation using a Gaussian fit of the correlation values allowed to achieve a correction accuracy well below the step size of 2 μm. For most experiments, the corrected drift was substantial over the time course of an experiment. In juveniles, the total drift ranged in 10-15 μm in pDp and 5-10 μm in the OB in z-direction, each region recorded for 80-90 min. In adults, the total drift ranged in 18-25 μm in z-direction over a full experiment within 50-60 min. During each trial, average projections of each imaging plane were created and inspected for artifacts (e.g., degraded signal due to bubbles or imaging plane drifts with respect to the reference stack). Defective trials were discarded and re-acquired. To measure neuronal activity during optogenetic stimulation of the olfactory bulb we scanned single optical planes at higher zoom at a frame rate of 30 Hz. Blue light was switched on/off for 22/11 ms, respectively, during each frame for 4 s. On/off cycles were synchronized to frame acquisition and image data were acquired only during the off phase (11 ms). Restricting image acquisition to 30% of the nominal frame time in a single plane reduced the field of view but allowed for optogenetic manipulations and image acquisition at high temporal resolution (30 Hz). Responses of the same neurons were measured in repeated trials with blue light stimulation starting at different interspersed time points (1 s, 2 s, 3.2 s after response onset). Usually, eight trials were acquired for each time point (in one fish, only five trials were acquired per condition). Only neurons showing strong odor responses were included in the analysis to ensure high signal-to-noise ratio (total of n = 22 neurons from 4 fish).

### Odor application

Amino acids (Ala, Arg, His, Phe, Ser, Trp; Sigma) were prepared as 100x stock solutions in double-distilled water, vortexed, sonicated, stored at -20°C, and diluted to a final concentration of 10^-4^ M in ACSF immediately before the experiment. Bile acid odorants (TDCA, TCA, GCA; Sigma) were also prepared as 100x stock solutions and diluted to a final concentration of 10^-5^ M. Food extract, which was used as a positive control stimulus, was generated by heating fish food (Gemma Micro 300) in double-distilled water and filtering the product through 0.22 μm pore size filters. The product was then stored as a stock solution and diluted 500x before the experiment.

Odors were applied to the nasal epithelium through a constant stream of ACSF using a computer-controlled odor-application system based on peristaltic pumps as described^42^. In experiments using juvenile fish, a circular symmetric 9-channel plastic manifold (Darwin microfluidics Manifold 9 Ports 1/4-28 PEEK, 1/16” OD) was used to combine the flow of multiple odor channels to the main ACSF perfusion channel. A bubble trap (Diba Omnifit® Bubble Traps, 21940-38) was placed in the tube delivering ACSF to the manifold. In experiments using adult fish, odors were applied using a linear manifold as described^42^. The time of response onset was determined by application of fluorescein in the absence of fish and fine-adjusted based on neural activity measurements in each experiment. Due to the fluid dynamics of aqueous media in narrow channels and the kinetics of media exchange in the flow chamber, odor offset was not abrupt but the concentration decayed gradually (approximately exponentially) over seconds. Both in juvenile and adult experiments, 1-3 odor applications were performed prior to the start of data acquisition to identify the odor-responsive brain region. Odors used in these pre-trials were food odor (adult fish) or a bile acid (juvenile fish).

To measure odor-evoked activity in pDp of juvenile fish, 6 odor stimuli (Phe, Arg, Trp, TDCA, TCA, GCA), a control stimulus (ACSF), and another control with no odor delivery were applied in a pseudo-random sequence. This procedure was repeated three times with different pseudo-random sequences. Odors were applied for 5 s with an inter-trial interval of at least 2 min. The same odor application procedure with new pseudo-randomized application sequences was repeated to measure odor-evoked activity in the olfactory bulb of the same fish. In experiments on adult fish, an equivalent protocol was used to measure activity in pDp using Trp, Ala, His, Ser, Food odor, TDCA, and ACSF as stimuli. Odors were applied for 10 s.

### Odor discrimination training

Adult zebrafish were trained in an odor discrimination task for six days as described^57^. Juvenile fish were trained using a similar procedure after pre-training in the home tank as a group (Temiz et al., manuscript in preparation). Naïve fish used as controls were obtained from the same parental stock. During the group training phase, 10-15 juvenile zebrafish (28-38 days post fertilization) were transferred into a housing tank (Tecniplast 3.5L tank, ZB30TK) that was continuously perfused with system water (26.5°C, 120–130 ml/min flow rate). During each trial, a CS^+^ or CS^-^ odor was delivered for 30 seconds using an Arduino-controlled peristaltic pump (Adafruit Peristaltic Liquid pump). Following CS^+^ delivery, fish food (Tetra TetraMin Flocken) was dispensed into a feeding ring floating in a specific location after a 30-second delay, whereas no food was dispensed following presentation of the CS^-^. Fish received 7 CS^+^ and 14 CS^-^ applications per day, presented in an alternating sequence with 40-minute intervals.

After 11 days of group training, fish were transferred individually to small tanks (Plastic-Haus AG, 12x6x6 cm) and allowed to acclimate for 1-3 days. Subsequently, each fish underwent individual training using the same procedure as for adult animals^57^ with minor modifications to account for differences in body size. CS^+^ and CS^-^ odors were the same as during the group training phase. Each odor was delivered in an alternating sequence for 9 trials with a 20-minute intertrial interval. Swimming behavior was monitored by 3D video imaging throughout the individual training phase, which lasted 2-4 days. The appetitive behavior score ζ for each odor was computed as described based on quantitative analyses of swimming speed, the z-position in the water column, the presence in the reward zone, water surface sampling, the distance to odor inflow tube and rhythmic circular swimming during the 30 seconds between odor onset and reward delivery^57^. ζ scores are combined and normalized measures similar to z-scores that quantify appetitive behavior in response to each odor. The behavior discrimination score was computed as the difference between the ζ scores for the CS^+^ and CS^-^, summed over trials and divided by the number of individual training days.

### Processing of image data

Automatic region of interest (ROI) extraction was performed using StarDist (https://github.com/stardist/stardist) and followed by manual correction to segment neuronal somata in the juvenile calcium imaging dataset. The model was trained with 20 RGB images (512 × 512), each color channel containing the anatomy image, ΔF/F_0_ map, and spatial correlation map as input and manually segmented neuron somata as ground-truth output. The ground-truth data was obtained from the same fish line using the same microscope as in the juvenile calcium imaging experiments. The threshold for ROI detection in StarDist was adjusted to avoid cross-neuron signal contamination. The ROIs detected by StarDist were imported into a custom MATLAB program (https://github.com/fmi-basel/neuRoi). ROIs were aligned between trials with fast Fourier transform (FFT) cross-correlation and individual neurons were manually redrawn or deleted depending on the quality of trial-by-trial alignment.

The raw fluorescence traces for each neuron were extracted by averaging the pixel intensity within the ROI and subtracting background fluorescence. The F_0_ for each neuron was taken as the median fluorescence intensity within a 2 s time window prior to stimulus onset.

Spike probability was inferred from ΔF/F_0_ traces using the CASCADE algorithm^43^. Global_EXC_7.5Hz_smoothing200ms_causalkernel was chosen as the spike inference model. The noise level was determined by pooling ΔF/F_0_ traces during a 2 s time window from all trials and used to choose the most appropriate noise level parameter used in the inference function. The output of CASCADE is the spike probability within 133 ms time bins (7.5 Hz frame rate). The spike probability of the CASCADE algorithm was converted to firing rate by multiplication with the frame rate.

Segmentation of neuronal somata in the adult calcium imaging dataset was performed manually using a custom MATLAB program (https://github.com/PTRRupprecht/Drawing-ROIs-without-GUI). Spike probability was inferred using the OGB_zf_pDp_7.5Hz_smoothing200ms model from CASCADE.

### Spiking network model of pDp

The spiking network model of pDp was the same as in a previous study^32^ except for the number of neurons and synaptic weights, to account for the smaller size of the juvenile zebrafish brain. The model of pDp consists of 1000 excitatory (E) and 250 inhibitory (I) adaptive leaky integrate-and-fire neurons with conductance-based synapses. The neurons in pDp receive inputs from 1500 excitatory mitral cells in the OB.

#### Neuronal dynamics

The membrane potential *V*_*x*_ of a neuron *x* in pDp evolved according to:

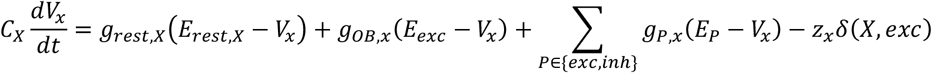

where X is the excitatory or inhibitory population to which x belongs. C_X_ is the membrane capacitance, g_rest,X_ is the leak conductance, and E_rest,X_ is the resting potential. The capital X in the subscripts means that the same values apply to all neurons in population X. g_OB,x_ is the conductance of the synapse from an olfactory bulb input to neuron x and g_P,x_ is the synaptic conductance from population P to neuron x. E_P_ is the reversal potential of a synapse of population P. z_x_ is the adaptation current if x is an excitatory neuron^69^:

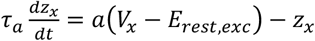

with z_x_ set to z_x_ + b after each spike. Parameters a, b and τ_a_ were the same as in a previous model^32^.

The conductances g_P,x_ evolved according to

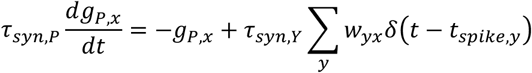

where τ_syn,P_ is the synaptic time constant, w_yx_ is the synaptic weight from neuron y to neuron x, and t_spike,y_ is the spike time of neuron y.

#### Olfactory bulb input

The input from the olfactory bulb to pDp was modeled by directly modulating the baseline 6 Hz firing rates of mitral cells in the olfactory bulb. Odor-evoked activities in the olfactory bulb were simulated by increasing the firing rate of 150 mitral cells and decreasing the firing rate of another 75 mitral cells. The activity pattern of mitral cells for each odor was generated to approximate the response amplitudes and pattern correlations observed in experiments.

#### Network connectivity and E/I assemblies

Connections between two neurons x and y from populations X and Y, respectively, were drawn from a Bernoulli distribution with parameter p_XY_. The strength of all existing connections was set to w_XY_. While parameters p_XY_ are the same as in a previous model^32^, w_XY_ were fitted to account for the reduced network size. To explore a broad parameter space, 4 connectivity matrices were fitted, each with 2 instantiations drawn, with parameter values shown in Table 1. The simulation results from these 8 instantiations were used in further analysis.

**Table 1.**
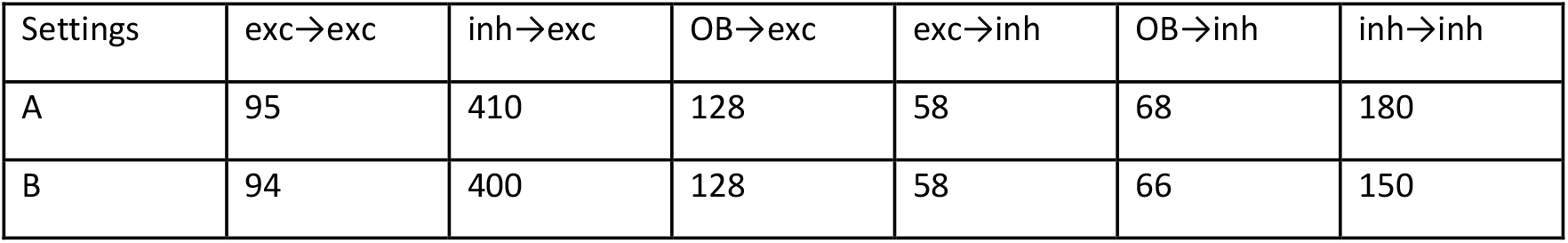
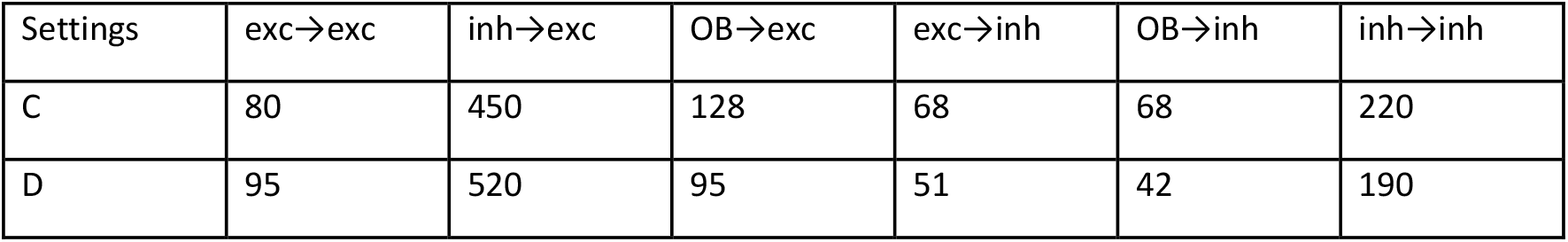
Weight strength parameter settings (Unit: pS)

| Settings | exc→exc | inh→exc | OB→exc | exc→inh | OB→inh | inh→inh |
| --- | --- | --- | --- | --- | --- | --- |
| A | 95 | 410 | 128 | 58 | 68 | 180 |
| B | 94 | 400 | 128 | 58 | 66 | 150 |
| C | 80 | 450 | 128 | 68 | 68 | 220 |
| D | 95 | 520 | 95 | 51 | 42 | 190 |

E/I assemblies were introduced in pDp as described^32^. The E-assembly comprised the 100 excitatory neurons receiving the highest degree of inputs from the mitral cells activated by the associated odor. The corresponding I-assembly comprised the 25 inhibitory neurons receiving the highest degree of inputs from the E-assembly. E/I assemblies were formed by increasing the number of connections within the E-assembly and between the E- and I-assembly. To maintain a constant number of connections, an equal number of connections was eliminated randomly between non-assembly neurons and assembly neurons. The network with assemblies in Fig. 3 has two sets of E/I assemblies corresponding to odor A1 and A2.

### Manifold construction and distance between manifolds

In experimental data, neural manifolds were constructed as follows: In each fish, the point cloud manifold representing an odor contained spike rate population vectors at each time point within the analysis time window, pooled over 3 trials (resulting in a total of 72 or 116 datapoints per manifold for the 3 s or 5 s time windows, respectively [24 or 39 timepoints × 3 trials]). Different starting points (0 s to 4 s after response onset) and durations (3 s, 5 s, 7 s) of the analysis window gave similar results (see also section “Time windows” below and Extended Data Fig. 10a). To compute dE and dM, 70 neurons (3 s time window) or 100 neurons (5 s window) were randomly subsampled from the manifold to ensure invertibility of the covariance matrix. Subsampling was repeated 50 times and results were averaged.

In simulated data, manifolds were constructed from population activity vectors containing spike rates in 100 ms time bins, covering the 2 s stimulus window. dE and dM were computed using 100 neurons that were randomly subsampled from the set of responsive neurons, which combined the 100 neurons with the highest activity in response to each odor. Results were averaged over 50 repeats.

dE between two manifolds X and Y was defined as:

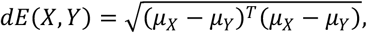

where 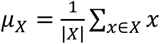 and 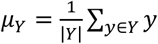 are the centers of the manifolds X and Y, respectively.

dM with X as sample and Y as reference was defined as:

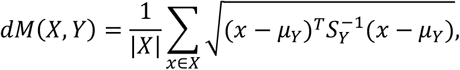

where |*X*| is the number of points in the manifold X, *x* is a point in the manifold X, μ_*y*_ is the center of the manifold Y, and *S*_*y*_ is the covariance matrix of the manifold Y. To compute dM, a small value (10^-5^) was added to the diagonal of the covariance matrix as a regularizer to avoid singularity.

To analyze distances between a pair of manifolds after shuffling of odor labels, data points of both manifolds were pooled and randomly assigned to two new sets of equal size. dE and dM were then computed between these sets and results were averaged over 50 repeats.

### Manifold capacity analysis

#### Data preprocessing and capacity estimation

Manifolds were constructed from experimental data as described above using an analysis time window of 7 s starting at the onset of the odor response unless noted otherwise. The time window was longer than for the analysis of dE and dM because manifold capacity can quantify separability even when manifolds exhibit curvature. Shorter time windows or different starting time points produced similar results. Time-resolved analysis (Extended Data Fig. 7d,e) was performed using a sliding 3 s time window. Data in Extended Data Fig. 7c were computed using a 5 s time window starting 1 s after odor onset as for LDA (Fig. 2e).

We ensured that point cloud manifolds were linearly separable after subsampling of neurons and time points. In juvenile fish, 700 neurons and 140 activity patterns were sampled for each manifold. In adults, 400 neurons and 80 activity patterns were sampled. The number of neurons for juvenile and adults were chosen to match the minimal number of neurons recorded at each developmental stage. Capacity analysis of simulated data was performed using a 2 s time window, 1000 neurons, and 50 repeats.

Capacity was computed separately for each pair of manifolds representing an odor pair^37^. Subsampling was repeated 50 times for each manifold pair and capacity and geometric measures were averaged over the 50 repeats. As a control, manifold points were shuffled by pooling data points from both manifolds and re-assigning labels randomly prior to the calculation of capacity. The global centering and bias parameters were both set to true during capacity computation, which translated the mean of the manifold centers to the origin and allowed for the hyperplane classifier to have an offset from the origin^37^.

#### Manifold capacity: overview

The algorithm for computing manifold capacity is detailed in ref. ^37^ and summarized in Supplementary Note 1. Here we briefly describe the key steps. A manifold is modeled as a convex set residing in an *N*-dimensional state space. It is represented by its center, its *K* axes stretching out from the manifold center, and the set of coordinates with respect to the manifold axes, specifying the location of the points belonging to this manifold. That is, denoting the manifold by *M*, a point *x* in the manifold can be expressed as

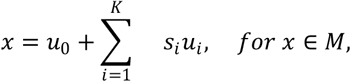

where *u*_0_ is the manifold center, *u*_*i*_ ∈ *R*^*N*^ for 1 ≤ *i* ≤ *K* < *N* is the set of manifold axes, and *s* ∈ *R*^*K*^ is the coordinate of *x* with respect to the manifold axes. We use *S* ⊂ *R*^*K*^ to denote the set of all possible coordinates of points contained in the manifold.

Consider *P* manifolds simultaneously residing in the *N*-dimensional state space. Each manifold *M*^μ^ is indexed with μ for 1 ≤ μ ≤ *P*. Consider a set of dichotomies *Y* ⊆ {-1,1}. As defined by GLUE theory^37^, the manifold capacity α is:

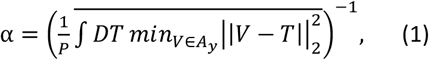

where *DT* is the zero-mean Gaussian measure and *T* is a random vector with dimension *N*. Finally, *A* is a convex set of vectors reflecting the geometry of the manifold shapes:

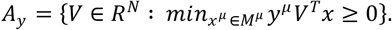

The set *A*_y_ is the collection of all linear classifiers for the dichotomy *y*, where *V* corresponds to the normal vector that uniquely defines a separating hyperplane.

The overline in equation (1) denotes the average with respect to the labels *y*.

#### Effective geometric measures for linear classification

According to GLUE theory, capacity is affected by geometric measures including effective radius, dimension, and alignments. “Effective” emphasizes that the measures are analytically connected to the capacity value; hence, this approach can be used to analyze geometrical changes underlying changes in manifold capacity. Thus, manifolds with different intrinsic geometries can have the same effective measures due to their relative configurations.

Detailed mathematical expressions of these measures are introduced in ref. ^37^ and summarized in Supplementary Note 1. Here we only give an intuitive overview of how they are computed.

To understand the effective geometric measures, it is important to introduce the concept of anchor points. The capacity formula (Equation 1) is a Gaussian average (over *T*) of a quadratic programming problem (a convex optimization problem). According to the strong duality theory, the solution of a quadratic program is equal to the solution of its dual problem. In this case, the dual problem is a function of the points from the manifolds. Concretely, let 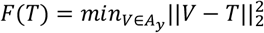, the duality theory gives that there exists a function *g* such that *F*(*T*) = *g*(*T, x*^1^(*T*), …, *x*^*P*^(*T*)) for some *x*^μ^(*T*) ∈ *M*^μ^ for each μ. These points *x*^1^(*T*), …, *x*^*P*^(*T*) are known as anchor points. As a result, the randomness of *T* Induces a distribution over each manifold. These distributions are the anchor point distributions. Notice that the anchor point distributions are analytically connected to the capacity value by the following equation:

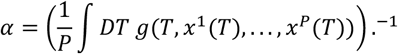

Finally, the effective geometric measures from GLUE theory are simply geometric terms extracted from the function *g* in the above formula. For reference, the following are the intuitive definitions of the geometric measures (see ref ^37^ and Supplementary Note 1 for details):

1. Effective radius measures how far the points spread away from the manifold center, and is normalized by the length of the manifold center. It can be seen as the amplitude of internal variability of the manifold.
2. Effective dimension measures the number of directions along which the points extend, with respect to the manifold center. It represents the degree-of-freedom of the variability.
3. Effective center-alignment measures the correlation between the centers of different manifolds. It is the cosine similarity under the anchor-point geometry.
4. Effective axis-alignment measures the correlation between the internal axis of different manifolds.

### UMAP

Neuronal population activity was embedded in a low-dimensional space using Uniform Manifold Approximation and Projection (UMAP)^70^. Neurons from fish in the same condition were pooled. UMAP embeddings were computed separately for each condition on activity patterns within a 5 s time window starting 1 s after odor onset including all trials and all odors. Two-dimensional UMAP was applied with a Euclidean metric and fixed hyperparameters (n_neighbors = 70, min_dist = 0.6 for inferred firing rate and n_neighbors = 100, min_dist = 0.8 for ΔF/F). To enable direct comparison across training conditions, embeddings were aligned to the centered embedding from the naïve condition with an orthogonal Procrustes transformation from the CEBRA package^71^.

### Linear Discriminant Analysis and support vector machine

Linear separability of neural manifolds was quantified using pairwise Linear Discriminant Analysis (LDA) or a support vector machine. Neurons were randomly subsampled to a fixed number. Each odor manifold consisted of all activity patterns within a 5 s time window starting 1 s after odor onset from all three trials. For each pair of manifolds, an LDA classifier or a support vector machine (both implemented in Python scikit-learn package) was trained to discriminate odor identity. The Ledoit-Wolf shrinkage of the classifier was set to 0.001 to avoid singular covariance matrices in LDA. The performance of the classifier was evaluated using 3-fold cross-validation. Accuracy values were averaged the three cross-validations and 50 repeats of neuron subsampling for each odor pair.

### Statistical analysis, correlation analysis and time windows

Comparisons between naïve and trained groups were performed using a Mann–Whitney U test. A nonparametric Kruskal–Wallis test followed by a post hoc Dunn’s test was used for multiple comparisons with the naïve group. Reported P values are adjusted for multiple comparisons with Bonferroni correction. The variance of cosine distances was compared between training groups using a one-sided F-test after averaging of cosine distance matrices over fish. The correlation between variables was evaluated using ordinary least squares (OLS) linear regression. In all statistical tests (implemented in Python with scipy.stats or scikit_posthocs package), P < 0.05 was considered statistically significant. Asterisks indicate standard significance levels (n.s.: P ≥ 0.05; * P < 0.05; ** P < 0.01; *** P < 0.001).

Different analyses depend differently on the length of the analysis time window. Generally, time-averaging can obscure information and generate artifacts when manifold geometry is complex, which can be partially avoided using short time windows. However, shortening the analysis time window obviously reduces information about manifold geometry. By default we therefore used different time windows depending on the analysis to be performed. For analyses of time-averaged activity we chose a short time window (3 s time window starting 1 s after response onset) to avoid artifacts. UMAP and LDA were performed using a time window of intermediate length (5 s starting 1 s after response onset) because activity patterns are not averaged over time but weighted equally. For manifold capacity analysis we used the longest time window (7 s starting at response onset) because this analysis preserves individual datapoints and explicitly extracts geometrical information. To ensure robustness of our conclusions we also performed all analyses using a common intermediate time window (5 s duration starting 1 s after response onset) and obtained very similar results (Extended Data Fig. 10a). Moreover, manifold capacity analysis was also performed with a sliding time window of 3 s (Extended Data Fig. 7d,e).

Previous results demonstrated that the inference of neuronal firing rates from fluorescence measurements (ΔF/F) using CASCADE is reliable and attenuates noise^43^. To confirm that this procedure did not introduce critical errors we performed two controls. First, we added a substantial amount of noise (0.5 SDs of the signal amplitude) to the firing rate data and repeated all analyses. Second, we performed all analyses using the raw ΔF/F data. Both manipulations had only minor effects on the main results and did not alter the main conclusions (Extended Data Fig. 10b,c).

## Acknowledgements

We thank members of the Friedrich lab for insightful discussions. This work was supported by the Novartis Research Foundation, by the European Research Council (ERC) under the European Union’s Horizon 2020 research and innovation program (grant agreement no. 742576), and by the Swiss National Science Foundation (grants no. 31003A_172925/1, 31003A_152833/1). C.C. and S.C. acknowledge support from the Center for Computational Neuroscience at the Flatiron Institute of the Simons Foundation. S.C. is also supported by the Klingenstein-Simons Award, a Sloan Research Fellowship, NIH award R01DA059220, and the Samsung Advanced Institute of Technology (under the project “Next Generation Deep Learning: From Pattern Recognition to AI”).

## CODE AVAILABILITY

Code for simulations is available at https://github.com/clairemb90/pDp-model_juvenile and code for analysis is available at and https://github.com/aloejhb/pDp_manifold_analysis and https://github.com/aloejhb/catrace.. The code for manifold capacity analysis is provided as Supplementary Software (zip file).

## DATA AVAILABILITY

The experimental data and the simulation results that support the findings of this study are available in Figshare (https://figshare.com/s/e1440b31c98c34721336; *private link will be replaced by public link with doi after acceptance for publication*).

## AUTHOR CONTRIBUTIONS

B.H. participated in behavioral and calcium imaging experiments on juvenile fish, made major contributions to data analysis at all levels, and participated in writing the manuscript. N.Z.T. participated in behavioral and calcium imaging experiments on juvenile and adult fish, contributed to data analysis, and participated in writing the manuscript. C.-N.C. and S.C. contributed to data analysis (manifold capacity analysis) and participated in writing the manuscript. P.R. performed calcium imaging and electrophysiological experiments in adult fish. C.M.-B. performed simulations and contributed to data analysis. B.T. contributed to behavioral and calcium imaging experiments in adult fish. R.W.F. organized and supervised the study, contributed to data analysis, and wrote the manuscript together with other co-authors.

## INCLUSION AND ETHICS STATEMENT

All authors have fulfilled the criteria for authorship required by Nature Portfolio journals and the study followed the ethical guidelines of Nature Portfolio journals.

---

**Extended Data Fig. 1.**
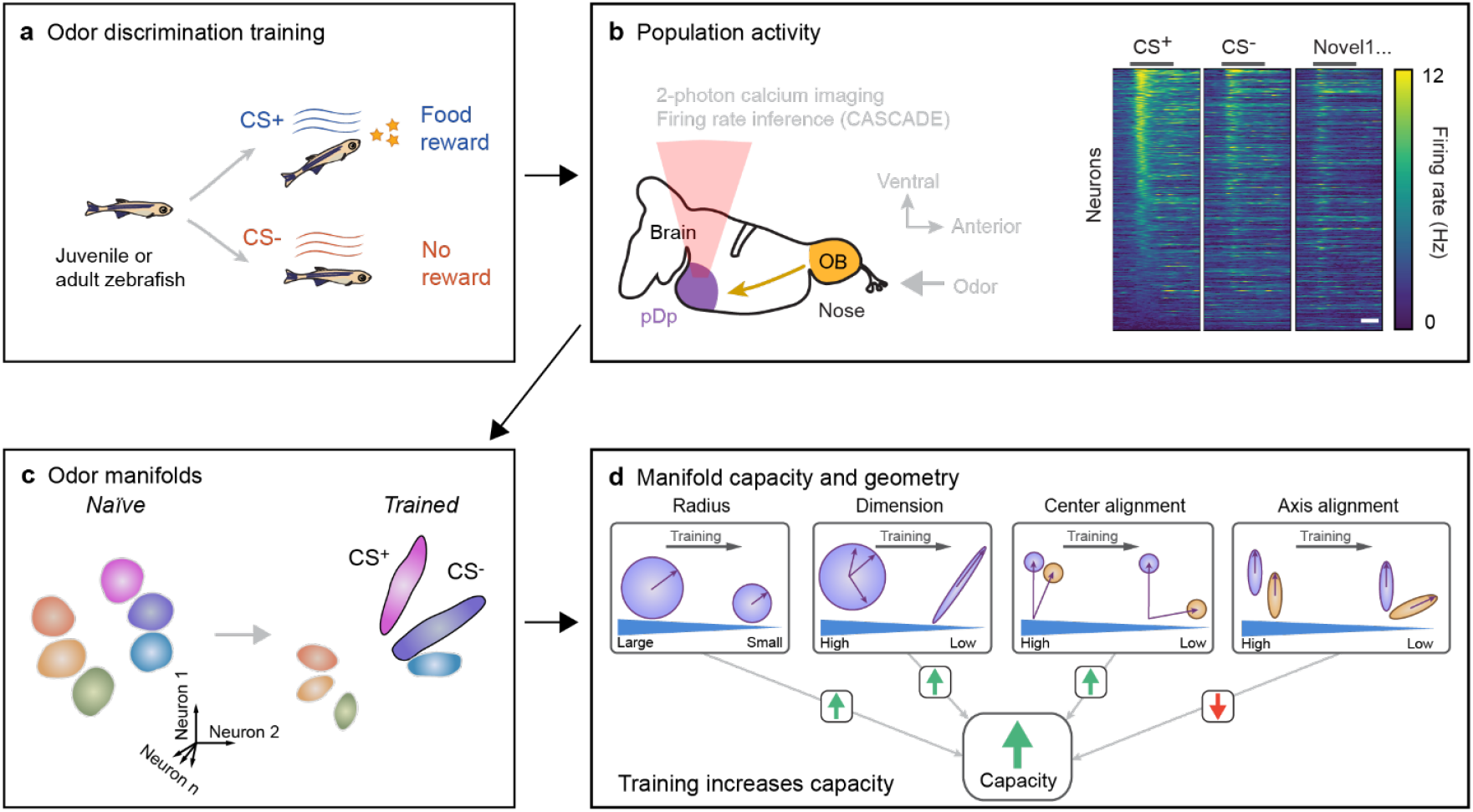
Graphical summary of approach and main results. **a**, Juvenile or adult zebrafish were trained in an odor discrimination task. **b**, Population activity evoked by conditioned (CS^+^, CS^-^) and novel odors was measured in telencephalic area pDp, the homolog of piriform cortex, using 2-photon calcium imaging and firing rate inference. **c**, Training enhanced the separation of neural manifolds representing conditioned odors (CS^+^, CS^-^) from representations of other odors. **d**, Analyses based on *manifold capacity theory* demonstrated that training enhanced the linear separability, or “untangledness”, of manifolds representing conditioned odors. Increased manifold capacity (separability) could be attributed to changes in multiple geometrical features: (1) a decrease in the effective radius (*manifolds become more “compact”*), (2) a decrease in effective dimensionality (*manifolds become more “flat”*) and (3) a decrease in effective center alignment (*manifolds become more decorrelated*). A concomitant decrease in effective axis alignment (*manifolds become less aligned*) had a negative effect on manifold capacity that was, however, outweighed by changes in the other features. The increase in manifold capacity was correlated to odor discrimination performance across individuals, indicating that geometrical features of representational manifolds are closely related to the behavioral readout of odor representations.

**Extended Data Fig. 2.**
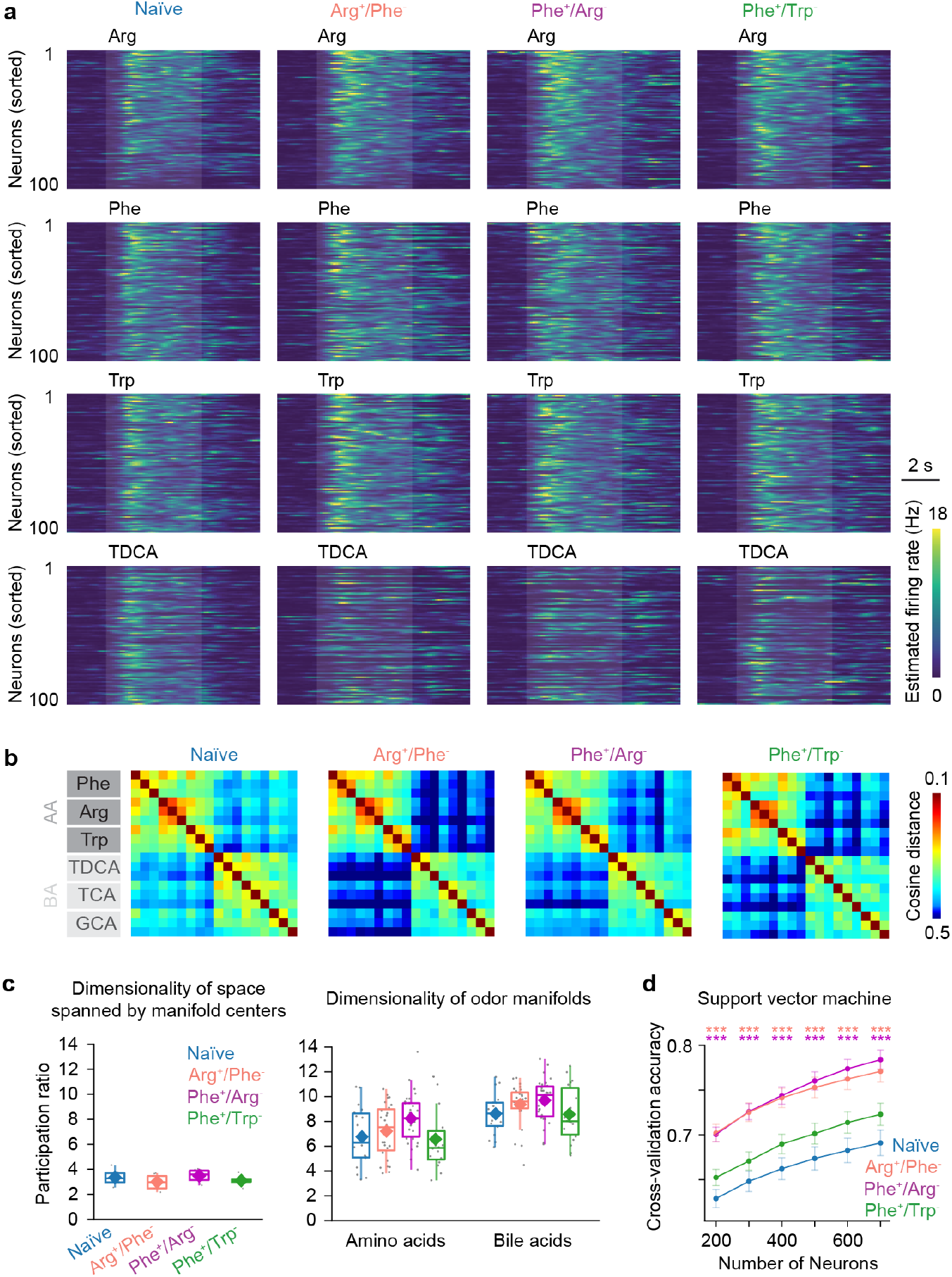
Odor-evoked population activity: additional observations. **a**, Inferred firing rates of neurons responding to four different odors (3 AAs, 1 BA) in naïve and trained fish. For each training group (column), panels show the activity of 100 randomly selected neurons from one representative fish, sorted by their response to Arg. Transparent shading depicts odor stimulation. **b**, Mean cosine distances between odor-evoked activity patterns in all training groups. Note that cosine distances were more broadly distributed in trained groups. **c**, Left: dimensionality (participation ratio) of the space spanned by manifold centers in each training group. The maximum possible dimensionality is six (number of odor manifolds). Each datapoint represents one fish. Right: dimensionality of the point cloud manifolds representing individual odors, analyzed separately for amino acids and bile acids across training groups. Each datapoint represents one manifold in one fish. **d**, Cross-validation accuracy of pairwise odor classification by a support vector machine as a function of neuron number (subsampling) across training groups (mean ± SEM; n = 50 repeats). Asterisks: statistical significance for comparison to Naïve group (***, P < 0.001).

**Extended Data Fig. 3.**
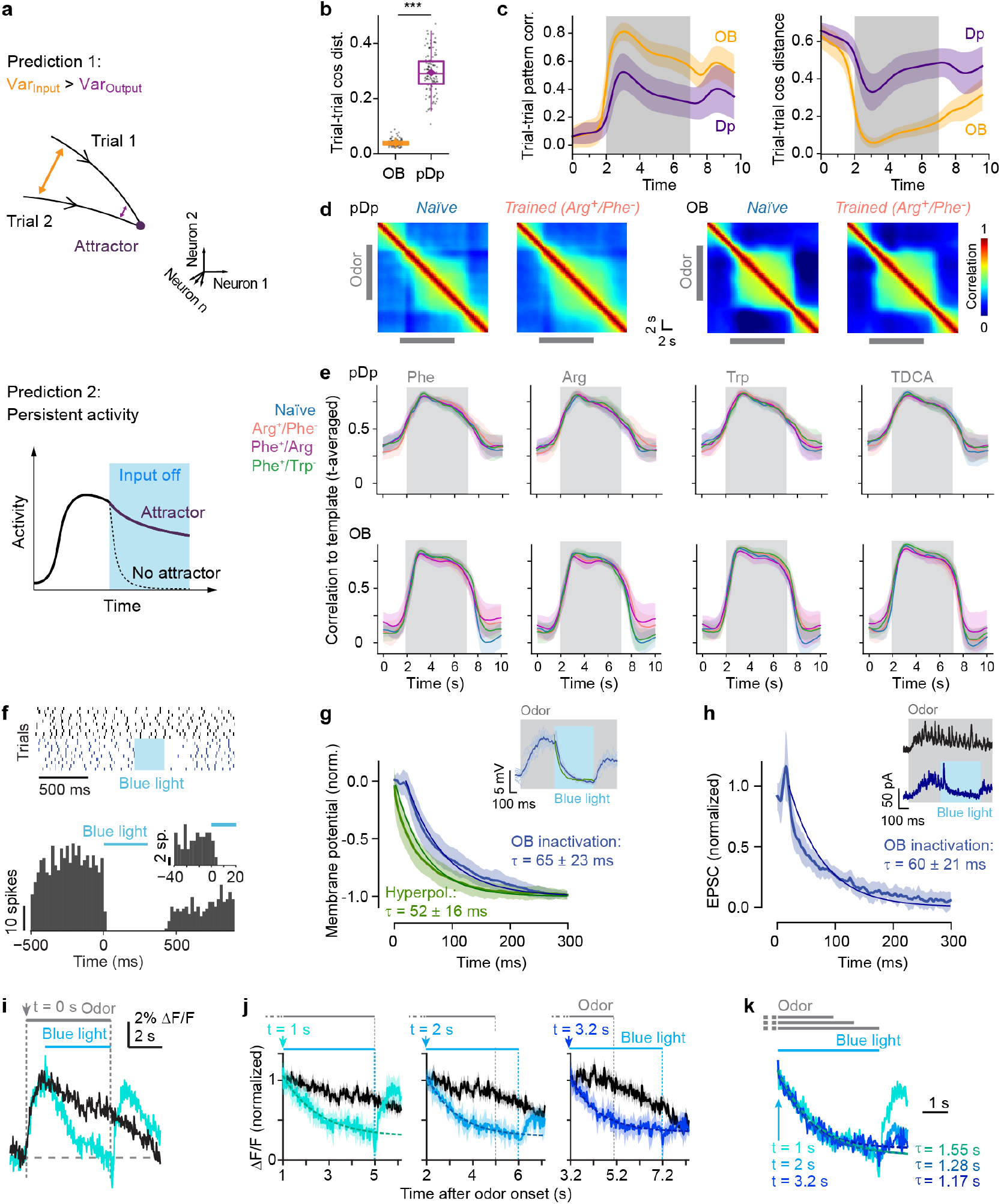
Analysis of attractor dynamics. **a**, Schematic: predictions for attractor networks. Prediction 1: variability decreases due to convergent dynamics. This prediction is based on the fact that activity patterns evoked by the same odor at different times or in different trials exhibit considerable variability due to stochastic firing of individual neurons and other physiological, behavioral and environmental factors. Prediction 2: activity persists after cessation of input. This prediction assumes that the sensory input is *not* part of the system itself. In other words, the prediction is based on the hypothesis that the dynamics is dominated by *intrinsic* activity, i.e., the system exhibits *intrinsic* attractor dynamics, as observed in integrator circuits^4^. To test prediction 1, we analyzed odor-evoked activity in the olfactory bulb and pDp of the same fish over time (**b, c**). To test prediction 2, we analyzed changes in activity patterns over time (**d**,**e**). Moreover, we optogenetically silenced sensory input to pDp during odor responses and examined the subsequent decay of activity in pDp. Optogenetic silencing of the olfactory bulb within 10 – 20 ms was achieved using Tg[dlx4/6:Chr2YFP] fish, which express channelrhodopsin-2 fused to YFP in superficial inhibitory interneurons of the olfactory bulb^63,64^ (**f**). Activity in pDp was recorded intracellularly using whole-cell current- and voltage-clamp recordings (**g**,**h**) or by 2-photon calcium imaging after bolus-loading of Oregon Green 488 BAPTA-1 (OGB-1; **i**-**k**)^43,58,66^. **b**, Variability of odor-evoked firing rates in the olfactory bulb (OB) and pDp of the same fish, quantified by the cosine distance between activity patterns evoked by the same odors in different trials. Activity was averaged over 3 s, starting 1 s after response onset. Variability was significantly higher in pDp (mean ± SD: 0.29 ± 0.07, 3 trials; 6 odors; N = 6 fish) than in the olfactory bulb (0.04 ± 0.01; Mann–Whitney U test, P = 6.1 × 10^-37^), contradicting attractor-based predictions. **c**, Trial-to-trial variability of odor-evoked activity patterns as a function of time in the OB and pDp of the same fish. Variability was quantified by the Pearson correlation (left) or cosine distance (right) between activity patterns evoked by the same odors in different trials. After odor onset, variability initially decreased but subsequently increased in the presence of the odor. This observation does not support the hypothesis that pDp exhibits convergent attractor dynamics and pattern completion. The observation that variability remains elevated after stimulus offset does also not support the hypothesis that pDp remains in an attractor state after stimulus withdrawal. However, post-stimulus activity is difficult to interpret because it comprises multiple interacting components including a prominent off-response (Extended Data Fig. 4). **d**, Pearson correlation between activity patterns at different time points in pDp (top) and in the olfactory bulb (bottom), averaged over all Arg trials of naïve (left; N = 6) and Arg^+^/Phe^-^-trained fish (right; N = 9). **e**, Mean Pearson correlation between activity patterns at different time points and the mean activity during the odor application period (gray shading) in pDp (top) and in the olfactory bulb (bottom). Curves show correlations of activity evoked by four odor stimuli (Phe, Arg, Trp, TDCA) in each training group (colors; mean ± SD). Note that activity patterns changed rapidly at the end of the odor stimulus even though the mean activity remained elevated (Fig. 1d; odor concentration decayed slowly after stimulus offset; Methods). **f**, Top: action potentials of a mitral cell in different trials (rows) during odor application without (black) and with (blue) optogenetic stimulation of inhibitory interneurons (cyan shading; 300 ms). Bottom: Peri-stimulus time histogram of spiking activity around the time of optical stimulation (blue bar; 4 mitral cells from two fish, 109 trials, 20 ms time bins; inset: zoom-in, 5 ms bins; note rapid silencing). **g**, Mean membrane potential change in pDp neurons after optogenetic silencing of the olfactory bulb (blue) and upon injection of a hyperpolarizing step current (green, 500 ms; n = 17 pDp neurons from 6 fish; total of 177 trials; shaded area shows SD). Individual traces were normalized prior to averaging. Smooth lines show single-exponential fits averaged over neurons; annotations show mean time constants (± SD). The first 20 ms were excluded for fits to data with optical stimulation to account for delayed silencing. Inset: membrane potential time course of a single neuron during odor stimulation (gray shading) and optogenetic silencing of olfactory bulb output (cyan shading). Thin traces show individual trials (n_trial_ = 9); dark blue trace shows average; green line shows average response to step current injection. **h**, Mean normalized excitatory postsynaptic current (EPSC) during an odor response after optogenetic silencing of olfactory bulb output (voltage clamp recordings; n = 9 pDp neurons from 4 fish, total of 54 trials). Traces from individual neurons were normalized prior to averaging. Smooth line shows single-exponential fits averaged over neurons (excluding first 20 ms); annotations report mean time constant ± SD. Inset: EPSCs evoked by the same odor in the same neuron in single trials without (top) or with (bottom) optogenetic silencing of olfactory bulb output (cyan shading). Note that upon silencing of the OB, the membrane potential (**g**) and synaptic currents (**h**) does not persist but decays with a time constant similar to the membrane time constant. These observations indicate the absence of attractor dynamics, even if an attractor state was not yet reached at the time of silencing. **i**, Two-photon calcium imaging of activity in pDp evoked by a strong odor stimulus (mixture of Ala, Arg, Trp; 100 μM each; 5 s) in the absence of blue light (black) and with blue light stimulation starting 1 s, 2 s or 3.2 s after response onset (green; average over n = 22 responsive neurons from 4 fish; frame rate, 30 Hz). Gray/blue bars show duration of odor/light stimuli. The calcium signal decreased during light stimulation, which was followed by a rebound response. **j**, Comparison of calcium signal kinetics without (black) and with (colored) light stimulation at different time points t after the onset of the odor response. Dashed lines show exponential fits. **c**, Overlay of the traces in **b**, aligned on light onset. Annotations indicate time of light onset relative to the onset of the odor response (left) and the time constant of exponential fits (right). Time constants are consistent with the expected decay of somatic/nuclear fluorescence signals upon rapid silencing^66^ and similar across time points. Although this approach does not allow for precise quantitative measurements of the decay kinetics of electrical activity the results imply that activity decays rapidly, independent of the time point of silencing.

**Extended Data Fig. 4.**
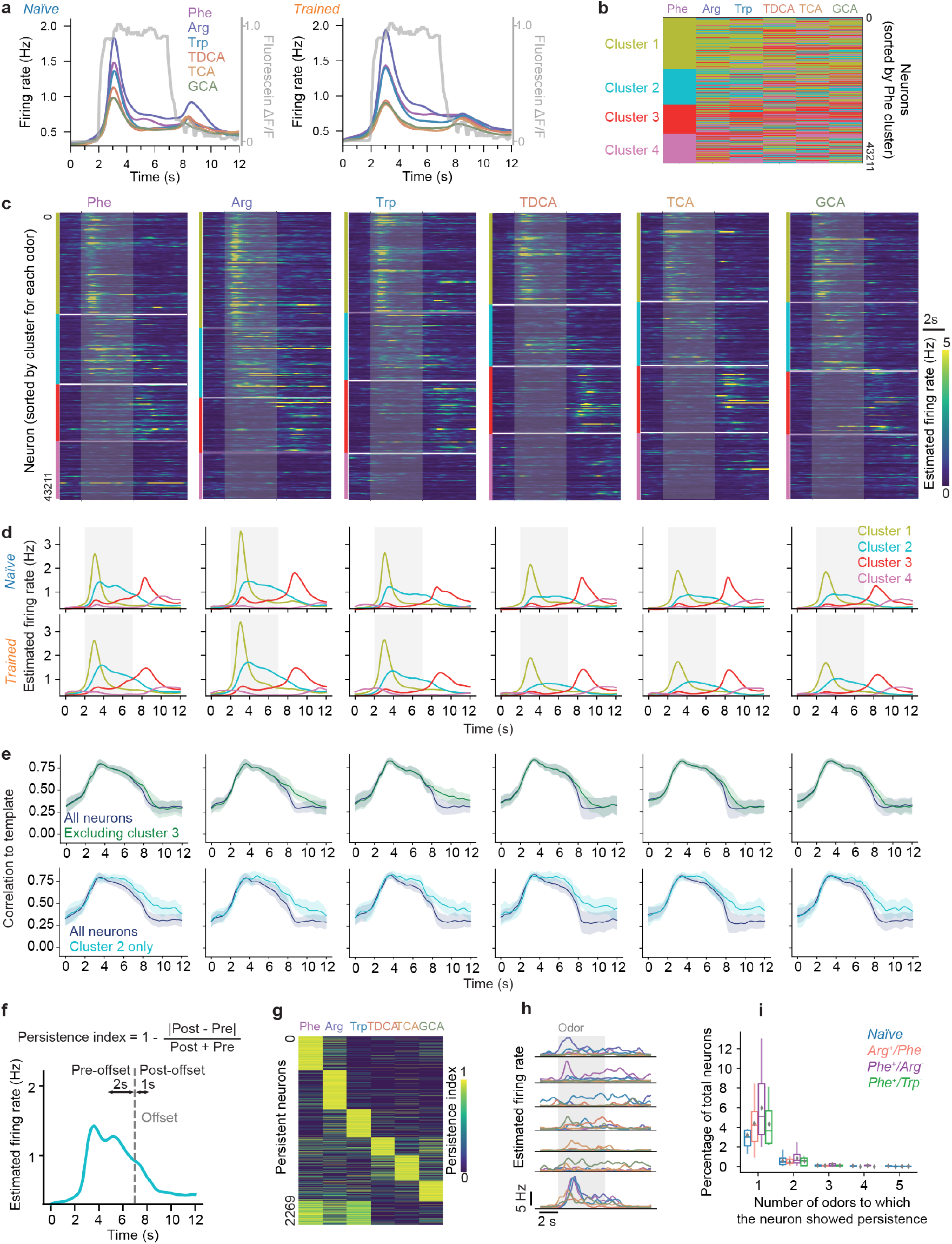
Time course analysis and post-stimulus activity. To analyze post-stimulus activity in more detail we compared the time course of activity to the time course of odor concentration (**a**). Note that odor concentration increased rapidly but decayed gradually after stimulus offset with an approximately exponential time course due to slow washout. Concentration remained elevated for at least 5 s after stimulus termination. Furthermore, we sorted neurons by k-means clustering into four clusters based on their firing rate time courses, independently for each odor (**b**,**c**). Clusters representing consistent temporal patterns were obtained across odors without manual forcing. Neurons with elevated firing rates that outlasted stimulus presentation were consistently enriched in cluster 2 while clusters 1 and 3 represented neurons whose activity decreased or increased, respectively, after stimulus onset (**c**-**e**). Furthermore, we defined a persistence index to identify neurons with elevated activity that extended beyond stimulus offset (**f**). On average, 5.4% ± 3.4% of neurons (N = 31 fish) showed a persistent response to at least one odor. The subset of persistent neurons was odor-specific and slightly but not significantly higher in trained fish (**g**-**i**). These results show that many neurons exhibit a classical off-response at the end of stimulation. In addition, odor-specific neuronal subsets show slowly decaying activity that outlasts the nominal duration of the stimulus. This activity may, in principle, reflect attractor states maintained by subsets of neurons in pDp or connected brain areas. Alternatively, or in addition, this activity may be driven by the gradual decline in odor concentration. **a**, Time course of stimulus concentration (gray trace), measured by fluorescein application (average over four repetitions). Colored traces show mean odor-evoked firing rates in naïve and trained fish (same as Fig. 1d,g). **b**, Clustering of all neurons (trained and naïve) based on their firing rate traces (k-means; four clusters). Colors depict cluster identity; neurons were sorted by cluster identity for Phe. **c**, Inferred firing rate of all neurons, sorted by cluster (color-coded) for each odor. As clustering was performed separately for each odor, the order of neurons differs between odors. **d**, Mean firing rates of each cluster as a function of time for each odor. **e**, Mean Pearson correlation between activity patterns at different time points and the mean activity during the odor application period, averaged over all neurons. Correlations were computed with all neurons (dark blue), after excluding the offset-responsive cluster 3 (green), or using only neurons from the persistent cluster 2 (cyan). **f**, Definition of persistence index for each neuron. Pre- and post-offset activities were computed as the median estimated firing rates during 2 s before odor offset and 1 s after odor offset, respectively. Neurons were defined as “persistent” when their persistence index was >0.9 and their pre-offset firing rate was >1.5 Hz. Cyan trace shows firing rate time course of cluster 2 in response to Phe (same as in **d**, top left). **g**, Persistence index of neurons defined as “persistent” in response to at least one odor, sorted by clustering based on odor-specificity. **h**, Mean firing rates of example neurons showing persistent responses to one or multiple odors. Trace colors depict odor identity as in other panels. **i**. Percentage of neurons showing persistent responses to one or multiple odors, separated by training groups. Box plots show median over fish, 1^st^ and 3^rd^ quartiles; whiskers show 1.5 times the interquartile range; diamonds show mean. On average, 5.4% ± 3.4% of neurons (N = 31 fish) showed a persistent response to at least one odor. The fraction of neurons with persistent activity was higher in trained fish but this effect was not statistically significant.

**Extended Data Fig. 5.**
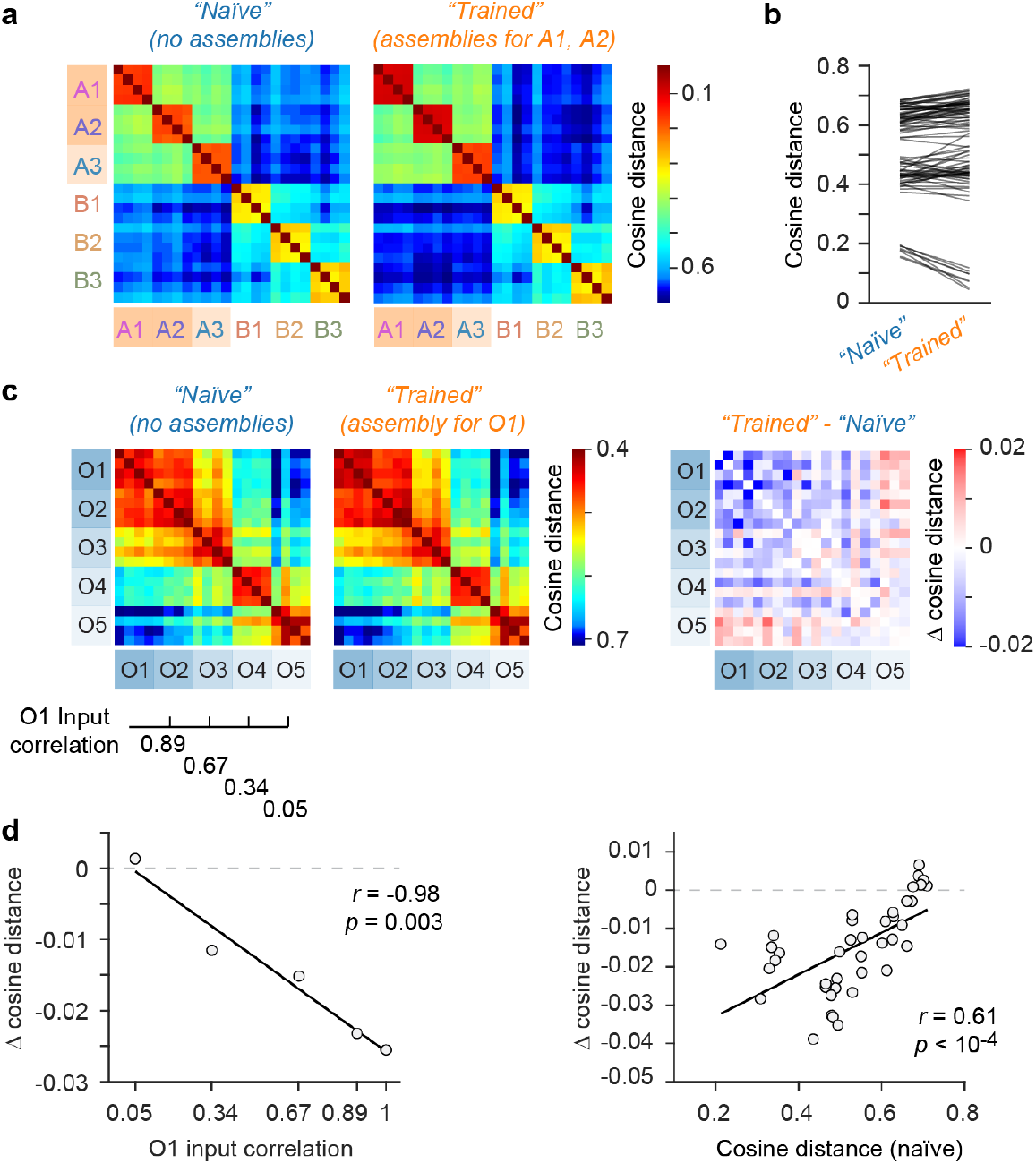
Computational modeling: additional results I. **a**, Cosine distance between activity patterns evoked by simulated odors (3 A-odors, 3 B-odors; four trials each) in randomly connected networks (*“naïve”*) and in precisely balanced networks with assemblies representing odors A1 and A2 (*“Trained”*). Matrices were averaged over results from eight “*Naïve”* networks and a corresponding set of eight “*Trained”* networks (Methods). Each *“Trained”* network was generated by rewiring a small number of connections in the corresponding *“Naïve”* network to create the assemblies^32^. Hence, the majority (∼80%) of connections in each *“Trained”* network were identical to the corresponding *“Naïve”* network. **b**, Each line shows the cosine distance between activity patterns evoked by two simulated odors in a *“Naïve”* and the corresponding *“Trained”* network (total of 100 randomly chosen comparisons from all odor pairs and networks). Note that cosine distances that are large in *“Naïve”* networks tend to become even larger in *“Trained”* networks while small cosine distances tend to become slightly smaller. As a consequence, the distribution of cosine distances was broader in *“Trained”* networks (p = 0.02; F-test). Because networks differ only in a small fraction of their connections, this broadening is likely to reflect a generic effect of assemblies that systematic modifies distances between activity patterns depending on the similarity between the corresponding input patterns. **c**, To further explore this hypothesis we simulated responses to five input patterns (virtual odors O1 – O5; four trials each) in randomly connected networks (*“Naïve”*) and in corresponding precisely balanced networks with a single assembly representing O1 (*“Trained”*). Correlations between O1 and the other input patterns (“O1 input correlation”) decreased systematically from O2 through O5. Matrices on the left show cosine distances averaged over sets of eight corresponding *“Naïve”* and *“Trained”* networks; the matrix on the right shows the difference. Note that differences in cosine distance depended systematically on the cosine distance in *“Naïve”* networks. **d**, Mean difference in cosine distance between activity patterns in *“Trained”* and *“Naïve”* networks as a function of O1 input correlation (left) and as a function of the cosine distance between activity patterns in the *“Naïve”* network (right). Lines show linear fits; annotations show Pearson correlation and statistical significance. Together, these simulations indicate that assemblies have small effects on the similarity relationships between activity patterns that systematically depend on the similarity of input patterns (odors). In particular, low (small positive) cosine distances were further decreased. As a consequence, cosine distances between activity patterns representing odors of intermediate similarity became more broadly distributed. This generic effect resembles the effects of training on cosine distances in juvenile (Fig. 2a, Extended Data Fig. 2b) and adult fish (Extended Data Fig. 9d).

**Extended Data Fig. 6.**
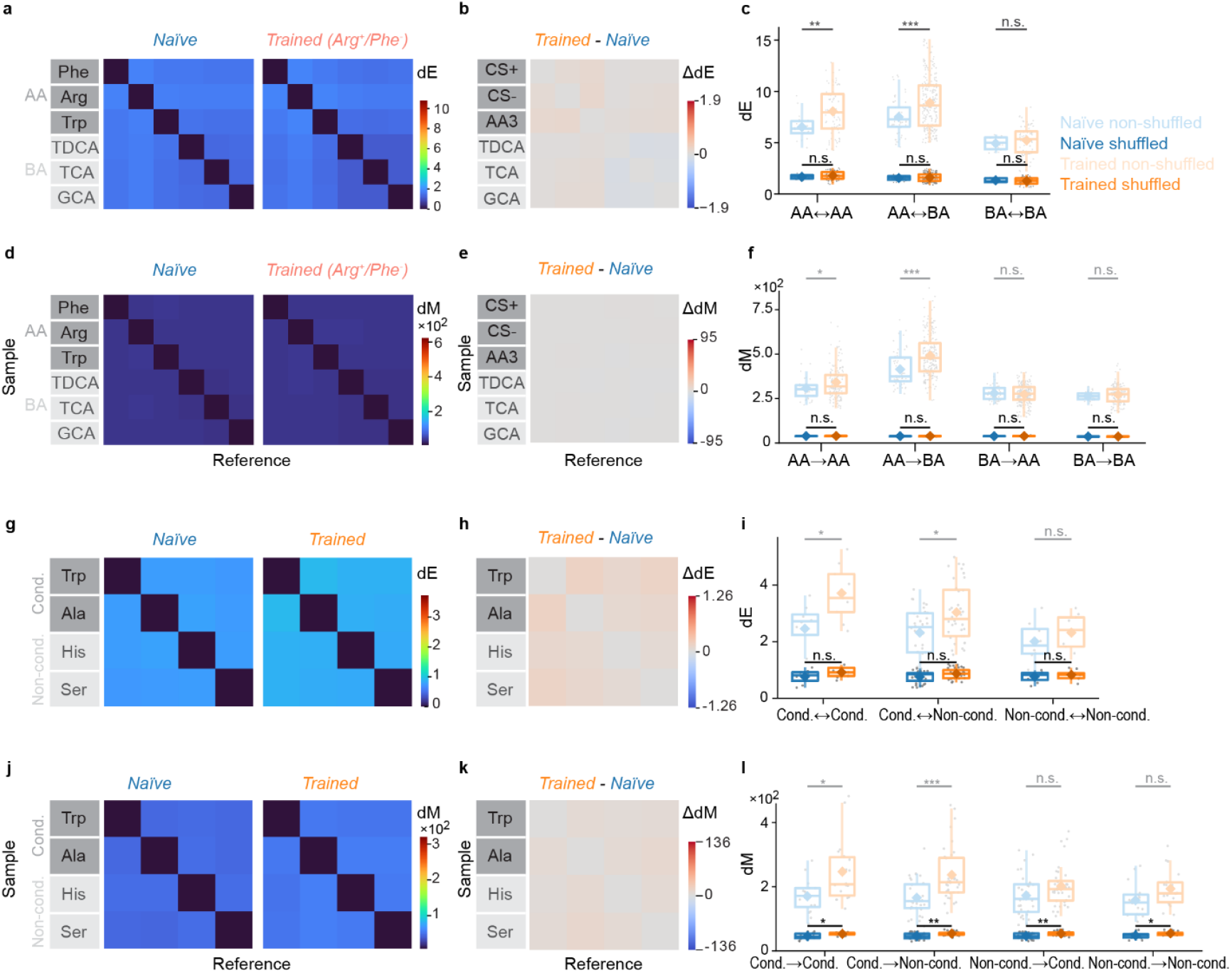
Distances between pairs of manifolds after shuffling manifold labels. **a**, Pairwise dE between manifolds constructed after shuffling labels of datapoints representing different odors. Left: naïve juvenile fish (matrix averaged over N = 6 fish); right: Arg^+^/Phe^-^-trained fish (N = 9 fish). The shuffling was performed such that for each pair of manifolds, all points from both manifolds were polled and randomly split into two new manifolds. In all panels, color scales are the same as in the figures showing corresponding results without shuffling (Fig. 4). **b**, Difference between dE matrices from trained and naïve fish after shuffling. Trained fish were from different training groups were combined (N = 25 in tital). Amino acid odors were reordered by reward assignment (CS^+^, CS^-^, third amino acid) prior to averaging over training groups. **c**, dE between manifolds representing different odor classes (AA: amino acids; BA: bile acids) after shuffling of data points from naïve (N = 6) and trained (N = 25) fish. Pale box plots show dE before shuffling (same data and axis scaling as in Fig. 4). For all panels: ns, P ≥ 0.05; *, P < 0.05; **, P < 0.01; ***, P < 0.001; see Supplementary Note 2 for more details. **d**-**f**, analysis of dM after shuffling of datapoint labels. Same procedures and plot conventions as in **a**-**c. g**-**l**, same analysis of dE and dM for data from adult fish after shuffling of datapoint labels. Color and axis scales are the same as in the figures showing corresponding results without shuffling (Extended Data Fig. 9). In **i** and **l**, manifold distances were analyzed separatedly for conditioned odors (Cond.; Trp and Ala) and non-conditioned odors (Non-cond.; Ser and His).

**Extended Data Fig. 7.**
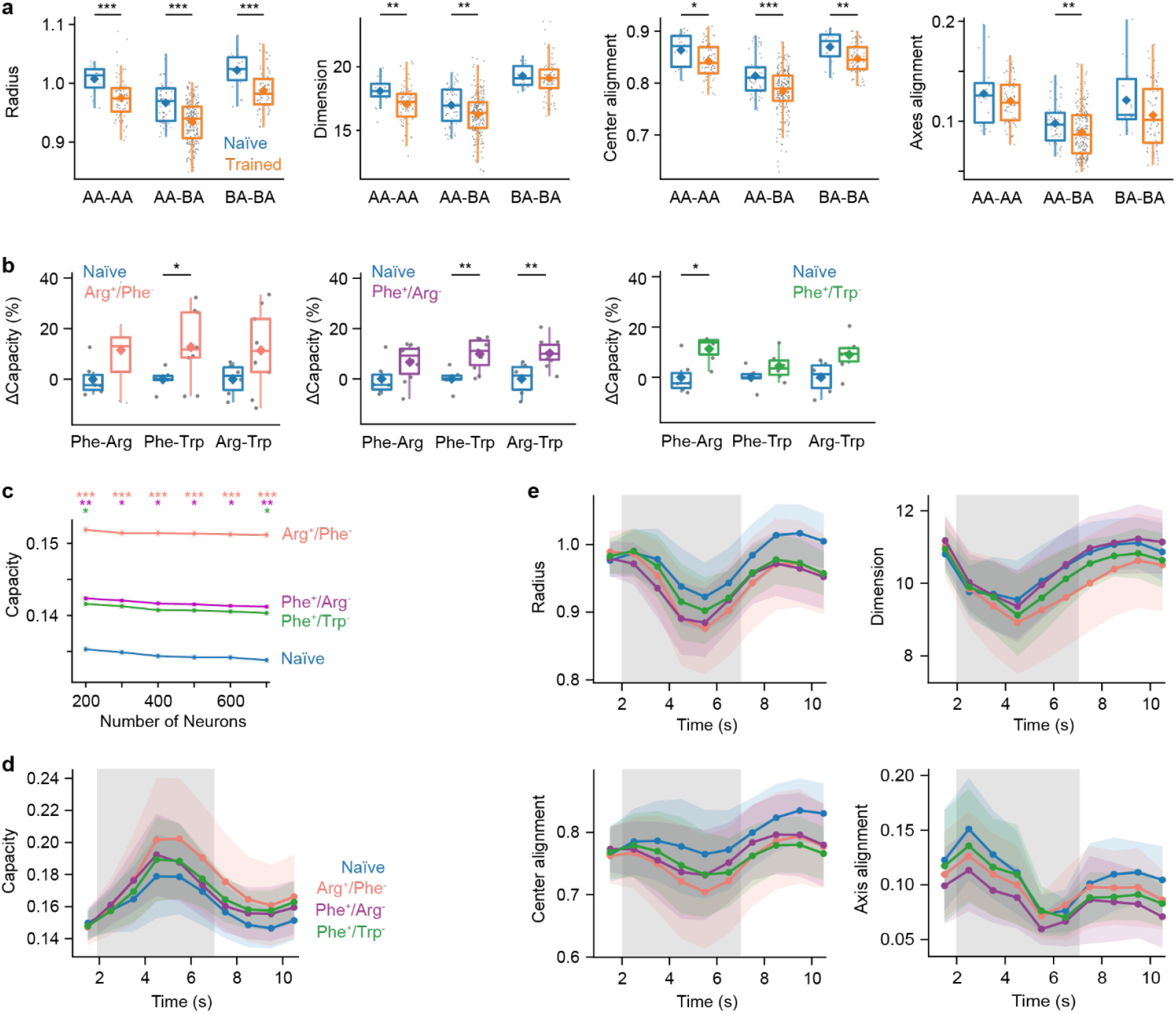
Geometrical measures contributing to manifold capacity: additional analyses. **a**, Effective geometric measures of manifolds representing different classes of odor pairs (AA: amino acids; BA: bile acids) based on data from naïve (N = 6) and trained fish (N = 25; all training groups combined). For all panels: ns, P ≥ 0.05; *, P < 0.05; **, P < 0.01; ***, P < 0.001; see Supplementary Note 2 for more details. **b**, Change in manifold capacity in each training group relative to the naïve group for each pair of amino acids. Capacity was consistently higher in trained fish but statistical significance was reached only in one case (Phe-Trp in Phe^+^/Arg^-^ fish; P = 0.002), possibly because the number of datapoints for comparison of single odor pairs in individual training groups is low. See Fig. 5 for statistical comparisons of pooled data. **c**, Manifold capacity as a function of the number of randomly subsampled neurons (n = 50 repeats; mean ± SEM over repeats). Note that manifold capacity is almost independent of the number of subsampled neurons, contrary to classification accuracy using LDA (Fig. 2e) or support vector machines (Extended Data Fig. 2d). This reflects the fact that manifold capacity reflects the minimum number of neurons (dimensions) required for classification, independent of the total population size. **d**, Manifold capacity as a function of time in different training groups (same as Fig. 5d). Capacity was determined in 3 s time windows centered on the specified time points. Gray shading depicts odor stimulation. Capacity is higher in trained fish and increases during stimulus presentation. Note that the apparent decrease in capacity prior to stimulus offset can be attributed to the fact that the analysis time window extended beyond the duration of the stimulus. **e**, Contributions of geometrical features to manifold capacity as a function of time. All measures are effective measures. Conventions as in **d**.

**Extended Data Fig. 8.**
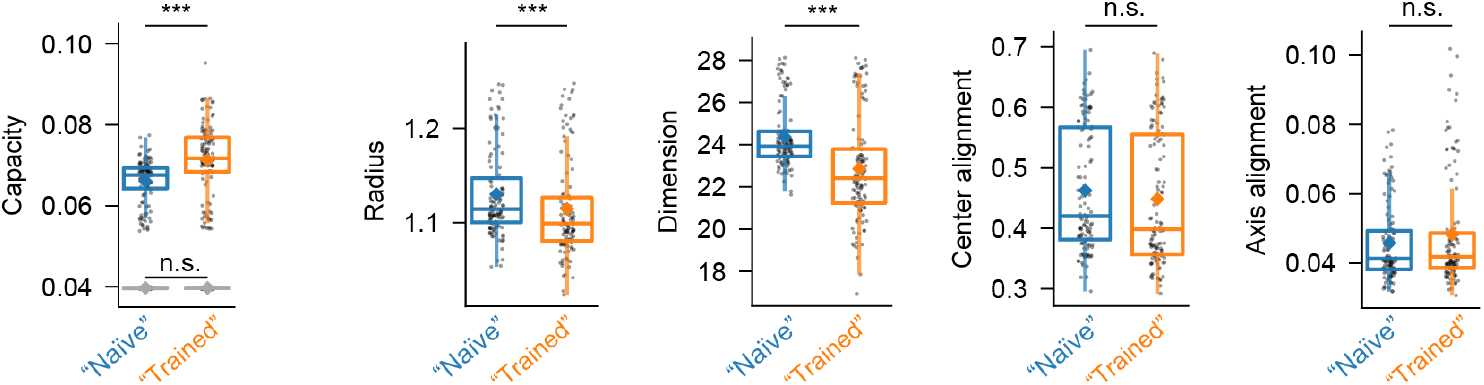
Manifold capacity analysis of simulated activity. Manifold capacity and effective geometric measures of neural manifolds in randomly connected spiking network models (“Naïve”; n = 120 odor pairs from N = 8 network configurations) and in network models containing E/I-assemblies representing two input patterns (“Trained”; n = 120 odor pairs from N = 8 network configurations). For all panels: ns, P ≥ 0.05; ***, P < 0.001; see Supplementary Note 2 for more details. Gray: results after shuffling of labels.

**Extended Data Fig. 9.**
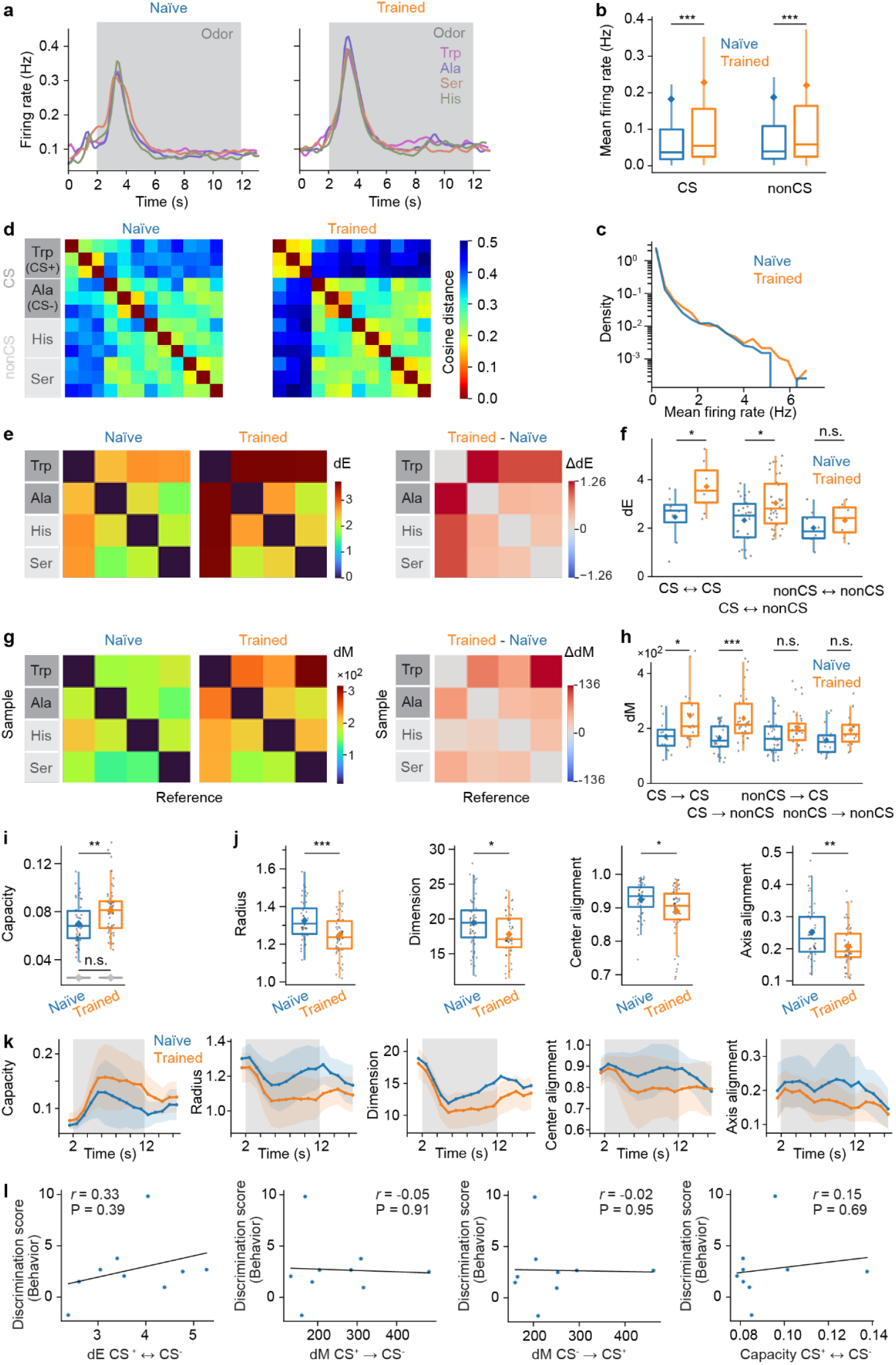
Learning-related plasticity of odor representations in pDp of adult zebrafish. **a**, Mean time course of inferred firing rates in response to each odor in naïve (N = 8 fish; left) and trained adult zebrafish (N = 9; right), averaged across all neurons, trials and fish. Gray shading indicates odor presentation (10 s). **b**, Response amplitude evoked by conditioned odors (CS; CS^+^: Trp, CS^-^: Ala) and non-conditioned odors (nonCS; Ser, His) was significantly higher in trained than in naïve fish. For all panels: ns, P ≥ 0.05; *, P < 0.05; **, P < 0.01; ***, P < 0.001; see Supplementary Note 2 for details. **c**, Amplitude histogram of responses to odors used for conditioning (Trp, Ala) in naïve and trained fish. Density is normalized by the area under each curve. **d**, Mean cosine distance between activity patterns evoked by different odors (3 trials each) in naïve and trained fish. **e**, Matrix of mean dE in naïve and trained adult zebrafish (left) and difference matrix (right). **f**, dE between pairs of manifolds representing different odor categories . **g**, Matrix of mean dM in naïve and trained fish (left) and difference matrix (right). **h**, dM between pairs of manifolds representing different odor categories. **i**, Manifold capacity was significantly higher in trained fish (gray: after shuffling of labels). **j**, Comparison of effective geometric measures contributing to manifold capacity in naïve and trained fish. **k**, Time course of manifold capacity and the underlying geometrical and alignment measures, determined in 3 s time windows centered on the specified time points. Shading depicts odor stimulation. **l**, Correlations between distance measures and behavioral discrimination score in adult zebrafish. Each datapoint represents one fish. Unlike in juvenile zebrafish (Fig. 6), correlations were not statistically significant, possibly due to small sample size. Moreover, adult fish received less training (no group training prior to individual training; Methods), different odors were used for training and stimulation, pDp is substantially larger in adults than in juveniles, and neuronal plasticity processes may differ.

**Extended Data Fig. 10.**
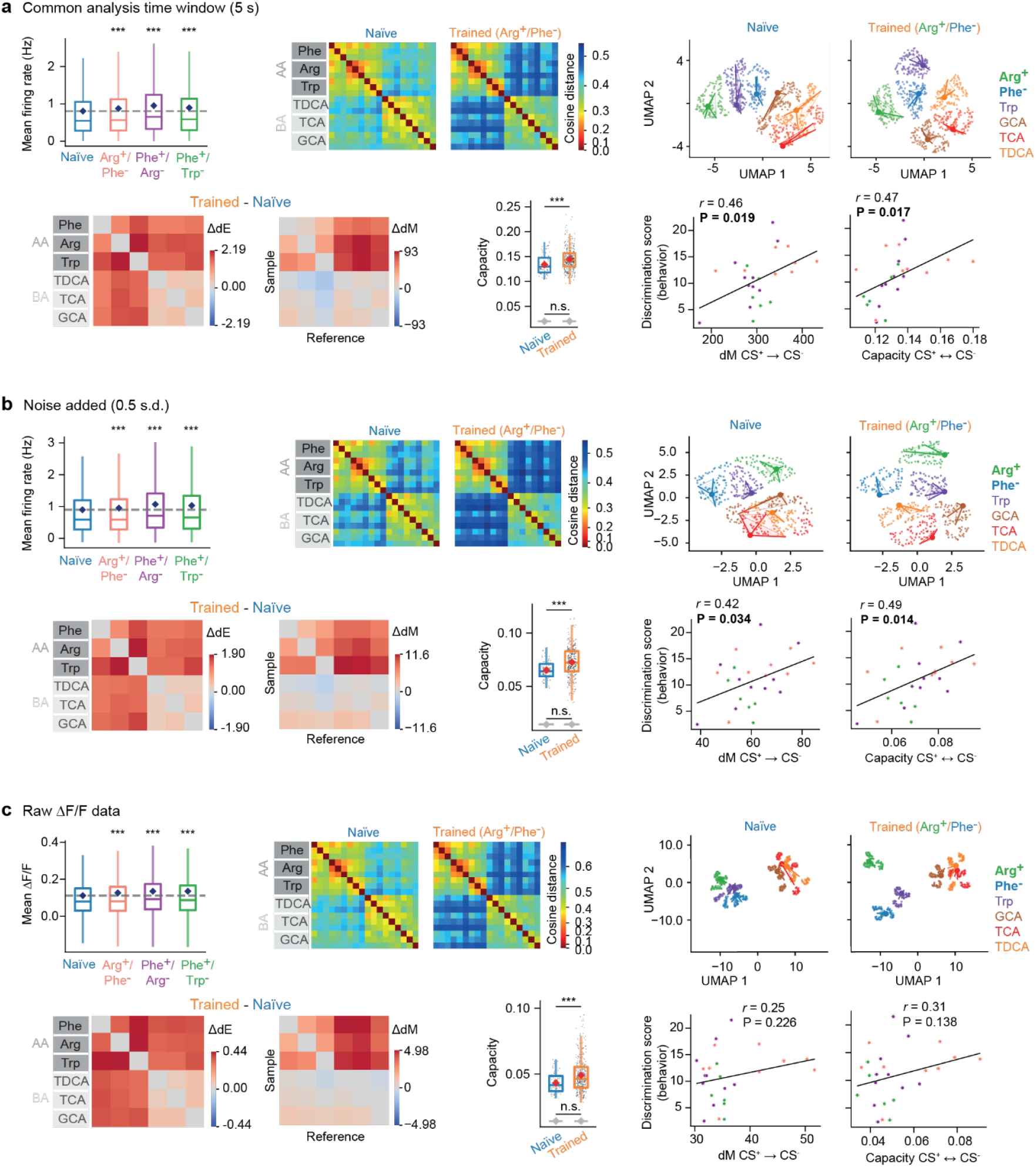
Variation of analysis parameters. **a**, Main results obtained with a common analysis time window (5 s starting 1 s after response onset; conventions as in the corresponding figures). Top row: mean responses per training group (c.f. Fig. 1h), mean cosine distances matrices in naïve and trained fish (c.f. Fig. 2a), projection of activity patterns into 2D UMAP space (same as Fig. 2d). Bottom row: mean difference matrices for dE and dM (trained – naïve; c.f. Fig. 4c,d), manifold capacity (c.f. Fig. 5b), behavioral odor discrimination score as a function of dM_CS+→CS-_ and manifold capacity (c.f. Fig. 6). **b**, Same analyses after adding noise to activity data (0.5 s.d.; default analysis time windows). **c**, Same analyses using raw ΔF/F as activity measurements (default analysis time windows). Note robustness of results against all parameter variations tested.

## Supplementary Note 1: Manifold Capacity and Effective Geometric Measures

Suppose we have simultaneous measurements of activity across many neurons from an animal presented with two stimulus conditions (e.g., two different odors). Imagine that there is some downstream neuron receiving signals from these recorded neurons: how can it tell the two stimulus conditions apart? Assuming this downstream neuron aggregates information by collecting upstream firing events weighted by the strength of synaptic connectivity, i.e., it essentially uses a weighted sum and then a threshold to determine whether the stimulus is odor A or odor B. Hence, we model such a downstream computation as a linear classification problem.

Before the animal learns to tell the difference between two distinct but similar odors, the two point clouds (or manifolds) of activity patterns could be very tangled with each other in the high-dimensional space where each coordinate corresponds to the activity level of a neuron. Such tangledness makes it hard for a downstream neuron to tell apart these two odors; in other words, it is hard to linearly classify the two odor manifolds. We asked whether the odor manifolds may become untangled and more separated from each other - and, thus, easier to discriminate for a downstream neuron - after the animal was trained in a task that required discrimination between two odors.

To formulate the above intuition into a rigorous quantitative analysis method, we need to answer two questions: (i) how to quantify the degree of separability of two odor manifolds, and (ii) how to connect the properties of the neural activity patterns - such as the firing statistics of neurons, the covariability between neurons, and the structure and shape of odor manifolds - to the degree of separability?

In our analysis, manifold capacity answers question (i) by quantifying the degree of separability of two odor manifolds. Effective geometric measures from GLUE answer question (ii) by rigorously linking the properties of the neural activity patterns to manifold capacity. Now, let’s dive into the formal definition of them.

### Manifold capacity

Suppose we have multiple datapoints representing simultaneous recordings across a population of neurons (e.g., at different time points or during different trials) from an animal presented with two stimulus conditions (e.g., two different odors). Concretely, let’s say we have N neurons recorded, and for each stimulus condition we have M datapoints. Namely, for each stimulus condition, there are M neural activity patterns in the form of N-dimensional vectors where each coordinate corresponds to the activity level of a neuron.

When the number of neurons N is very large (N ≫ M), these two point clouds (or manifolds) are very likely to be linearly separable from each other due to basic facts from linear algebra. However, when N becomes smaller and smaller (e.g., we randomly sample a subset of neurons), then the chance that the manifolds become linearly inseparable increases. Quantitatively, we can analyze the *manifold separability curve* p(N), defined as the probability of linear separability after subsampling the number of neurons to N (see Fig. 1a below). The area above the curve corresponds to the critical number of neurons needed for a downstream neuron to read out from for linear classification. We call this number *critical dimension N\**.

**Figure 1:**
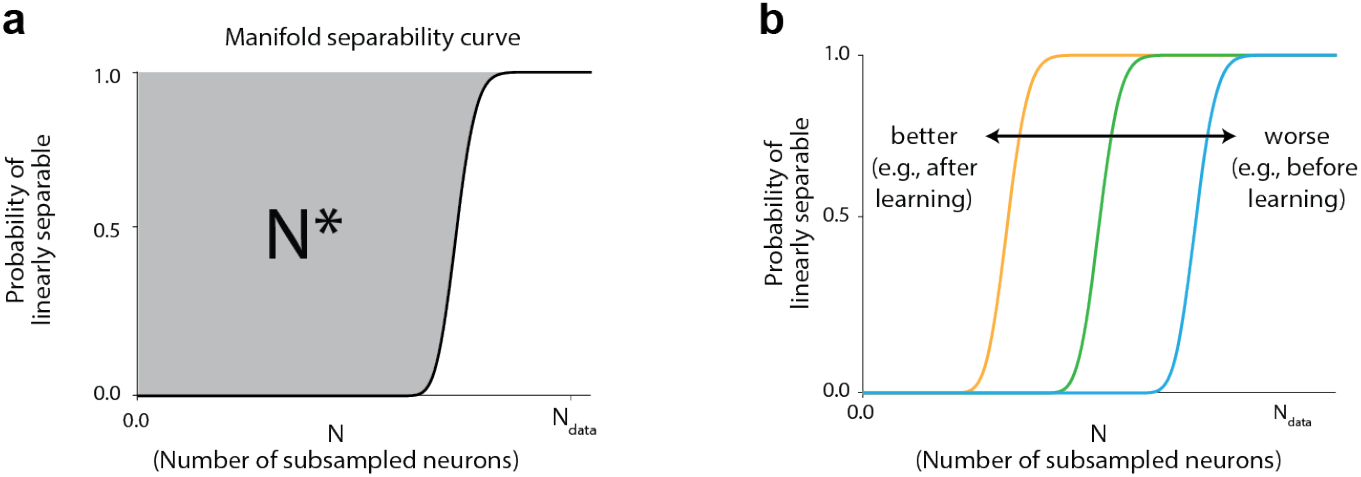
Manifold separability curve. Adapted from ref. 1 (Fig. S2 and S5b).

If the manifold separability curve moves from right to left (see Fig. 1b above), then a downstream neuron can tell the two odors apart using a smaller number of neurons. Namely, the odor manifolds become more separable, and this is quantified by N*. As a result, to evaluate if the separability of odor manifolds improved by training of an animal in a task, we can compare the N* between naïve animals and animals that have been trained. Indeed, what we see in our analysis is that N* is smaller in zebrafish that learned to discriminate the odors.

Manifold capacity α is simply a rescaling of critical dimension by the number of manifolds P (in our case here P = 2): α = P/N*. The larger the capacity is (smaller N*), the more separable the odor manifolds are.

Before moving on to effective geometric measures, we briefly compare manifold capacity to conventional approaches to assess separability. A traditional approach to quantify separability between two manifolds is based on support vector machine (SVM) theory: one identifies the linear classifier with the maximum margin, and the larger this margin, the greater the separability. In contrast, GLUE theory does not focus on the best possible classifier in the full neural space. Instead, it examines linear classifiers built from random sub-populations of neurons. The fewer neurons required for successful separation - the higher the manifold capacity - the more separable the manifolds are. Thus, while SVM captures the best case using all neurons, manifold capacity reflects the average performance across random subsets of neurons.

Finally, we note two key assumptions underlying the theory. First, each object manifold is modeled as the convex hull of its activity patterns. This follows from the fact that, for linear classification, considering the convex hull of points from the same class is equivalent to considering the original points themselves. Second, when analyzing subpopulations of neurons, we model them as random subsets of the full population. Beyond these, the theory does not impose additional assumptions on the structure of the data. Hence, unlike various alternative measures of separability, GLUE does not approximate manifolds by multivariate normal or other distributions.

### Effective geometric measures

The manifold separability curve can be analyzed mathematically, allowing manifold capacity α (and the critical dimension N*) to be estimated directly from a simple formula:

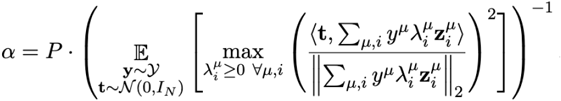

In the formula above, *P* is the number of manifolds and 𝔼 stands for taking the expectation. *y*^μ^ is a class label (*y*^1^ = 1 for odor 1 and *y*^2^ = -1 for odor 2) and the vector ***t*** represents a classifier drawn randomly from a multivariate normal distribution (i.e., a random direction in which population activity is projected). The expectation is taken by randomly sampling over *y* (from a set of label vectors *y*) and ***t*** (from the isotropic N-dimensional Gaussian distribution *N*(0, *I*_*N*_)). Inside the expectation is a term that involves maximizing a convex function. The variables 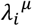 are non-negative weights, μ is the index for the manifold, *i* is the index for the point (i.e., datapoint number) in the μ-th manifold, and 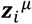 is an activity vector (i.e., neural population activity) for the *i*-th datapoint in the μ-th manifold. Hence, the term 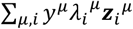 is a convex combination of the activity vectors weighted by the label. Thus, it is also an N-dimensional vector. Finally, in the numerator ⟨∙,∙ ⟩ stands for the inner product between two vectors, and in the denominator ‖∙ ‖_2_ stands for the L2 norm of a vector. In summary, the capacity formula computes the average (over random draws of *y* and ***t***) of the squared L2 norm of the projection of ***t*** onto the convex cone spanned by the activity vectors weighted by their manifold labels. We remark that the random sampling of ***t*** is done by random subsampling of neurons; as pointed out above, it does not imply anything about the structure of the data.

This formula has two key implications. First, it enables fast estimation of manifold capacity without explicitly computing the separability curve. Second, it provides an analytical link between manifold capacity and the underlying activity patterns 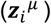, thereby revealing which structural properties of the manifolds contribute to separability. In what follows, we outline how the GLUE approach leverages this formula to derive effective geometric measures.

### 1. Anchor point distribution

When analyzing recordings of neural activity patterns with M datapoints per condition, a common assumption one would make is to treat every datapoint equally. Namely, for each condition, the uniform distribution over the M datapoints would be used to calculate properties such as mean response, covariance, etc.

**Figure 2:**
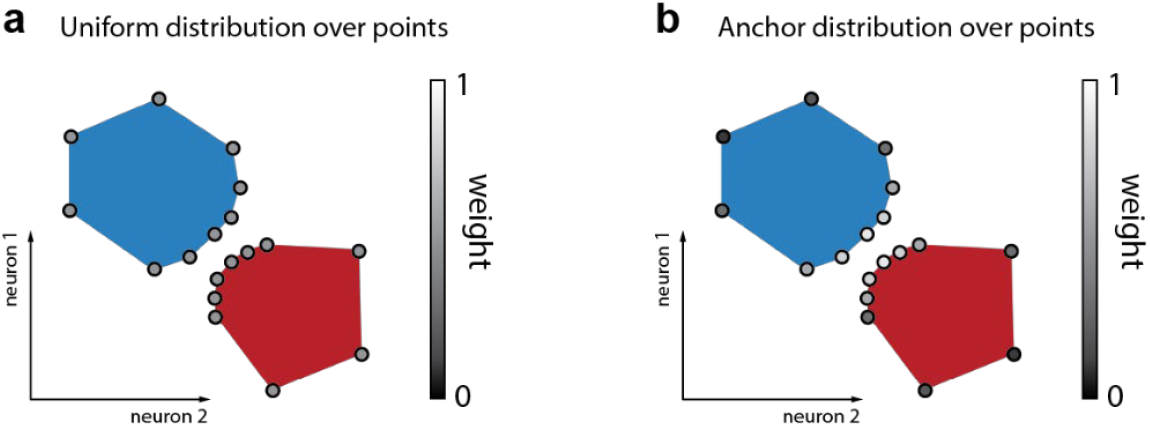
Schematic illustration of the difference between the uniform distribution over data point (**a**) and the anchor point distribution over data points (**b**).

In the GLUE approach, however, we do not draw datapoints randomly from a uniform distribution but choose them based on the manifold capacity formula. Concretely, for each sampled pair of (*y*, ***t***), the optimizer 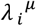 can be thought of as a function of (*y*, ***t***). By rescaling 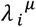 (*y*, ***t***) so that 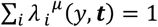 for each μ, the function 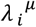 (*y*, ***t***) can be interpreted as a probability distribution over the M datapoints from condition μμ. This naturally leads to the definition of *anchor points*:

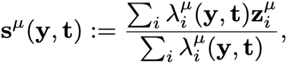

Intuitively, the anchor with respect to (*y*, ***t***) is the most relevant point in the μ-th manifold for the classification on top of a random sub-population of neurons. Furthermore, in GLUE, the capacity formula can be rewritten as

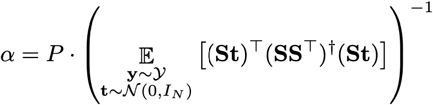

Where ***S*** is a P × N matrix with ***s*** ^μ^(*y*, ***t***) on its rows. To understand the above variation form of the capacity formula, recall that the previous form of the formula can be understood as the squared L2 norm of the projection of ***t*** onto the convex cone spanned by the activity vectors weighted by their manifold labels. Furthermore, the squared L2 norm term can be shown to be the squared L2 norm of the projection of vector ***t*** onto the subspace spanned by the anchor points. Through this observation, one can express the projection of ***t*** onto this subspace via the orthogonal projection operator ***S***^T^(***SS***^T^)^†^***SS. St*** is the projection of the random vector ***t*** onto the anchor points, ***SS***^T^is the covariance (Gram) matrix of the anchor points (representing the “geometry” of the anchor space), and † denotes the pseudoinverse. The projection operator therefore quantifies the squared length of the projection of ***t*** onto the anchor points, measured in the space spanned by the anchor points after correcting for their covariances.

To summarize, the manifold capacity is now analytically connected to a simple formula (i.e., the squared L2 norm of the projection of vector ***t*** to the subspace spanned by the anchor points, averaged with respect to the anchor point distribution).

### 2. Center-axis decomposition

Next, after replacing the uniform distribution over the datapoints from the same experimental condition with the anchor point distribution, we can conduct standard statistical analyses - such as estimating mean and variance of the responses - by averaging with respect to the anchor point distribution as opposed to the conventionally used uniform distribution. Conventionally, one would have defined the center of the μ-th manifold as the average of the activity patterns over all datapoints, i.e., 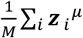. Now, it is natural to define the center of the μ-th manifold as the average of its anchor points, i.e., 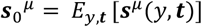. Furthermore, the axis part of an anchor point is then defined as 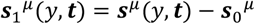.

Intuitively, the above *anchor center-axis decomposition* is analogous to what one would conventionally do by segregating the set of datapoints into components representing their mean and their variation. The main difference is that, traditionally, the mean is defined as the uniform average over the datapoints, while here the average is calculated with respect to the anchor point distribution. Likewise, the variation is determined here based on the anchor point distribution rather than the uniform distribution.

### 3. Effective manifold dimension and radius

After segregating the center and axis components of an anchor point we define *effective geometric measures* such as effective dimension and effective radius that are (i) analogous to the conventional measures for manifold dimension and radius using the uniform distribution and (ii) analytically linked to the capacity formula. In particular, in the original GLUE paper^1^, the authors showed that these effective geometric measures can faithfully track the ground truth dimension and radius of simulated examples when the underlying correlations between manifolds are mild. And when the correlations are strong, the effective geometric measures can more robustly track their effect on classification efficiency.

#### Formulas for effective manifold dimension and radius

Similar to the matrix ***S*** defined above, here we define matrix ***S***_0_ and ***S***_1_ as P × N matrices with 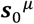 and 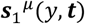 on their rows, respectively. In GLUE, the following three quantities were defined:

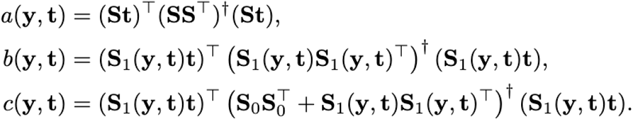

Here *a*(*y*, **t**) is the squared L2 norm of the projection of vector **t** onto the subspace spanned by the anchor points as we saw in the capacity formula. b(*y*, **t**) is an analogous projection term but replacing anchor points with their axial parts. c(*y*, **t**) is a correction term between *a* (*y*, **t**) and b(*y*, **t**) assuming no center-axis correlations (i.e., removing all cross terms that contain inner products (covariances) between the center part of an anchor point and the axis part of an anchor point).

Notice that the capacity α = *P*/*E*_*y*,**t**_[*a*(*y*, **t**)]. Now, the effective dimension and radius are defined as

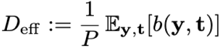

and

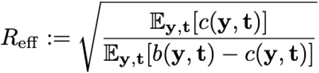

To build up intuitions for effective dimension and radius, it is helpful to first consider a scenario where there is no correlation between the anchor points across manifolds. Namely, the inner product between ***s***^μ^(*y*, **t**) and ***s***^*v*^(*y*, **t**) is zero when μ ≠ *v*. In this case, the above equations can be simplified:

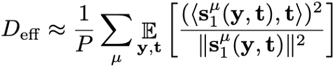

and

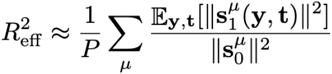

Now, both effective dimension and radius can be approximated by an average of some quantities about each individual manifold. Furthermore, if the manifolds are *D*-dimensional balls with radius (normalized by the length of the manifold center) being *R*, then the quantity on the right-hand side of the equations are exactly *D* and *R*^2^, respectively.

When there are correlations between manifolds, the effective dimension and radius can detect their effect on the separability (through the analytical connection to manifold capacity). Specifically, the GLUE approach shows that the effective dimension captures the correlation between manifolds’ axes (i.e., within-manifold variations) while the effective radius captures the correlations between manifolds’ centers (i.e., mean responses). See Figure S6 in GLUE paper^1^ for a quantitative example.

### 4. Anchor statistics

Finally, one can also utilize simple statistics from the anchor points to quantify structure in data, just like what is conventionally done by using mean firing rate and/or covariance etc. Some common statistical measures from anchor point distribution include:

- Anchor center alignment: For manifold μ and manifold *v*, the anchor center alignment between the two is:

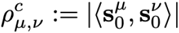

which captures the cosine similarity between the mean responses defined by weighting over the anchor points (as opposed to over the uniform distribution).
- Anchor axis alignment: For manifold μμ and manifold *vv*, the anchor axis alignment between the two is:

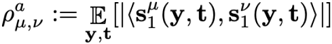

which captures the cosine similarity between the within-manifold variations defined by weighting over the anchor points (as opposed to over the uniform distribution).
- Anchor center-axis alignment: For manifold μμ and manifold *vv*, the anchor center alignment between the two is:

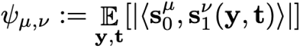

which captures the cosine similarity between the mean responses and within-manifold variations defined by weighting over the anchor points (as opposed to over the uniform distribution).

In all our analysis, we consider the average anchor statistical measures by taking the mean over all pairs of manifolds.

### Summary of effective geometric measures

To summarize, the GLUE theory establishes an analytical connection between separability and manifold geometry through a formula for manifold capacity. This formula naturally defines a non-uniform distribution over points on the manifolds, referred to as the anchor point distribution. Building on this, the theory introduces two classes of geometric measures:

1. **Effective dimension and effective radius**, which reduce to the conventional notions of dimension and radius when correlations are weak but also incorporate corrections that account for correlations relevant to downstream classification.
2. **Anchor statistics** - including center, axis, and center–axis alignment - which parallel conventional statistical measures but are computed with respect to the anchor point distribution rather than a uniform distribution, thereby capturing task-relevant structural properties of the manifolds.

**Figure 3:**
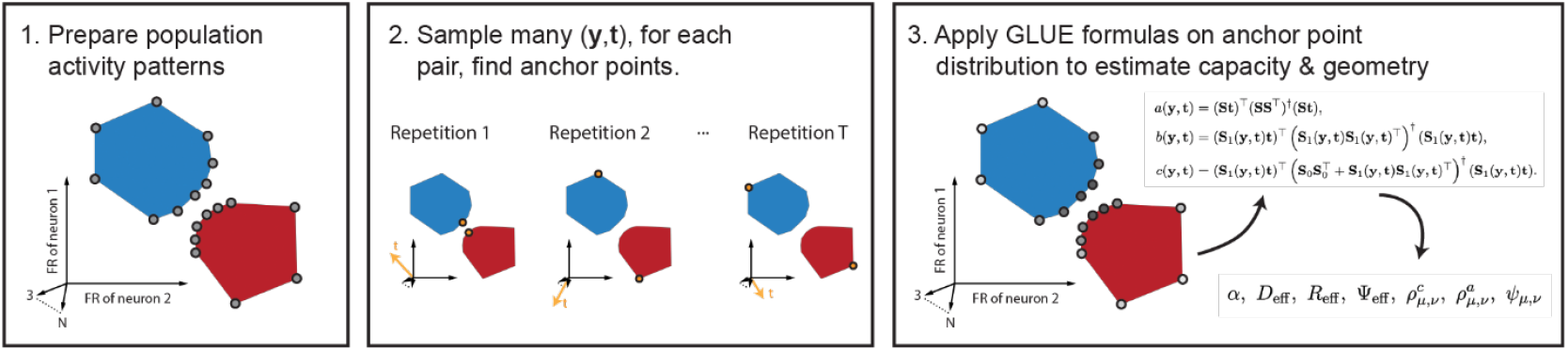
Schematic: Estimation of GLUE geometry from the anchor point distribution.

#### Difference Between GLUE Analysis & Conventional Methods

GLUE analysis provides two types of measurements for data analysis: (i) manifold capacity, which quantifies the degree of manifold separability, and (ii) task-relevant geometric measures, which quantify the contributions of structural properties of the manifolds to their separability. The key novelty of GLUE is that it establishes a rigorous mathematical link between these two levels of measurement, thereby connecting function (separability) and structure (geometry). Importantly, unlike many conventional analysis methods that rely on strong distributional assumptions (e.g., normality or Gaussian noise models), GLUE makes no such assumptions about the underlying data, making it broadly applicable to high-dimensional, complex neural representations.

To provide more intuition for GLUE analysis, we include below a comparison with conventional data analysis methods. Specifically, we compare GLUE analysis with (a) decoding analysis, (b) dimensionality reduction analysis (e.g., PCA, UMAP), and (c) spectral analysis (e.g., participation ratio).

### Comparison between GLUE and decoding analysis

A common approach in neural data analysis is decoding with linear classifiers. The most basic example is *Linear Discriminant Analysis (LDA)*, which assumes that neural responses for each class follow Gaussian distributions with shared covariance. LDA assesses separability by finding the projection that maximizes the ratio of between-class to within-class variance. While simple and interpretable, this approach depends strongly on normality assumptions and covariance estimation, which are often violated in real neural data.

An advanced alternative to LDA is the *Support Vector Machine (SVM)*, which does not rely on distributional assumptions and instead finds the separating hyperplane that maximizes the margin between classes. SVM provides a “best-case” estimate of separability given the full set of recorded neurons.

In contrast, GLUE is not tied to any particular classifier and makes no assumptions about data normality. Rather than evaluating separability once with a single optimal decision boundary, GLUE quantifies how reliably random subsets of neurons can achieve linear classification. The fewer neurons required for reliable separation, the higher the manifold capacity. This shift provides a population-level, assumption-free, and geometrically interpretable measure of separability that links *function* (classification capacity) with *structure* (manifold geometry).

### Comparison between GLUE and dimensionality reduction

Dimensionality reduction methods such as PCA or UMAP summarize neural activity by projecting it into a lower-dimensional space, inevitably discarding part of the original information. These projections are often useful for visualization or exploratory analysis but the resulting coordinates are not directly tied to variables of interest such as task performance. In contrast, GLUE defines its geometric measures - such as effective dimension, radius, and anchor statistics - directly in the original high-dimensional activity space. Rather than compressing the data, GLUE extracts task-relevant geometric structure without information loss from projection, ensuring that the measures remain fully connected to manifold separability.

### Comparison between GLUE and spectral analysis

Spectral analysis methods study the structure of neural activity by looking at the eigenvalue spectrum of the covariance matrix. For example, the participation ratio is a common metric that estimates the dimensionality of neural responses based on how variance is spread across eigenmodes. These measures provide a global picture of variability but rely on covariances computed with uniform weighting over all data points. In contrast, GLUE defines its effective geometric measures and anchor statistics using the anchor point distribution, which emerges directly from the manifold capacity formula. Because this distribution is task-relevant, the resulting measures are also task-relevant: they describe the geometry and statistics of manifolds in direct relation to their separability.

### Computational interpretation of manifold capacity: additional remarks

Unlike classical approaches, GLUE does not analyze optimal linear classification based on all neurons but asks how many randomly selected neurons are, on average, required to classify manifolds with a given probability. This approach more directly reflects the readout of information in biological networks where decoding neurons typically do not have access to all neurons in a network. Moreover, the resulting measure of separability *-* manifold capacity *-* differs from classical measures with respect to its computational interpretation. Manifold capacity is directly and analytically linked to computationally relevant quantities (see ref. 1 for mathematical description):

1. *Efficiency* (how many neurons are needed for the retrieval of information?). Higher manifold capacity means that the retrieval of information contained in neural manifolds requires access to fewer neurons in the network. Hence, less resources are required for readout. Moreover, finding a classifier (the learning process) becomes easier.
2. *Packing density* (how many manifolds can be fit into the activity space?). Higher manifold capacity means that more manifolds, each representing different information, can be packed into activity space, implying higher storage capacity.
3. *Robustness* (how much imperfection can be tolerated in the classifier?). Higher manifold capacity means that performance degrades more slowly when classifiers deviate from the optimum, implying that classification becomes more noise-resistant. Moreover, as it becomes easier to find classifiers with suboptimal yet acceptable performance, learning processes are facilitated and accelerated.

In summary, classical approaches such as SVMs lack biological plausibility and provide limited information on relevant features of neural representations such as efficiency and robustness because they focus on classification under optimal conditions. Hence, manifold capacity is a novel measure that provides additional computationally relevant information. From a neuroscience perspective, manifold capacity is of particular interest because it can be regarded as a measure that quantifies the “suitability” of neural manifolds for the representation of task-relevant information in biological (and artificial) networks.

#### A Simulated Toy Example for Intuition Building

Here we provide a toy example of synthetic Gaussian clouds, simulating representational odor manifolds, and examine how manifold capacity and geometric measures change with respect to the latent parameters in the simulated model. These examples were modified from the supplementary materials of the GLUE paper^1^.

Concretely, we consider *P* = 2 manifolds, each containing *M* points in an *N*-dimensional neural activity space. In all examples below we pick *M* = 200 and *N* = 1000, and we remark that for other choices of *M* and *N* the results would hold qualitatively. There are 4 parameters to determine the structure and correlations of manifolds:

- *d*: latent manifold dimension; we consider values ranging from 2 to 10.
- *r*: latent manifold radius; we consider values ranging from 0.3 to 2.
- 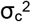: latent center correlations; we consider values ranging from 0 to 1.
- 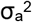 latent axis correlations; we consider values ranging from 0 to 1.

The following is the procedure to generate two synthetic Gaussian clouds from the above parameters:

1. Randomly sample 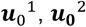 independently from N-dimensional isotropic Gaussian distribution with mean 0 and variance 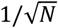 in each coordinate.
2. Correlate each pair of coordinates from 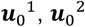 so that the covariance becomes 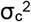.
3. Randomly sample 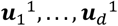 and 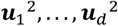 independently from N-dimensional isotropic Gaussian distribution with mean 0 and variance 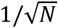 in each coordinate.
4. For each *i* = 1, …, *d*, correlate each pair of coordinates from 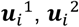 so that the covariance becomes 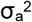.
5. Repeat the following for *M* times for each manifold (indexed by μ): sample (b_1_, b_2_, …, b_*d*_) from the d-dimensional unit sphere, and output 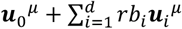.

Recall that GLUE analysis gives us the following measures: manifold capacity α, effective manifold dimension *D*_*eff*_, effective manifold radius *R*_*eff*_, anchor center alignment ρ_c_, anchor axis alignment ρ_a_, and anchor center-axis alignment ψ. In the following, we are going to see how different values of the latent parameters 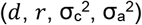 in the simulated model mapped to the GLUE measures (α, *D*_*eff*_, *R*_*eff*_, ρ_c_, ρ_a_, ψ).

**Figure 4:**
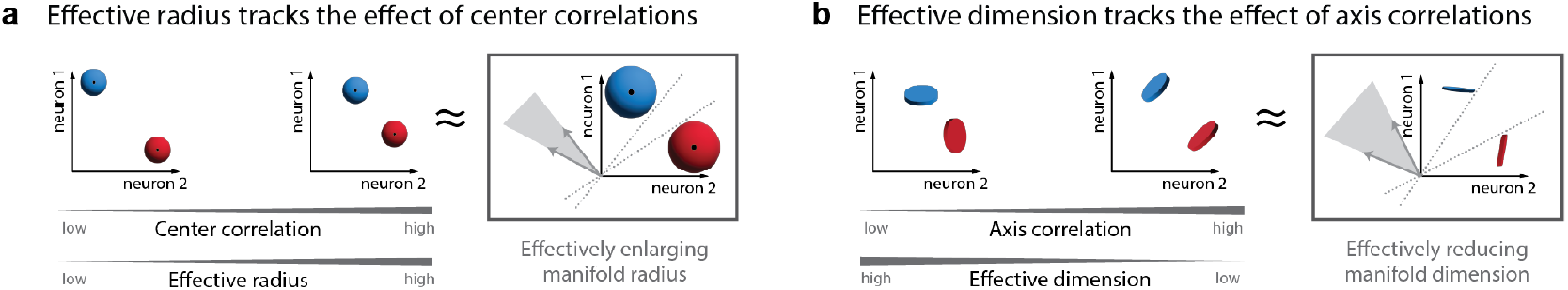
Schematic: connection between latent geometry and GLUE effective radius (**a**) and effective dimension (**b**).

### Case 1: No correlation between manifolds

Let us start with simple cases where 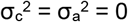. We swept through a grid of latent parameters with *d* in [2, 10] and *r* in [0.1, 2]. For each pair of (*d, r*), we ran 50 repetitions of random simulations and analyzed the resulting GLUE measures. To understand the results, first we plot all GLUE measures with respect to varying the latent manifold radius *r* as follows.

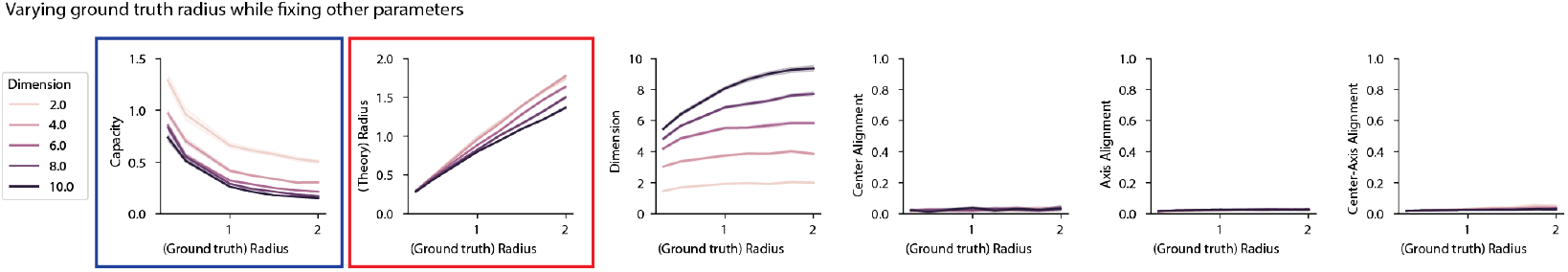

Notice that manifold capacity α decreases as the latent manifold radius *r* increases, agreeing with the intuition that manifolds become more tangled when *r* is larger. Moreover, the effective manifold radius *R*_*eff*_ faithfully tracks the value of latent manifold radius *r*, especially when the latent manifold dimension *d* is small (remark: the reason for the decrease in *R*_*eff*_ when *d* becomes large is a finite size effect: because *M* is small, the simulated procedure cannot sample the whole sphere when *d* is large).

Similarly, we plot all GLUE measures with respect to varying the latent manifold dimension *d* as follows.

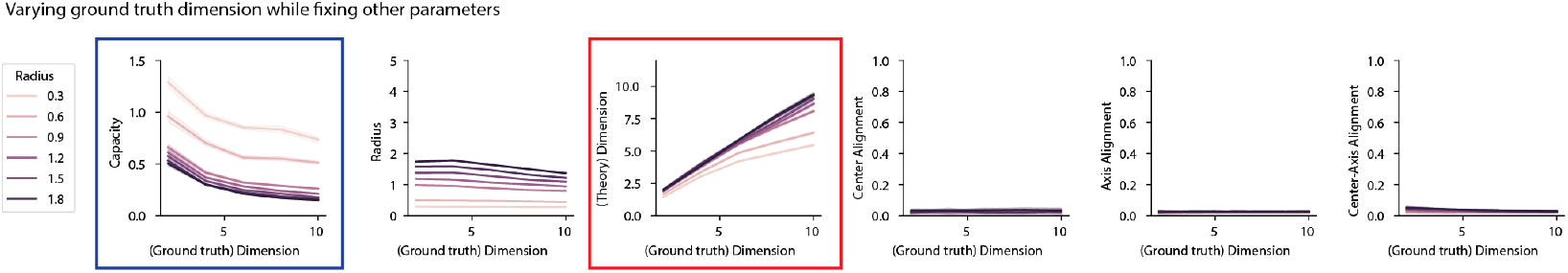

Notice that manifold capacity α decreases as the latent manifold radius *d* increases, agreeing with the intuition that manifolds become more tangled when *d* is larger. Moreover, the effective manifold dimension *D*_*eff*_ faithfully tracks the value of latent manifold dimension *d*, especially when the latent manifold dimension *r* is large (remark: the reason why *D*_*eff*_ decreases when *r* becomes small is that the point clouds become more point-like when the radius is small).

We conclude that in the regime when there are no correlations between the manifolds, effective dimension and radius (*D*_*eff*_, *R*_*eff*_) faithfully track the value of latent dimension and radius (*d, r*).

### Case 2: Introducing correlation between manifold centers

Next, we investigate the effect of correlation between manifold centers on capacity and GLUE geometry. We set *d* = 4, *r* = 1, and swept through a grid of latent parameters with 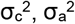 in [0, 1]. For each pair of 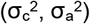, we ran 50 repetitions of random simulations and analyzed the resulting GLUE measures. To understand the results, first we plot all GLUE measures with respect to varying the latent manifold center correlations 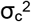 as follows.

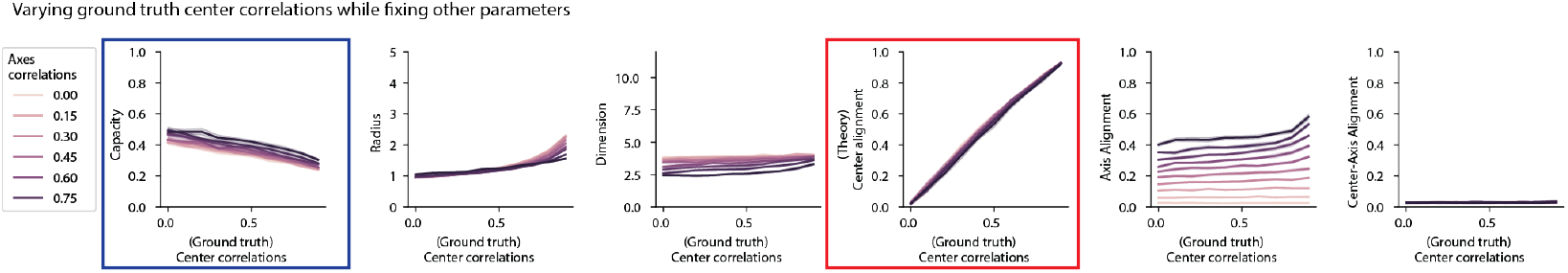

Notice that manifold capacity α decreases as the latent manifold center correlations 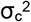 increase, agreeing with the intuition that manifolds become more tangled when they are closer to each other. Moreover, the anchor center alignment ρ_c_ faithfully tracks the value of latent manifold center correlations 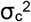. In addition, effective radius *R*_*eff*_ also tracks the effect of center correlations (as explained intuitively in Fig. 4a). Specifically, the larger the latent manifold center correlations 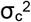, effectively the size of the manifolds becomes larger (hence decreasing capacity).

### Case 3: Introducing correlation between manifold axes

Finally, we investigate the effect of correlation between manifold axes on capacity and GLUE geometry. We set *d* = 4, *r* = 1, and swept through a grid of latent parameters with 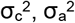 in [0, 1]. For each pair of 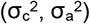, we ran 50 repetitions of random simulations and analyzed the resulting GLUE measures. To understand the results, first we plot all GLUE measures with respect to varying the latent manifold axis correlations 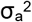 as follows.

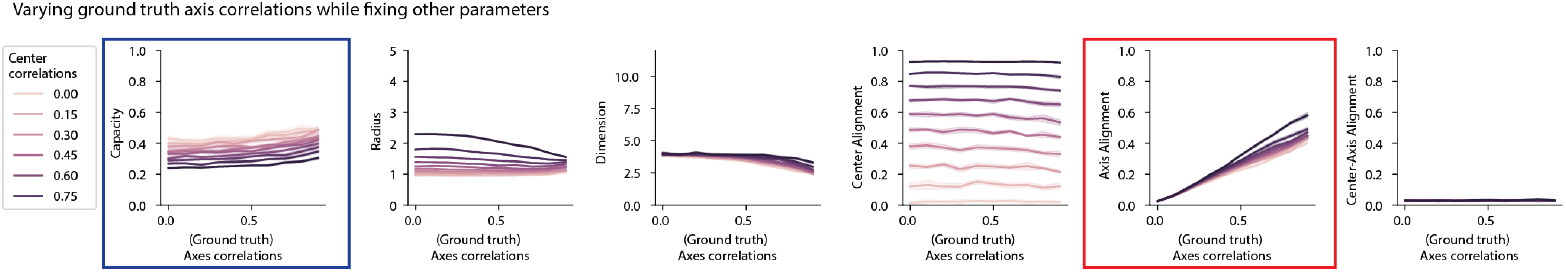

Notice that manifold capacity α increases as the latent manifold axis correlations 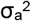 increase, agreeing with the intuition that manifolds become more untangled when they are internally more aligned with each other. Moreover, the anchor axis alignment ρ_a_ faithfully tracks the latent manifold axis correlations 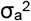. In addition, effective dimension *D*_*eff*_ also tracks the effect of axis correlations (as explained intuitively in Fig. 4b). Specifically, as latent manifold axis correlations 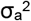 increase, effectively the manifolds become lower dimensional (hence increasing capacity).

## SUPPLEMENTARY NOTE 2

Detailed information on statistical analyses shown in figures:

**Fig. 1f**: Appetitive ζ scores were significantly higher for CS+ than CS-on all days (day 1, N = 25 fish, P = 6.0 × 10^-8^; day 2, N = 25, P = 1.8 × 10^-7^; day 3, N = 21, P = 6.7 × 10^-6^; day 4, N = 14, P = 1.2 × 10^-4^; Wilcoxon signed-rank test).

**Fig. 1h**: Activity was significantly higher in all trained groups (Arg^+^/Phe^-^: 0.95 ± 1.06 [mean ± SD], n_odor_ = 3, n = 13013 neurons from N = 9 fish; Phe^+^/Arg^-^: 1.07 ± 1.10, n_odor_ = 3, n = 13379 neurons from N = 10 fish); Phe^+^/Trp^-^: 1.03 ± 1.11, n_odor_ = 3, n = 7760 neurons from N = 6 fish) than in naïve fish (0.90 ± 0.93, n_odor_ = 3, n = 9059 neurons from N = 6 fish; Kruskal–Wallis test, n = 129633, P = 3.7 × 10^-143^). Posthoc Dunn’s test: Arg^+^/Phe^-^ vs naïve, P = 2.9 × 10^-10^; Phe^+^/Arg^-^ vs naïve, P = 6.1 × 10^-104^; Phe^+^/Trp^-^ vs naïve, P = 2.8 × 10^-45^.

**Fig. 2b**: Amino acids vs amino acids (AA-AA), naïve (mean ± SD): 0.336 ± 0.064, n = 18 odor pairs from N = 6 fish; trained: 0.354 ± 0.053, n = 75 odor pairs from N = 25 fish, Mann–Whitney U test, P = 0.107. Amino acids vs bile acids (AA-BA), naïve: 0.433 ± 0.058, n = 54 odor pairs from N = 6 fish; trained: 0.471 ± 0.057, n = 225 odor pairs from N = 25 fish, Mann–Whitney U test, P = 0.0002. Bile acids vs bile acids (BA-BA), naïve: 0.354 ± 0.055, n = 18 odor pairs from N = 6 fish; trained: 0.389 ± 0.054, n = 75 odor pairs from N = 25 fish, Mann–Whitney U test, P = 0.0007.

**Fig. 2c**: Amino acids, naïve (mean ± SD): 0.270 ± 0.076, n = 54 trial pairs from N = 6 fish; trained: 0.257 ± 0.069, n = 225 trial pairs from N = 25 fish, Mann–Whitney U test, P = 0.202. Bile acids, naïve: 0.318 ± 0.060, n = 54 trial pairs from N = 6 fish; trained: 0.329 ± 0.076, n = 225 trial pairs from N = 25 fish, Mann– Whitney U test, P = 0.264).

**Fig. 2e**: LDA cross-validation accuracy was significantly higher in all trained groups relative to the naïve group across all sample sizes. For 200 neurons, group differences were significant (Kruskal–Wallis test, n = 465 odor pairs from N = 31 fish, P = 3.7 × 10^-9^), with post-hoc Dunn tests showing significant differences between the naïve group and Arg^+^/Phe^−^ (P = 7.0 × 10^-8^), Phe^+^/Arg^−^ (P = 2.5 × 10^-6^), and Phe^+^/Trp^−^ (P = 0.018). For sample sizes of 300, 400, 500, 600, and 700 neurons, group differences remained significant (Kruskal–Wallis tests: P = 1.8 × 10^-8^, 1.8 × 10^-7^, 4.0 × 10^-7^, 1.0 × 10^-6^, and 4.9 × 10^-6^, respectively), and post-hoc comparisons against the naïve group were significant for Arg^+^/Phe^−^ (P = 3.1 × 10^-7^, 4.7 × 10^-6^, 2.0 × 10^-5^, 8.6 × 10^-5^, and 4.0 × 10^−4^), Phe^+^/Arg^−^ (P = 4.5 × 10^-6^, 1.3 × 10^-5^, 3.5 × 10^-5^, 8.8 × 10^-5^, and 5.0 × 10^−4^), and Phe^+^/Trp^−^ (P = 0.009, 0.008, 0.010, 0.008, and 0.015), respectively.

**Fig. 4e**: dE between manifolds representing two amino acids (AA↔AA), and between manifolds representing one amino acid and one bile acid (AA↔BA), was significantly higher in trained fish (AA↔AA: 8.05 ± 2.27, mean ± SD; n = 75 pairs from N = 25 fish; AA↔BA: 8.89 ± 2.50, n = 225 pairs from N = 25 fish) than in naïve fish (AA↔AA: 6.53 ± 1.26, n = 18 pairs from N = 6 fish, Mann–Whitney U test, P = 0.009; AA↔BA: 7.51 ± 1.76, n = 54 pairs from N = 6 fish, P = 0.0003). dE between manifolds representing two bile acids (BA↔BA) was not significantly different between trained (5.22 ± 1.42, n = 75 pairs from N = 25 fish) and naïve fish (4.93 ± 0.61, n = 18 pairs from N = 6 fish, Mann–Whitney U test, P = 0.253).

**Fig. 4f**: dM was significantly higher in trained fish between manifolds representing amino acids (AA→AA; trained: 341.3 ± 90.8, n = 150 pairs from N = 25 fish; naïve: 308.6 ± 67.4, n = 36 pairs from N = 6 fish, Mann–Whitney U test, P = 0.044) and in the direction from amino acids to bile acids (AA→BA; trained: 492.0 ± 118.1, n = 225 pairs from N = 25 fish; naïve: 414.3 ± 98.6, n = 54 pairs from N = 6 fish, Mann– Whitney U test, P = 3.1 × 10^-6^). No significant differences in dM between trained and naïve fish were observed in the direction from bile acids to amino acids (BA→AA; trained: 273.5 ± 53.0, n = 225 pairs from N = 25 fish; naïve: 280.3 ± 40.6, n = 54 pairs from N = 6 fish, Mann–Whitney U test, P = 0.657) and between manifolds representing bile acids (BA→BA; trained: 271.7 ± 50.4, n = 150 pairs from N = 25 fish; naïve: 264.3 ± 24.5, n = 36 pairs from N = 6 fish, Mann–Whitney U test, P = 0.272).

**Fig. 5b**: Left: Naïve (mean ± SD): 0.100 ± 0.011, n = 90 odor pairs from N = 6 fish; trained: 0.108 ± 0.014, n = 375 odor pairs from N = 25 fish (Mann–Whitney U test, P = 2.8 × 10^-7^). Shuffled naïve: 0.014 ± 0.0001; shuffled trained: 0.014 ± 0.0001 (Mann–Whitney U test, P = 0.851). Right: Manifold capacity was significantly increased in all trained groups relative to the naïve group. Arg^+^/Phe^-^ (mean ± SD): 0.112 ± 0.019, n = 135 odor pairs from N = 9 fish; Phe^+^/Arg^-^: 0.106 ± 0.010, n = 150 odor pairs from N = 10 fish; Phe^+^/Trp^-^: 0.105 ± 0.010, n = 90 odor pairs from N = 6 fish. Differences in manifold capacity were statistically significant between the naïve group (0.100 ± 0.011, n = 90 odor pairs from N = 6 fish) and each of the three training groups (Kruskal–Wallis test, n = 465, d.f. = 3, H = 31.08, P = 8.2 × 10^-7^; post-hoc Dunn test: Arg^+^/Phe^-^, P = 1.2 × 10^-7^; Phe^+^/Arg^-^, P = 5.6 × 10^-5^; Phe+/Trp^-^, P = 0.002). Gray: manifold capacity after shuffling manifold labels was not significantly different between the naïve and trained groups (Kruskal–Wallis test, P = 0.965).

**Fig. 5c**: Amino acids versus amino acids (AA-AA; mean ± SD), naïve: 0.094 ± 0.006, n = 18 pairs from N = 6 fish; trained: 0.103 ± 0.010, n = 75 pairs from N = 25 fish, Mann–Whitney U test, P = 7.4 × 10^-5^. Amino acids versus bile acids (AA-BA), naïve: 0.105 ± 0.011, n = 54 pairs from N = 6 fish; trained: 0.114 ± 0.013, n = 225 pairs from N = 25 fish, Mann–Whitney U test, P = 1.7 × 10^-5^. Bile acids versus bile acids (BA-BA), naïve: 0.089 ± 0.005, n = 18 pairs from N = 6 fish; trained: 0.094 ± 0.008, n = 75 pairs from N = 25 fish, Mann–Whitney U test, P = 0.009.

**Fig. 5e**: Effective radius in the trained group (mean ± SD 0.953 ± 0.041, n = 375 pairs from N = 25 fish) was significantly lower than in the naïve group (0.986 ± 0.039, n = 90 pairs from N = 6 fish; Mann– Whitney U test, P = 4.0 × 10^-10^). Effective dimension in the trained group (17.0 ± 1.9) was significantly lower than in the naïve group (17.7 ± 1.6; Mann–Whitney U test, P = 0.0007). Effective center alignment in the trained group (0.808 ± 0.049) was significantly lower than in the naïve group (0.835 ± 0.044; Mann–Whitney U test, P = 5.0 × 10^-5^). Effective axes alignment in the trained group (0.099 ± 0.028) was significantly lower than in the naïve group (0.109 ± 0.028; Mann–Whitney U test, P = 0.006).

**Extended Data Fig. 2d**: Cross-validation accuracy of pairwise odor classification by a support vector machine was significantly higher in trained groups than in the naïve group across all sample sizes. For 200 neurons, group differences were significant (Kruskal–Wallis test, n = 465 odor pairs from N = 31 fish, P = 2.9 × 10^-9^), with post-hoc Dunn tests showing significant differences between the naïve group and Arg^+^/Phe^−^ (P = 1.5 × 10^−6^), Phe^+^/Arg^−^ (P = 4.0 × 10^−7^), but not Phe^+^/Trp^−^ (P = 0.264). For sample sizes of 300, 400, 500, 600, and 700 neurons, group differences remained significant (Kruskal–Wallis tests: P = 4.6 × 10^−9^, 1.6 × 10^−8^, 1.5 × 10^−8^, 2.0 × 10^−8^, and 3.7 × 10^−8^, respectively), and post-hoc comparisons against the naïve group remained significant for Arg^+^/Phe^−^ (P = 3.8 × 10^−6^, 7.8 × 10^-6^, 2.4 × 10^-5^, 4.8 × 10^-5^, and 8.0 × 10^−5^), Phe^+^/Arg^−^ (P = 8.4 × 10^−7^, 1.1 × 10^−6^, 1.1 × 10^−6^, 9.1 × 10^−7^, and 1.5 × 10^−6^), but not Phe^+^/Trp^−^ (P = 0.428, 0.284, 0.337, 0.261, and 0.275).

**Extended Data Fig. 6c**: After shuffling of manifold labels (juvenile fish), dE between manifolds representing two amino acids (AA↔AA), one amino acid and one bile acid (AA↔BA) and two bile acids (BA↔BA) were not significantly different in naïve (AA↔AA: 1.7 ± 0.2, mean ± SD, AA↔BA: 1.6 ± 0.2, BA↔BA: 1.3 ± 0.2) and trained (AA↔AA: 1.8 ± 0.4, Mann-Whitney U test, P = 0.20, AA↔BA: 1.6 ± 0.4, P = 0.52, BA↔BA: 1.3 ± 0.3, P = 0.45).

**Extended Data Fig. 6f**: After shuffling of manifold labels (juvenile fish), dM from an amino acid manifold to another amino acid manifold (AA→AA), from one bile acid manifold to another bile acid manifold (BA→BA), from one amino acid manifold to a bile acid manifold (AA→BA) and from a bile acid manifold to an amino acid manifold (BA→AA) in naïve (AA→AA: 39.2 ± 1.5, mean ± SD, AA→BA: 38.5 ± 1.5, BA→AA: 38.6 ± 1.5, BA→BA: 36.7 ± 1.5) and trained (AA→AA: 39.6 ± 2.3, Mann-Whitney U test, P = 0.19, AA→BA: 39.0 ± 2.3, P = 0.06, BA→AA: 39.0 ± 2.3, P = 0.19, BA→BA: 36.8 ± 2.0, P = 0.62) were not significantly different.

**Extended Data Fig. 6i**: After shuffling (adult fish), dE between CS+ and CS-(CS↔CS), between conditioned and non-conditioned (CS↔nonCS), and between non-conditioned odors (nonCS↔nonCS) were not significantly different in naïve (CS↔CS: 0.76 ± 0.25, mean ± SD, CS↔nonCS: 0.76 ± 0.20, nonCS↔nonCS: 0.77 ± 0.20) and trained (CS↔CS: 0.92 ± 0.18, Mann-Whitney U test, P = 0.24, CS↔nonCS: 0.87 ± 0.19, P = 0.07, nonCS↔nonCS: 0.83 ± 0.18, P = 0.67). Pale boxes show dE before shuffling, same as in Fig. 8h.

**Extended Data Fig. 6**l: After shuffling (adult fish), dM between CS+ and CS-(CS↔CS), from conditioned to non-conditioned (CS↔nonCS), from non-conditioned to conditioned (nonCS↔CS), and between non-conditioned odors (nonCS↔nonCS) were significantly higher in trained (CS↔CS: 53.3 ± 3.8, mean ± SD, CS↔nonCS: 54.1 ± 5.4, nonCS↔CS: 54.2 ± 6.0, nonCS↔nonCS: 54.9 ± 6.0) than in naïve (CS↔CS: 46.1 ± 9.5, Mann-Whitney U test, P = 0.018, CS↔nonCS: 47.1 ± 9.3, P = 0.003, nonCS↔CS: 46.9 ± 9.0, P = 0.003, nonCS↔nonCS: 48.3 ± 9.3, P = 0.022) fish.

**Extended Data Fig. 7a**: Effective radius: amino acids vs amino acids (AA-AA) [naïve] 1.01 ± 0.02 (mean ± SD), n = 18 pairs from N = 6 fish, [trained] mean ± SD 0.98 ± 0.03, n = 75 odor pairs from N = 25 fish, Mann–Whitney U test, P = 9.4 × 10^-5^. Amino acids vs bile acids (AA-BA) [naïve] 0.97 ± 0.03, n = 54 pairs from N = 6 fish, [trained] 0.93 ± 0.03, n = 225 pairs from N = 25 fish, Mann–Whitney U test, P = 7.8 × 10^-8^. Bile acids vs bile acids (BA-BA) [naïve] 1.02 ± 0.03, n = 18 pairs from N = 6 fish, [trained] 0.99 ± 0.03, n = 75 pairs from N = 25 fish, Mann–Whitney U test, P = 0.0005. Effective dimension: AA-AA [naïve] 18.1 ± 1.1, [trained] 17.1 ± 1.5, P = 0.002. AA-BA [naïve] 17.0 ± 1.5, [trained] 16.3 ± 1.6, P = 0.006. BA-BA [naïve] 19.3 ± 0.9, [trained] 19.1 ± 1.5, P = 0.563. Effective center alignment: AA-AA [naïve] 0.86 ± 0.03, [trained] 0.84 ± 0.03, P = 0.015. AA-BA [naïve] 0.81 ± 0.04, [trained] 0.78 ± 0.04, P = 0.0002. BA-BA [naïve] 0.87 ± 0.03, [trained] 0.85 ± 0.03, P = 0.006. Effective axis alignment: AA-AA [naïve] mean ± SD 0.128 ± 0.033, [trained] 0.120 ± 0.022, P = 0.550. AA-BA [naïve] 0.098 ± 0.020, [trained] 0.089 ± 0.024, Mann–Whitney U test, P = 0.005. BA-BA [naïve] 0.121 ± 0.032, [trained] 0.106 ± 0.031, Mann–Whitney U test, P = 0.061.

**Extended Data Fig. 7b**: Percentage of capacity increase compared to the naïve group. Arg^+^/Phe^−^: Phe vs Arg: 11.55 ± 14.91, n = 9, P = 0.145. Phe vs Trp: 12.72 ± 13.96, n = 9, P = 0.050. Arg vs Trp: 11.40 ± 15.81, n = 9, P = 0.145. Phe^+^/Arg^−^: Phe vs Arg: 6.66 ± 7.32, n = 10, P = 0.118. Phe vs Trp: 9.87 ± 6.13, n = 10, P = 0.007. Arg vs Trp: 10.19 ± 5.49, n = 10, P = 0.005. Phe^+^/Trp^−^: Phe vs Arg: 11.30 ± 5.23, n = 6, P = 0.015. Phe vs Trp: 4.44 ± 5.59, n = 6, P = 0.180. Arg vs Trp: 8.99 ± 7.61, n = 6, P = 0.065.

**Extended Data Fig. 7c**: For a sample size of 200 neurons, group differences were significant (Kruskal– Wallis test, n = 465 odor pairs from N = 31 fish, P = 6.5 × 10^-6^), with post-hoc Dunn tests indicating higher manifold capacity relative to the naïve group for Arg^+^/Phe^−^ (P = 9.2 × 10^-7^), Phe^+^/Arg^−^ (P = 0.010), and Phe^+^/Trp^−^ (P = 0.049). For sample sizes of 300, 400, 500, 600, and 700 neurons, group differences remained significant (Kruskal–Wallis tests: P = 7.8 × 10^-6^, 4.6 × 10^-6^, 5.4 × 10^-6^, 5.5 × 10^-6^, and 3.0 × 10^-6^, respectively). Post-hoc comparisons against the naïve group were consistently significant for Arg^+^/Phe^−^ (P = 1.7 × 10^-6^, 8.5 × 10^-7^, 9.1 × 10^-7^, 9.5 × 10^-7^, and 5.4 × 10^-7^) and Phe^+^/Arg^−^ (P = 0.012, 0.011, 0.010, 0.013, and 0.009), whereas differences for Phe^+^/Trp^−^ were marginal or weakly significant (P = 0.054, 0.059, 0.051, 0.057, and 0.049)

**Extended Data Fig. 8**: “Naïve” (random; mean ± SD): 0.066 ± 0.005, n = 120 odor pairs from N = 8 network configurations; “Trained” (with assemblies): 0.071 ± 0.009, n = 120 odor pairs from N = 8 network configurations (Mann–Whitney U test, P = 1.5 × 10^-10^). Shuffled “naïve”: 0.040 ± 0.0001; shuffled “trained”: 0.040 ± 0.0001 (Mann–Whitney U test, P = 0.22). Effective radius was significantly lower for “trained” (mean ± SD 1.11 ± 0.05, n = 120 pairs from N = 8 network configurations) than for “naïve” networks (1.11 ± 0.05, n = 120 pairs from N = 8 network configurations; Mann–Whitney U test, P = 0.0002). Effective dimension was significantly lower for “trained” (22.9 ± 2.5) than for “naïve” networks (24.4 ± 1.5; Mann–Whitney U test, P = 2.2 × 10^-11^). Effective center alignment was not significantly different (“trained”: 0.45 ± 0.11; “naïve”: 0.46 ± 0.10 Mann–Whitney U test, P = 0.084). Effective axis alignment was not significantly different (“trained”: 0.048 ± 0.016; “naïve”: 0.046 ± 0.011; Mann–Whitney U test, P = 0.60).

**Extended Data Fig. 9b**: Conditioned odors [mean ± SD], trained: 0.23 ± 0.55, n_odor_ = 2, n = 6092 neurons from N = 9 fish; naïve: 0.18 ± 0.49, n_odor_ = 2, n = 5164 neurons from N = 8 fish; Mann–Whitney U test, P = 7.6 × 10^-88^; non-conditioned odors, trained: 0.22 ± 0.52, n_odor_ = 2, n = 6092 neurons from N = 9 fish; naïve: 0.19 ± 0.49, n_odor_ = 2, n = 5164 neurons from N = 8 fish; Mann–Whitney U test, P = 4.1 × 10^-66^.

**Extended Data Fig. 9f**: dE between CS+ and CS-(CS↔CS) was significantly higher in trained (mean ± SD: 3.72 ± 0.98, n = 9 pairs from N = 9 fish) than in naïve fish (2.46 ± 0.98, n = 8 pairs from N = 8 fish, Mann– Whitney U test, P = 0.046). dE between conditioned and non-conditioned (CS↔nonCS) was also significantly higher in trained (3.04 ± 1.03, n = 36 pairs from N = 9 fish) than in naïve fish (2.32 ± 0.86, n = 32 pairs from N = 8 fish, Mann–Whitney U test, P = 0.014). dE between non-conditioned odors (nonCS↔nonCS) was not significantly different (trained: 2.32 ± 0.66, n = 9 pairs from N = 9 fish; naïve: 2.01 ± 0.70, n = 8 pairs from N = 8 fish, Mann–Whitney U test, P = 0.423).

**Extended Data Fig. 9h**: dM between conditioned odors (CS→CS) was higher in trained (247 ± 99, n = 18 pairs, from N = 9 fish) than in naïve (170 ± 52, n = 16 pairs, from N = 8 fish, Mann–Whitney U test, P = 0.012). dM from conditioned to non-conditioned odors (CS→nonCS) was higher in trained (237 ± 85, n = 36 pairs from N = 9 fish) than in naïve (166 ± 55, n = 32 pairs from N = 8 fish, Mann–Whitney U test, P = 2×10^-4^). dM from non-conditioned to conditioned odors (nonCS→CS) was not significantly different (trained: 202 ± 64, n = 36 pairs from N = 9 fish; naïve: 173 ± 63, n = 32 pairs from N = 8 fish, Mann– Whitney U test, P = 0.072). dM from non-conditioned to non-conditioned odors (nonCS→nonCS) was not significantly different (trained: 194 ± 58, n = 18 pairs from N = 9 fish; naïve: 158 ± 52, n = 16 pairs from N = 8 fish, Mann–Whitney U test, P = 0.060).

**Extended Data Fig. 9i**: Capacity in trained fish (mean ± SD 0.081 ± 0.020, n = 54 pairs from N = 9 fish) was higher than in naïve fish (0.070 ± 0.017, n = 48 pairs from N = 8 fish; Mann–Whitney U test, P = 0.004). Differences after shuffling of labels were not significant (trained: 0.025 ± 0.0002; naïve: 0.025 ± 0.0002; Mann–Whitney U test, P = 0.8).

**Extended Data Fig. 9j**: Effective radius was significantly lower in trained (1.25 ± 0.11, n = 54 pairs from N = 9 fish) than in naïve fish (1.33 ± 0.10, n = 48 pairs from N = 8 fish; Mann–Whitney U test, P = 0.0006). Effective dimension was significantly lower in trained (17.8 ± 3.0) than in naïve fish (19.4 ± 3.7; Mann– Whitney U test, P = 0.032). Effective center alignment was lower in trained (0.89 ± 0.077) than in naïve fish (0.92 ± 0.051); Mann–Whitney U test, P = 0.013). Effective axis alignment was significantly lower in trained (0.207 ± 0.062) than in naïve fish (0.252 ± 0.084; Mann–Whitney U test, P = 0.004).

## REFERENCES

1 Nieh, E. H. et al. Geometry of abstract learned knowledge in the hippocampus. Nature 595, 80–84 (2021).

2 Chung, S. & Abbott, L. F. Neural population geometry: An approach for understanding biological and artificial neural networks. Curr Opin Neurobiol 70, 137–144 (2021).

3 Hopfield, J. J. Neural networks and physical systems with emergent collective computational abilities. Proc. Natl. Acad. Sci. U. S. A. 79, 2554–2558. (1982).

4 Khona, M. & Fiete, I. R. Attractor and integrator networks in the brain. Nat Rev Neurosci 23, 744–766 (2022).

5 Kohonen, T. Self-organization and associative memory. (Springer, 1984).

6 Pereira, U. & Brunel, N. Attractor Dynamics in Networks with Learning Rules Inferred from In Vivo Data. Neuron 99, 227–238 e224 (2018).

7 Aljadeff, J., Gillett, M., Pereira Obilinovic, U. & Brunel, N. From synapse to network: models of information storage and retrieval in neural circuits. Curr Opin Neurobiol 70, 24–33 (2021).

8 Burak, Y. & Fiete, I. R. Accurate path integration in continuous attractor network models of grid cells. PLoS computational biology 5, e1000291 (2009).

9 Morcos, A. S. & Harvey, C. D. History-dependent variability in population dynamics during evidence accumulation in cortex. Nat Neurosci 19, 1672–1681 (2016).

10 Langdon, C., Genkin, M. & Engel, T. A. A unifying perspective on neural manifolds and circuits for cognition. Nat Rev Neurosci 24, 363–377 (2023).

11 Gallego, J. A., Perich, M. G., Miller, L. E. & Solla, S. A. Neural Manifolds for the Control of Movement. Neuron 94, 978–984 (2017).

12 Cimesa, L., Ciric, L. & Ostojic, S. Geometry of population activity in spiking networks with low-rank structure. PLoS computational biology 19, e1011315 (2023).

13 Mastrogiuseppe, F. & Ostojic, S. Linking Connectivity, Dynamics, and Computations in Low-Rank Recurrent Neural Networks. Neuron 99, 609–623 e629 (2018).

14 Cohen, U., Chung, S., Lee, D. D. & Sompolinsky, H. Separability and geometry of object manifolds in deep neural networks. Nat Commun 11, 746 (2020).

15 Chung, S., Lee, D. D. & Sompolinsky, H. Classification and Geometry of General Perceptual Manifolds. Physical Review X 8, 031003 (2018).

16 Neville, K. R. & Haberly, L. B. in The synaptic organization of the brain (ed G. M. Shepherd) 415–454 (Oxford University Press, 2004).

17 Hasselmo, M. E. & Barkai, E. Cholinergic modulation of activity-dependent synaptic plasticity in the piriform cortex and associative memory function in a network biophysical simulation. J Neurosci 15, 6592–6604 (1995).

18 Franks, K. M. et al. Recurrent circuitry dynamically shapes the activation of piriform cortex. Neuron 72, 49–56 (2011).

19 Haberly, L. B. Parallel-distributed processing in olfactory cortex: new insights from morphological and physiological analysis of neuronal circuitry. Chem. Senses 26, 551–576. (2001).

20 Wilson, D. A. & Sullivan, R. M. Cortical processing of odor objects. Neuron 72, 506–519 (2011).

21 Bolding, K. A. & Franks, K. M. Recurrent cortical circuits implement concentration-invariant odor coding. Science 361 (2018).

22 Miura, K., Mainen, Z. F. & Uchida, N. Odor representations in olfactory cortex: distributed rate coding and decorrelated population activity. Neuron 74, 1087–1098 (2012).

23 Rennaker, R. L., Chen, C. F., Ruyle, A. M., Sloan, A. M. & Wilson, D. A. Spatial and temporal distribution of odorant-evoked activity in the piriform cortex. J. Neurosci. 27, 1534–1542 (2007).

24 Iurilli, G. & Datta, S. R. Population Coding in an Innately Relevant Olfactory Area. Neuron 93, 1180–1197 e1187 (2017).

25 Pashkovski, S. L. et al. Structure and flexibility in cortical representations of odour space. Nature 583, 253–258 (2020).

26 Roland, B., Deneux, T., Franks, K. M., Bathellier, B. & Fleischmann, A. Odor identity coding by distributed ensembles of neurons in the mouse olfactory cortex. Elife 6 (2017).

27 Poo, C., Agarwal, G., Bonacchi, N. & Mainen, Z. F. Spatial maps in piriform cortex during olfactory navigation. Nature 601, 595–599 (2022).

28 Calu, D. J., Roesch, M. R., Stalnaker, T. A. & Schoenbaum, G. Associative encoding in posterior piriform cortex during odor discrimination and reversal learning. Cereb Cortex 17, 1342–1349 (2007).

29 Chapuis, J. & Wilson, D. A. Bidirectional plasticity of cortical pattern recognition and behavioral sensory acuity. Nat Neurosci 15, 155–161 (2011).

30 Schoonover, C. E., Ohashi, S. N., Axel, R. & Fink, A. J. P. Representational drift in primary olfactory cortex. Nature 594, 541–546 (2021).

31 Fink, A. J. P. et al. Experience-dependent reorganization of inhibitory neuron synaptic connectivity. biorxiv 10.1101/2025.01.16.633450 (2025).

32 Meissner-Bernard, C., Zenke, F. & Friedrich, R. W. Geometry and dynamics of representations in a precisely balanced memory network related to olfactory cortex. eLife 13, RP96303 (2025).

33 Mueller, T., Dong, Z., Berberoglu, M. A. & Guo, S. The dorsal pallium in zebrafish, Danio rerio (Cyprinidae, Teleostei). Brain Res 1381, 95–105 (2011).

34 Friedrich, R. W. Neuronal computations in the olfactory system of zebrafish. Annu Rev Neurosci 36, 383–402 (2013).

35 Hennequin, G., Agnes, E. J. & Vogels, T. P. Inhibitory Plasticity: Balance, Control, and Codependence. Annu Rev Neurosci 40, 557–579 (2017).

36 Renart, A. et al. The asynchronous state in cortical circuits. Science 327, 587–590 (2010).

37 Chou, C.-N. et al. Neural Manifold Capacity Captures Representation Geometry, Correlations, and Task-Efficiency Across Species and Behaviors. bioRxiv, 2024.2002.2026.582157 (2024).

38 Wakhloo, A. J., Sussman, T. J. & Chung, S. Linear Classification of Neural Manifolds with Correlated Variability. Physical Review Letters 131, 027301 (2023).

39 Yaksi, E., von Saint Paul, F., Niessing, J., Bundschuh, S. T. & Friedrich, R. W. Transformation of odor representations in target areas of the olfactory bulb. Nature Neurosci. 12, 474–482 (2009).

40 Blumhagen, F. et al. Neuronal filtering of multiplexed odour representations. Nature 479, 493–498 (2011).

41 Jacobson, G. A., Rupprecht, P. & Friedrich, R. W. Experience-Dependent Plasticity of Odor Representations in the Telencephalon of Zebrafish. Curr Biol 28, 1–14 e13 (2018).

42 Rupprecht, P. & Friedrich, R. W. Precise Synaptic Balance in the Zebrafish Homolog of Olfactory Cortex. Neuron 100, 669–683 e665 (2018).

43 Rupprecht, P. et al. A database and deep learning toolbox for noise-optimized, generalized spike inference from calcium imaging. Nat Neurosci 24, 1324–1337 (2021).

44 Meissner-Bernard, C. et al. Computational functions of precisely balanced neuronal microcircuits in an olfactory memory network. Cell Reports 44, 115330 (2025).

45 Blazing, R. M. & Franks, K. M. Odor coding in piriform cortex: mechanistic insights into distributed coding. Curr Opin Neurobiol 64, 96–102 (2020).

46 Stettler, D. D. & Axel, R. Representations of odor in the piriform cortex. Neuron 63, 854–864 (2009).

47 Poo, C. & Isaacson, J. S. A major role for intracortical circuits in the strength and tuning of odor-evoked excitation in olfactory cortex. Neuron 72, 41–48 (2011).

48 Rupprecht, P., Prendergast, A., Wyart, C. & Friedrich, R. W. Remote z-scanning with a macroscopic voice coil motor for fast 3D multiphoton laser scanning microscopy. Biomed. Opt. Express 7, 1656–1671 (2016).

49 Zhu, P., Fajardo, O., Shum, J., Zhang Schärer, Y.-P. & Friedrich, R. W. High-resolution optical control of spatiotemporal neuronal activity patterns in zebrafish using a digital micromirror device. Nat. Protoc. 7, 1410–1425 (2012).

50 Diaz-Verdugo, C., Sun, G. J., Fawcett, C. H., Zhu, P. & Fishman, M. C. Mating Suppresses Alarm Response in Zebrafish. Curr Biol 29, 2541–2546 e2543 (2019).

51 Deneve, S., Alemi, A. & Bourdoukan, R. The Brain as an Efficient and Robust Adaptive Learner. Neuron 94, 969–977 (2017).

52 Deneve, S. & Machens, C. K. Efficient codes and balanced networks. Nat Neurosci 19, 375–382 (2016).

53 Friedrich, R. W. & Laurent, G. Dynamic optimization of odor representations in the olfactory bulb by slow temporal patterning of mitral cell activity. Science 291, 889–894 (2001).

54 Friedrich, R. W., Habermann, C. J. & Laurent, G. Multiplexing using synchrony in the zebrafish olfactory bulb. Nat. Neurosci. 7, 862–871 (2004).

55 Niessing, J. & Friedrich, R. W. Olfactory pattern classification by discrete neuronal network states. Nature 465, 47–52 (2010).

56 Chen, T. W. et al. Ultrasensitive fluorescent proteins for imaging neuronal activity. Nature 499, 295–300 (2013).

57 Namekawa, I., Moenig, N. R. & Friedrich, R. W. Rapid olfactory discrimination learning in adult zebrafish. Exp Brain Res 236, 2959–2969 (2018).

58 Frank, T., Mönig, N. R., Satou, C., Higashijima, S.-i. & Friedrich, R. W. Associative conditioning remaps odor representations and modifies inhibition in a higher olfactory brain area. Nature Neurosci 22, 1844–1856 (2019).

59 Carr, W. E. S. in Sensory biology of aquatic animals (eds J. Atema, R. R. Fay, A. N. Popper, & W. N. Tavolga) 3–27 (Springer, 1988).

60 Friedrich, R. W. & Korsching, S. I. Combinatorial and chemotopic odorant coding in the zebrafish olfactory bulb visualized by optical imaging. Neuron 18, 737–752 (1997).

61 Lillicrap, T. P., Santoro, A., Marris, L., Akerman, C. J. & Hinton, G. Backpropagation and the brain. Nat Rev Neurosci 21, 335–346 (2020).

62 Poo, C. & Isaacson, J. S. Odor representations in olfactory cortex: “sparse” coding global inhibition, and oscillations. Neuron 62, 850–861 (2009).

63 Bundschuh, S. T., Zhu, P., Zhang Schärer, Y.-P. & Friedrich, R. W. Dopaminergic modulation of mitral cells and odor responses in the zebrafish olfactory bulb. J. Neurosci. 32, 6830–6840 (2012).

64 Zhu, P., Frank, T. & Friedrich, R. W. Equalization of odor representations by a network of electrically coupled inhibitory interneurons. Nature Neurosci. 16, 1678–1686 (2013).

65 Mathieson, W. B. & Maler, L. Morphological and electrophysiological properties of a novel in vitro preparation: the electrosensory lateral line lobe brain slice. J. Comp. Physiol. A 163, 489–506 (1988).

66 Yaksi, E. & Friedrich, R. W. Reconstruction of firing rate changes across neuronal populations by temporally deconvolved Ca2+ imaging. Nature Methods 3, 377–383 (2006).

67 Kitamura, K., Judkewitz, B., Kano, M., Denk, W. & Hausser, M. Targeted patch-clamp recordings and single-cell electroporation of unlabeled neurons in vivo. Nat Methods 5, 61–67 (2008).

68 Pologruto, T. A., Sabatini, B. L. & Svoboda, K. ScanImage: flexible software for operating laser scanning microscopes. BioMed. Eng. OnLine 2, 13 (2003).

69 Brette, R. & Gerstner, W. Adaptive exponential integrate-and-fire model as an effective description of neuronal activity. J Neurophysiol 94, 3637–3642 (2005).

70 McInnes, L., Healy, J. & Melville, J. UMAP: Uniform Manifold Approximation and Projection for Dimension Reduction. arXiv 10.48550/arXiv.1802.03426 (2020).

71 Schneider, S., Lee, J. H. & Mathis, M. W. Learnable latent embeddings for joint behavioural and neural analysis. Nature 617, 360–368 (2023).

## References

1 Chou C-N, Kim R, Arend LA, Yang Y-Y, Mensh BD, Shim WM, Perich MG, Chung S (2024) Neural Manifold Capacity Captures Representation Geometry, Correlations, and Task-Efficiency Across Species and Behaviors. bioRxiv:2024.2002.2026.582157; doi: 10.1101/2024.02.26.582157

